# Impaired synthesis of both telomere strands and adaptive TERT reduction in RTEL1 ATPase-dead cells

**DOI:** 10.64898/2026.07.23.740398

**Authors:** Guanhui Wu, Mingfeng Liu, Arthur J. Zaug, John L. Rinn, Thomas R. Cech

**Affiliations:** Department of Biochemistry, University of Colorado Boulder, Boulder, CO, 80303, USA; BioFrontiers Institute, University of Colorado Boulder, Boulder, CO, 80303, USA; Howard Hughes Medical Institute, University of Colorado Boulder, Boulder, CO, 80303, USA; Department of Molecular, Cellular and Developmental Biology, University of Colorado Boulder, Boulder, CO, 80309, USA

## Abstract

Regulator of Telomere Elongation Helicase 1 (RTEL1) plays a critical role in telomere replication by disassembling DNA secondary structures such as G-quadruplexes and telomeric loops (T-loops). However, its precise mechanism remains unclear. Previously, we generated HeLa cells homozygous for ATPase-dead *RTEL1*. Here, we found that these cells have a 50-77% reduction in the synthesis of both telomere strands, indicating that RTEL1 ATPase activity is important for both leading-strand and lagging-strand synthesis. Surprisingly, these cells also showed a substantial reduction in Telomerase Reverse Transcriptase (TERT) mRNA levels and an approximately 80% reduction in telomerase activity, without activating alternative lengthening of telomeres (ALT). Forced TERT overexpression suppressed proliferation and caused late S/G2 accumulation, implying that the naturally occurring reduction in TERT provides an adaptive mechanism for cell survival. Expressing the longest RTEL1 isoform at close to physiological levels failed to rescue growth defects, suggesting other isoforms are important for cell survival. These results reveal an unexpected interplay between RTEL1 and telomerase and show how cancer cells compensate for RTEL1 dysfunction by decreasing TERT expression.

**Graphical abstract:** 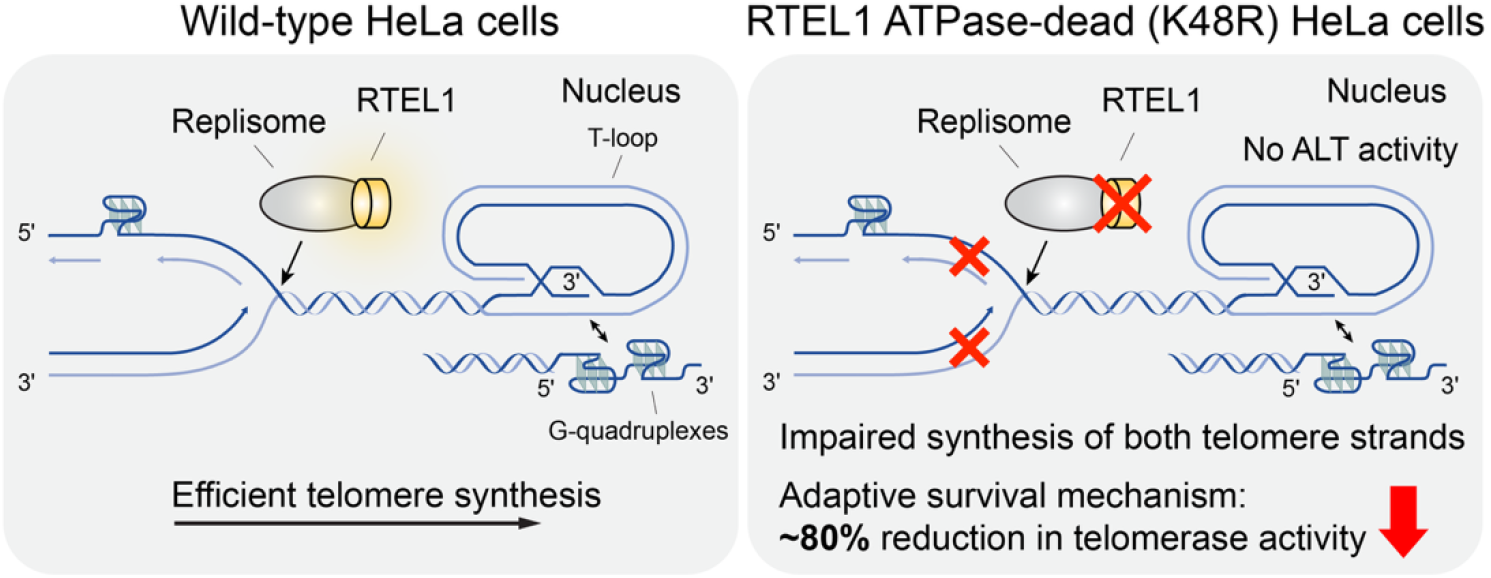

## Introduction

Telomeres are protective caps at the ends of chromosomes and are associated with cancer development and aging(1,2). In normal somatic cells, telomeric DNA gradually shortens with each division. Telomere shortening is associated with replicative senescence(3), although recent studies suggest that senescence is more complex(4). To achieve continuous proliferation, cells including cancer cells must maintain their telomeres. The common solution is reactivation of telomerase, which extends the G-rich telomeric overhangs(5). The CTC1-STN1-TEN1 (CST) complex then binds the single-stranded DNA (ssDNA) overhang and recruits Polα-primase to fill in the complementary C-strand(6).

Telomeric DNA is particularly difficult to replicate because of its repetitive and G-rich sequences. These sequences spontaneously form G-quadruplex structures in ssDNA, which act as physical barriers to the replication machinery(7–10). Moreover, telomeric DNA can adopt T-loop structures, in which the single-stranded telomere overhang invades the double-stranded telomeric DNA, creating another potential obstacle to fork progression(11).

RTEL1 is an iron-sulfur cluster-containing DEAH-box helicase essential for maintaining telomere and genome stability(12–14). The human chromosome region 20q13.33, which harbors the RTEL1 gene, is frequently amplified in cancers such as gastrointestinal tumors, hepatocellular carcinoma, and gliomas(15–17). In transgenic mouse models, conditional overexpression of Rtel1 leads to liver tumors in over 70% of mice, suggesting that RTEL1 can act as an oncogenic driver(18). Conversely, mutations in RTEL1 are linked to telomere biology disorders (TBDs), including dyskeratosis congenita (DKC), Hoyeraal-Hreidarsson syndrome (HHS), and pulmonary fibrosis(19–21). Knockout of Rtel1 in mouse cells causes severe telomere defects, including loss, fragility, and fusions, while a single amino acid change in the ATPase domain reduces the set-point for telomere length, highlighting the critical role of RTEL1 in telomere maintenance(12,22).

However, it remains unclear how RTEL1 maintains telomeres. Multiple mechanisms have been proposed: (i) unwinding G-quadruplex structures in the telomere overhang to facilitate telomerase-mediated extension(23,24); (ii) acting as a replisome-associated protein(25,26) that resolves T-loops and G-quadruplexes to support telomere replication(27); and (iii) regulating R-loops formed by telomeric repeat-containing RNA (TERRA) to prevent telomere fragility(28). Recent research also suggests that telomerase may contribute to telomere catastrophe in Rtel1-deficient mouse cells by blocking the restart of reversed replication forks(29).

To help dissect the role of RTEL1’s ATPase activity in human telomere maintenance, we previously generated RTEL1 ATPase-dead (K48R) HeLa cell lines (C17 and C19) using CRISPR genome editing(30). These cells grow much more slowly than parental HeLa cells and exhibit late S-phase arrest. They also display severe telomere defects, including shortened telomeres and elevated rates of chromosome end-to-end fusions. In addition, we established two cell lines (C1 and C15) expressing wild-type (WT) Halo-tagged RTEL1. Live-cell imaging revealed that RTEL1 forms more extensive interactions with telomeres during S-phase, supporting its role in telomere replication(30).

In this study, we initially aimed to investigate RTEL1 function using purified protein and characterize it using biochemical assays. However, due to the technical difficulty in purifying RTEL1 and validating its helicase activity, presumably due to its labile iron-sulfur cluster, we pursued an alternative approach by analyzing telomere defects in our ATPase-dead RTEL1 cell lines. We found that these cells have impaired synthesis of both telomere strands but do not show elevated T-circles or altered telomere overhangs.

Next, we performed RNA-seq using these ATPase-dead cells and found that immune and apoptotic pathways are highly upregulated, while many RNA-processing pathways and telomere organization are downregulated. Unexpectedly, TERT is among the most downregulated genes in the telomere organization pathway. Using a doxycycline (Dox)-inducible TERT overexpression system, we found that TERT downregulation may serve as a mechanism of survival in the absence of functional RTEL1. This TERT-RTEL1 interplay is further supported by the analysis of the Cancer Dependency Map (DepMap) database. Furthermore, expressing the longest WT RTEL1 isoform at close to physiological levels does not rescue the growth defects, suggesting that the other isoforms also contribute to survival. Overall, our study provides mechanistic insights into how RTEL1 maintains telomeres and uncovers an adaptive mechanism by which cancer cells evade RTEL1 dysfunction through TERT downregulation.

## Materials and methods

### Cell lines and cell cultures

Cell culture was performed as described previously(30). The parental HeLa-EM2-11ht cells(31) and genome-edited derivatives (WT C1, WT C15, ATPase-dead C17, and ATPase-dead C19)(30) were cultured in high-glucose Dulbecco’s Modified Eagle Medium (DMEM) supplemented with 10% fetal bovine serum (FBS), 1× GlutaMAX-I, and 1× Penicillin-Streptomycin at 37°C with 5% CO_2_.

### CsCl density gradient ultracentrifugation

The procedure follows a previous publication(32). Cells were incubated with 100 μM 5-bromo-2’-deoxyuridine (BrdU) for 18 hours. Genomic DNA was then extracted, and 15 μg was digested with HinfI and RsaI to release intact telomeric DNA. The DNA was combined with saturated CsCl to achieve a final density of 1.74 g/mL in about 5.9 mL. This density was verified by weighing 10 μL on an analytical balance. The samples were transferred to Quick-Seal Ultra-Clear tubes (cat# 344075, Beckman Coulter) and centrifuged at 45000 rpm using a VTi65.2 rotor for 18 hours at 25 °C. After centrifugation, the tubes were clamped vertically on a stand, and small holes were punched in the top and bottom to allow collection of fractions (approximately 1 drop per fraction). The densities of fractions #12, #24, #36, and #48 were measured by weighing 10 μL of each. Next, 20 μL of each fraction was denatured with 10 μL of 0.5 M NaOH and 1.5 M NaCl for 30 minutes at 37 °C, then neutralized with 60 μL of 20× SSC. All fractions were transferred onto a positively charged nylon membrane, UV-irradiated at 260 nm (1200 μJ × 100 energy) for crosslinking, and hybridized at 50 °C for 2 hours with gentle rotation. Tel-G probe ([TTAGGG]_4_) and PerfectHyb Plus Hybridization Buffer (Sigma-Aldrich) were used for the hybridization.

### Native and denatured in-gel hybridization

In-gel hybridization to assess telomeric G-strand tails was performed according to a previously established protocol(33). Genomic DNA was extracted and measured using the Qubit 1× dsDNA HS Assay Kit (Thermo Fisher Scientific). DNA (3 μg) was digested overnight with HinfI and RsaI in the presence or absence of Exo I. The digested DNA was separated on a 0.8% agarose gel containing ethidium bromide (EtBr) at 50 V for 18 hours. The gel was then dried at 30°C for 2 hours using a gel drier, with a piece of Whatman 3MM paper and a piece of nylon membrane placed underneath. The EtBr signal was captured with a Typhoon imager.

To prepare a probe with high radioactivity, 50 ng of the Tel-C oligo ([CCCTAA]_4_) was labeled with 5 μL of ^32^P-γ-ATP (6000 ci/mmol) and 1 μL of T4 Polynucleotide Kinase (New England Biolabs) in a 10 μL reaction at 37°C for 1 hour, and unincorporated nucleotides were removed with Micro Bio-Spin P-6 Gel Columns (cat# 7326221, Bio-Rad).

The dried gel was pre-hybridized with Church and Gilbert’s hybridization buffer at 43°C for 45 minutes. The labeled probe was then added directly to the hybridization tube and incubated overnight. The next day, the gel was washed twice with 4× SSC and once with 4× SSC + 0.1% SDS, each wash lasting 30 minutes. To minimize background signals, the gel was transferred to a large glass dish and washed with over 200 mL of water for 30 minutes at room temperature, with shaking. After washing, the gel was transferred to an exposure cassette, covered with plastic wrap, exposed to a phosphor screen overnight, and imaged with a Typhoon imager.

For the denatured condition to assess total telomeric DNA, the gel was incubated with 0.5 M NaOH and 1.5 M NaCl for 30 minutes, then neutralized with Tris-NaCl buffer (1 M Tris, pH 7.5, 1.5 M NaCl) for 45 minutes. Afterward, the gel was hybridized with the Tel-C probe as described above and imaged again using a Typhoon imager.

### T-circle assay

The T-circle assay was conducted according to a previous publication(34). In brief, 10 µg of genomic DNA was digested with HinfI and RsaI. The resulting DNA was then annealed in 200 µL of solution containing 1× TE, 50 mM NaCl, and 250 pmol of the Tel-C primer ([CCCTAA]_4_), which has thiophosphate linkages at its three terminal nucleotides. After ethanol precipitation, the DNA was resuspended in water. Approximately 20-25% of the DNA (equivalent to 2-2.5 µg of input DNA) was used in a 40 µL reaction mixture that included 7.5 U of phi29 DNA polymerase (New England Biolabs), 1× phi29 reaction buffer, 100 µg/mL recombinant albumin, and 370 µM dNTPs. The polymerase was added last after mixing all components. The mixture was incubated at 30°C for 12 hours in a thermocycler, then heat-inactivated at 65°C for 20 minutes. The extension products were ethanol-precipitated, resuspended in alkaline gel running buffer with 1× loading dye, and separated on a 0.6% alkaline agarose gel at 760-800 V×h. The next day, the gel was neutralized with Tris-NaCl buffer (1 M Tris, pH 7.5, 1.5 M NaCl) for 45 minutes, dried at 40°C for 1.25 hours using a gel dryer, and stained with ethidium bromide. After imaging, the gel was incubated in Church and Gilbert’s hybridization buffer for 45 minutes and hybridized overnight at 50 °C in a hybridization oven with a 5’-end ^32^P-labeled Tel-G probe ([TTAGGG] ).

The following morning, the gel was washed twice with 4× SSC and once with 4× SSC + 0.1% SDS, each for 30 minutes. To reduce background signals, the gel was transferred to a large glass dish and washed with over 200 mL of water for 30 minutes at room temperature, with shaking. Finally, the gel was placed in an exposure cassette, covered with plastic wrap, exposed to a phosphor screen overnight, and imaged using a Typhoon imager.

### Telomeric repeat amplification protocol (TRAP)

The TRAP assay was performed as previously described(35). All buffer solutions were prepared in-house using DEPC-treated water. One million cells were lysed in 400 µL of NP-40 lysis buffer supplemented with RNasin Plus Ribonuclease Inhibitor (cat# N2611, Promega). The appropriate volume of cell lysate was used in the TRAP reaction as indicated above the gel. Each 50 µL reaction contained 1× TRAP buffer, 20 U of Taq DNA polymerase, 50 µM dNTPs, 0.2 fM TSNT, 2 ng/µL ACX primer, 2 ng/µL NT primer, 2 ng/µL Cy5-TS primer (5’-Cy5-AATCCGTCGAGCAGAGTT-3’), and 20 µg/mL BSA. Reaction products were resolved on a 10% acrylamide gel and imaged on a Typhoon imager.

### C-circle assay

The C-circle assay was conducted as previously detailed(36). Unlike the T-circle assay, it does not require DNA digestion and an exogenous primer. In brief, a 20 µL reaction mixture included 30 ng of genomic DNA, 7.5 U of phi29 DNA polymerase (New England Biolabs), 1× phi29 reaction buffer (New England Biolabs), 0.1% Tween 20, 1 mM dNTPs, and 4 µg/mL BSA. The reaction was incubated at 30°C for 8 hours and then heat-inactivated at 70°C for 20 minutes. After transferring the products onto a positively charged nylon membrane, it was UV-irradiated at 260 nm (1200 μJ × 100 energy) and pre-hybridized in PerfectHyb Plus Hybridization Buffer (Sigma-Aldrich) for 30 minutes at 50°C. Subsequently, a 5’-end ^32^P-labeled C-circle probe (5’-CTAACCCTAACCCTAACC-3’) was added and incubated for 3 hours. The membrane was washed twice with 2× SSC and once with 2× SSC + 0.1% SDS, each wash lasting 15 minutes.

To strip the C-circle probe, the membrane was incubated with 50 mM NaOH at 45°C for 30 minutes, followed by a 15-minute wash in 0.1× SSC + 0.1% SDS. Finally, the membrane was re-hybridized with the Alu probe (5’-GTAATCCCAGCACTTTGG-3’) to confirm loading consistency.

### Co-immunoprecipitation

The co-immunoprecipitation (co-IP) was performed as previously described with minor modifications(37). At least 50 million HeLa-EM2-11ht cells were harvested and lysed in an ice-cold buffer containing 50 mM Tris-Cl (pH 7.5), 20% glycerol, 1 mM EDTA, 150 mM NaCl, 0.5% Triton X-100, 0.02% SDS, 1 mM DTT, and 1× complete protease inhibitor cocktail (Roche). After a 30-minute incubation at 4°C, 5 M NaCl was added to a final concentration of 400 mM. The mixture was incubated on ice for 5 minutes, then centrifuged at 14000 rpm for 10 minutes. The supernatant was collected for the IP. For each IP, 10 μg of anti-RTEL1 (cat# NBP2-22360, Novus Biologicals) or control antibody (cat# 12-370, Sigma-Aldrich) was added to the lysate, then incubated at 4°C for 2 hours. Subsequently, pre-blocked protein A and G agarose beads were added, and the mixture was incubated overnight at 4°C. The next day, the binding complexes were washed three times with HYPO-150 buffer, which contains 20 mM Na-HEPES (pH 8.0), 150 mM NaCl, 2 mM MgCl2, 0.2 mM EGTA, 1 mM DTT, 0.1% NP-40, and 10% glycerol. Proteins were eluted using 1× NuPAGE LDS sample buffer and analyzed by Western blot with the following antibodies: anti-PCNA (1:250; cat# 13110, Cell Signaling Technology) and anti-RTEL1 (1:100; cat# HPA078328, Sigma-Aldrich).

### Lentivirus packaging, infection, and genome editing

HEK293T cells were seeded in 10-cm dishes on day 1 and transfected on day 2 with 5.4 μg of transfer vector (see Supplementary Transfer Vector Sequences), 3.2 μg of pDML6 gag/pol vector, 1.8 μg of pRSV-Rev vector, and 1.8 μg of pVSV-G vector. Six hours after transfection, the medium was replaced with tetracycline- and antibiotic-free medium. On day 4, the supernatant containing the lentivirus was collected and filtered through a 0.45 μm filter. Subsequently, 0.5 mL of the filtered supernatant was mixed with 1.5 mL of non-virus medium and used to infect HeLa-EM2-11ht cells in 6-well plates. One day post-infection, the virus was removed, and selection was carried out using 0.2 mg/mL zeocin for two weeks. Stable cell lines were maintained in the presence of 0.1 mg/mL zeocin.

For CRISPR-mediated introduction of an ATPase-dead RTEL1 (K48R) mutant, we followed our established protocol(30). One day after transfection with donor and Cas9/sgRNA-encoding vectors, edited cells were treated with 2 μg/mL puromycin for 24 hours, then allowed to recover for 24 hours, and then plated in either a 6-well plate for colony formation assays or in T-75 flasks for genomic DNA extraction at specified time points. After initial selection, edited cells were cultured in the presence of 1 μg/mL puromycin unless otherwise stated.

### RNA-seq

Total RNA was isolated using the Direct-zol RNA Miniprep Plus Kits (Zymo Research). Next-generation sequencing of rRNA-depletion libraries was performed using GENEWIZ® NGS Services from Azenta Life Sciences, with an average sequencing depth of approximately 57 million paired-end reads per sample (range: 44-80 million per sample) and a mean uniquely mapped rate of 70.7% (range: 62.9-72.7%).

RNA samples were quantified using a Qubit 4.0 Fluorometer (Thermo Fisher Scientific), and RNA integrity was checked with 4200 TapeStation (Agilent Technologies). rRNA-depletion sequencing library was prepared using QIAGEN FastSelect rRNA HMR Kit (Qiagen). RNA sequencing library preparation used NEBNext Ultra II RNA Library Preparation Kit for Illumina (New England Biolabs), following the manufacturer’s recommendations. Briefly, enriched RNAs were fragmented for 15 minutes at 94°C. First-strand and second-strand cDNA were subsequently synthesized. cDNA fragments were end-repaired and adenylated at 3’ ends, and universal adapters were ligated to cDNA fragments, followed by index addition and library enrichment with limited cycle PCR. Sequencing libraries were validated using the Agilent TapeStation 4200 (Agilent Technologies) and quantified using a Qubit 4.0 Fluorometer (Thermo Fisher Scientific) as well as by quantitative PCR (KAPA Biosystems).

The sequencing libraries were multiplexed and clustered onto a flow cell on the Illumina NovaSeq instrument according to manufacturer’s instructions. The samples were sequenced using a 2×150 bp Paired End (PE) configuration. Image analysis and base calling were conducted by the NovaSeq Control Software (NCS). Raw sequence data (.bcl files) generated from Illumina NovaSeq were converted into fastq files and de-multiplexed using Illumina bcl2fastq 2.20 software. One mismatch was allowed for index sequence identification.

### RNA-seq analysis

Quality control, read mapping, and quantification were performed using nf-core/rnaseq v3.20.0(38). The cleaned reads were aligned to the GRCh38.p13 human reference genome and quantified using GENCODE v38 annotations(39,40). The alignment process was performed using STAR v5.1.0(41), and the quantification process was performed using Salmon v1.10.3(42).

Differential expression analysis was performed with DESeq2 (v1.42.1)(43) in R (v4.3.2). Genes were considered significantly differentially expressed (DEGs) if they had a Benjamini-Hochberg adjusted p-value (padj) < 0.05 and an absolute log2 fold change > 1, comparing ATPase-dead versus WT.

Pre-ranked GSEA(44) was performed on the genotype (ATPase-dead vs. WT) DESeq2 results using the fgsea package (v1.28.0)(45), with genes ranked by the DESeq2 Wald statistic. Gene sets corresponded to the Gene Ontology Biological Process collection (MSigDB (46,47) C5, BP subcategory) for Homo sapiens, obtained via msigdbr (v25.1.0, https://CRAN.R-project.org/package=msigdbr) and restricted to sets containing 15–500 genes. Enrichment scores and normalized enrichment scores (NES) were calculated with fgsea (minSize = 15, maxSize = 500, seed = 123 for reproducibility), and p-values were adjusted using the Benjamini-Hochberg method; pathways with padj < 0.05 were considered significantly enriched. Leading-edge genes contributing to enrichment of the GOBP_TELOMERE_ORGANIZATION gene set in the genotype (ATPase-dead vs. WT) contrast were extracted from the GSEA results. Normalized read counts and DESeq2 log2 fold changes for these genes were visualized to compare expression between WT and ATPase-dead genotypes. All plots were generated in R using ggplot2 v4.0.1(48).

### Immunostaining

In brief, cells were fixed with 4% formaldehyde for 15 minutes and then permeabilized with 1× PBS containing 0.3% Triton X-100 for an additional 15 minutes. The samples were then blocked with SuperBlock (PBS) Blocking Buffer (Thermo Fisher Scientific) supplemented with 0.1% Triton X-100 for 1 hour. Next, cells were incubated overnight at 4°C with the following primary antibodies: anti-TRF2 mouse monoclonal antibody (1:200; cat# NB100-56506, Novus Biologicals) and anti-HA rabbit monoclonal antibody (1:800; cat# 3724, Cell Signaling Technology). After incubation, cells were washed three times with 1× PBS containing 0.1% Triton X-100 for 5 minutes each, then incubated with secondary antibodies for 1 hour at room temperature: donkey anti-rabbit Alexa Fluor Plus 488 (1:200; cat# A-32790, Thermo Fisher Scientific) and donkey anti-mouse Alexa Fluor 647 (1:200; cat# A-31571, Thermo Fisher Scientific). Finally, cells were washed three times with 1× PBS containing 0.1% Triton X-100 for 10 minutes each. Slides were mounted with mounting medium containing DAPI (cat# H-1200, Vectashield) for nuclear staining.

## Results

### RTEL1 ATPase-dead mutant has impaired synthesis of both telomere strands

In attempts to characterize RTEL1 function biochemically, we purified FLAG-tagged RTEL1 from HEK293T cells following transient overexpression (Supplementary Figure 1). Unfortunately, WT FLAG-RTEL1 showed negligible ATPase activity, in contrast to UvrD helicase (positive control), and was similar to ATPase-dead RTEL1 (negative control), even when RTEL1 was prepared under anaerobic conditions. Moreover, these purified proteins lacked the characteristic UV-Vis absorption and brown color of iron-sulfur clusters(49), explaining the absence of ATPase activity.

We then explored the role of RTEL1 using an alternative approach. Spiking with labeled nucleosides or nucleoside analogs is a conventional strategy for detecting newly synthesized daughter telomeres(32). To assess the effect of RTEL1 ATPase-dead mutant on telomeric DNA synthesis, we treated cells with 5-bromo-2’-deoxyuridine (BrdU) and then performed CsCl density-gradient ultracentrifugation. This method distinguishes telomeres with newly synthesized G- and C-strands from unreplicated ones by density (Figure 1A, top diagram). Fractions were collected either by an automated system (Supplementary Figure 2) or manually (Figure 1A). These fractions were then hybridized with telomeric DNA probes on a dot blot.

**Figure 1.**
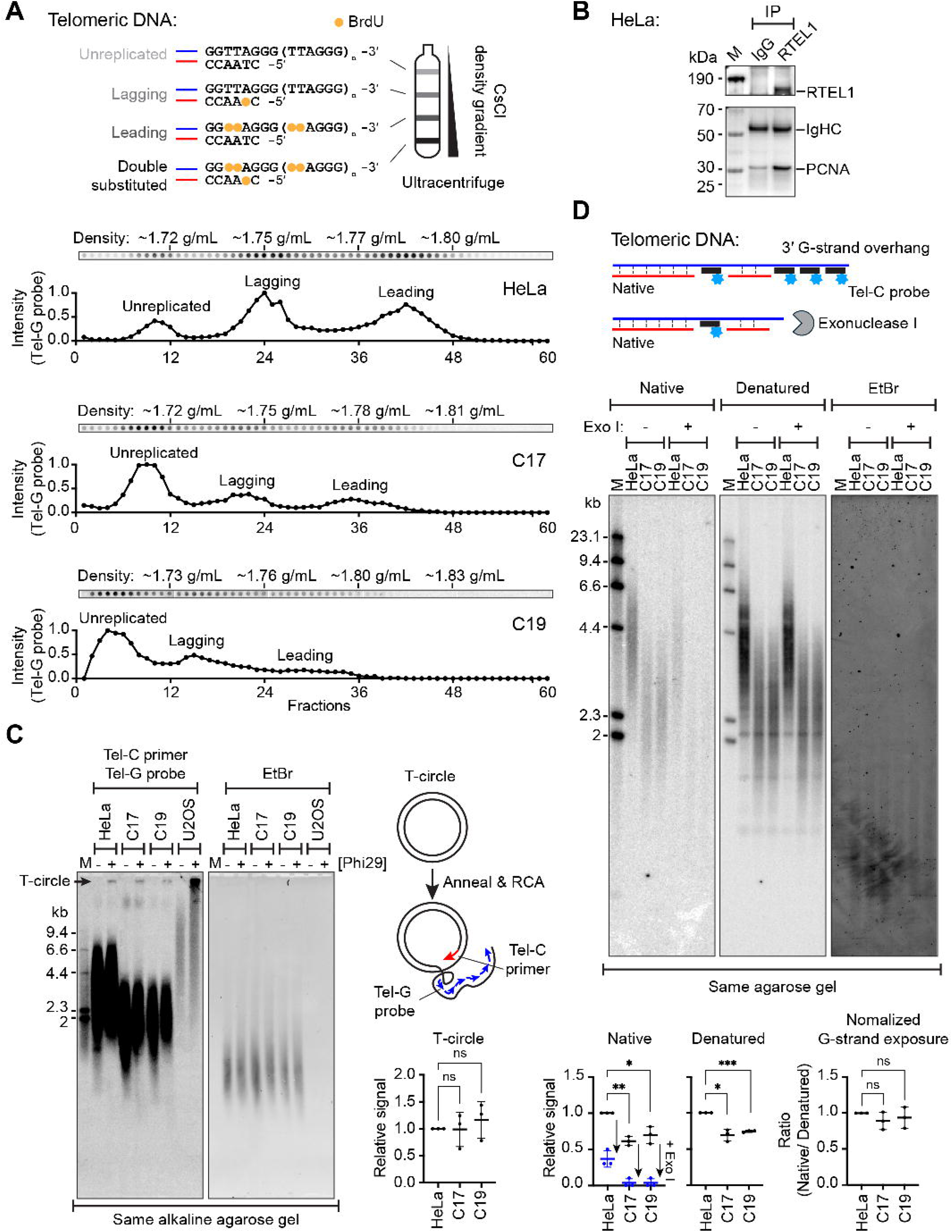
RTEL1 ATPase activity is required for the synthesis of both telomere strands. (**A**) CsCl density-gradient ultracentrifugation analysis of newly synthesized telomeres in parental HeLa and RTEL1 ATPase-dead (C17 and C19) cells. After 18 h of BrdU labeling, newly synthesized G- and C-rich telomeric strands incorporated different amounts of BrdU, resulting in distinct densities (top diagram). Following manual fractionation, each telomere species was quantified by denaturing dot-blot hybridization with a telomere-specific probe (Tel-G), which hybridizes to the C-rich strand. CsCl densities for fractions 12, 24, 36, and 48 are indicated above the dot blots. Quantification results are shown below each blot. Relative to HeLa, leading- and lagging-strand synthesis were respectively reduced by ∼63% and ∼62% in C17 and by ∼77% and ∼51% in C19. Additional validations using an automated fractionation system are provided in Supplementary Figure 2. (**B**) Co-immunoprecipitation (co-IP) analysis of RTEL1-PCNA interactions in parental HeLa cells. RTEL1 was immunoprecipitated using an anti-RTEL1 antibody, and PCNA was detected by Western blot. Normal rabbit IgG served as a mock control to assess non-specific binding. (**C**) T-circles in parental HeLa and ATPase-dead cells. T-circles were amplified by rolling circle amplification (RCA) using a Tel-C primer, which hybridizes to the telomeric G-rich strand. Products were separated on a 0.6% alkaline agarose gel and detected by in-gel hybridization with a ^32^P-end-labeled Tel-G probe. Ethidium bromide (EtBr) staining was used to assess loading. U2-OS (ALT-positive) cells served as a positive control; only 1% of the RCA product was loaded for U2-OS. Quantifications are shown next to the panel. Data are shown as mean ± SD, n = 3 independent replicates. Two additional independent replicates are provided in Supplementary Figure 3. ns, p > 0.1234; Fisher’s LSD test after one-way ANOVA. (**D**) In-gel hybridization analysis of telomeric G-strand ssDNA extension. Genomic DNA was separated on a 0.8% native agarose gel, and single-stranded G-rich telomeric DNA was detected by in-gel hybridization with a ^32^P-end-labeled Tel-C probe. The same gel was then denatured, neutralized, and rehybridized with the same probe to detect total telomeric DNA. EtBr staining was used to assess loading. Exo I treatment was used to confirm that the native signal originates from telomeric overhangs (top diagram). Quantifications are shown below the panel. Data are shown as mean ± SD, n = 3 independent replicates. *p < 0.0332, **p < 0.0021, ***p < 0.0002; Fisher’s LSD test after one-way ANOVA. Two additional independent replicates are provided in Supplementary Figure 4.

Results revealed that parental HeLa cells had high levels of newly synthesized telomeres, whereas RTEL1 ATPase-dead cells showed approximately 50-77% less BrdU incorporation, indicating defects in telomere synthesis on both strands.

Given that RTEL1 interacts with the replisome(25,26,50) to facilitate telomere replication, we performed co-immunoprecipitation (co-IP) experiments (Figure 1B). PCNA, a core replisome component, was enriched in RTEL1 pull-downs relative to IgG controls, consistent with prior observations, confirming the RTEL1-replisome association in HeLa cells.

In mouse cells, RTEL1 unwinds T-loops(27). Mutations lead to persistent T-loops that recruit SLX1/4 endonucleases, resulting in T-loop cleavage and increased T-circle formation. We therefore performed T-circle assays on DNA from our human RTEL1 mutant cells using an ALT cell line, U2-OS, as a positive control (Figure 1C and Supplementary Figure 3)(51). As expected, U2-OS showed strong T-circle signals even when only 1% of the rolling circle amplification (RCA) products were loaded in gels. In contrast, both parental HeLa and RTEL1 ATPase-dead cells exhibited similarly weak signals, suggesting that T-circles are largely suppressed in these cells, in contrast to the previous results in mouse cells(27). This finding prompted us to conduct additional T-circle experiments following the acute ATPase-dead RTEL1 knock-in, as described later.

Next, to test whether ATPase-dead RTEL1 causes strand-specific defects, we performed in-gel hybridization experiments (Figure 1D and Supplementary Figure 4). Under native conditions, the Tel-C probe binds single-stranded G-rich telomeric DNA, including both the overhang and the single-stranded G-rich region generated during telomere replication. RTEL1 ATPase-dead cells showed shorter telomeres and weaker native signals than parental HeLa. Under denatured conditions, signals reflect total telomere length. Normalizing native-to-denatured signals yielded similar ratios, suggesting comparable G-strand exposure. Exo I treatment, which removes overhangs, significantly reduced native signals but had little effect on denatured signals. The similar reduction across three cell lines suggests that ATPase-dead cells maintain telomere overhang lengths comparable to those of parental cells. Overall, these in-gel hybridization results align with the ultracentrifugation data (Figure 1A), confirming defects in the synthesis of both telomere strands.

Unexpected downregulation of TERT in RTEL1 ATPase-dead cells

To compare WT and ATPase-dead cells at the transcriptomic level, we performed RNA-seq. Principal component analysis clearly demonstrated that ATPase-dead cells have a distinct transcriptomic profile from WT cell lines (Figure 2A). Subsequent gene set enrichment analysis (GSEA) revealed that the most upregulated pathways in ATPase-dead cells involve immune activation and apoptotic signaling (Figure 2B), which are common responses to DNA repair defects(52,53). The most downregulated pathways were related to ribosome biogenesis and RNA processing (Figure 2B and Supplementary Table 1), possibly reflecting transcriptional disruption due to abnormal DNA replication, although a direct role for RTEL1 in RNA regulation cannot be excluded. Notably, telomere organization was among the most significantly downregulated pathways, and TERT was the second-most downregulated gene within this pathway (Figure 2B-C and Supplementary Figure 5A-B).

**Figure 2.**
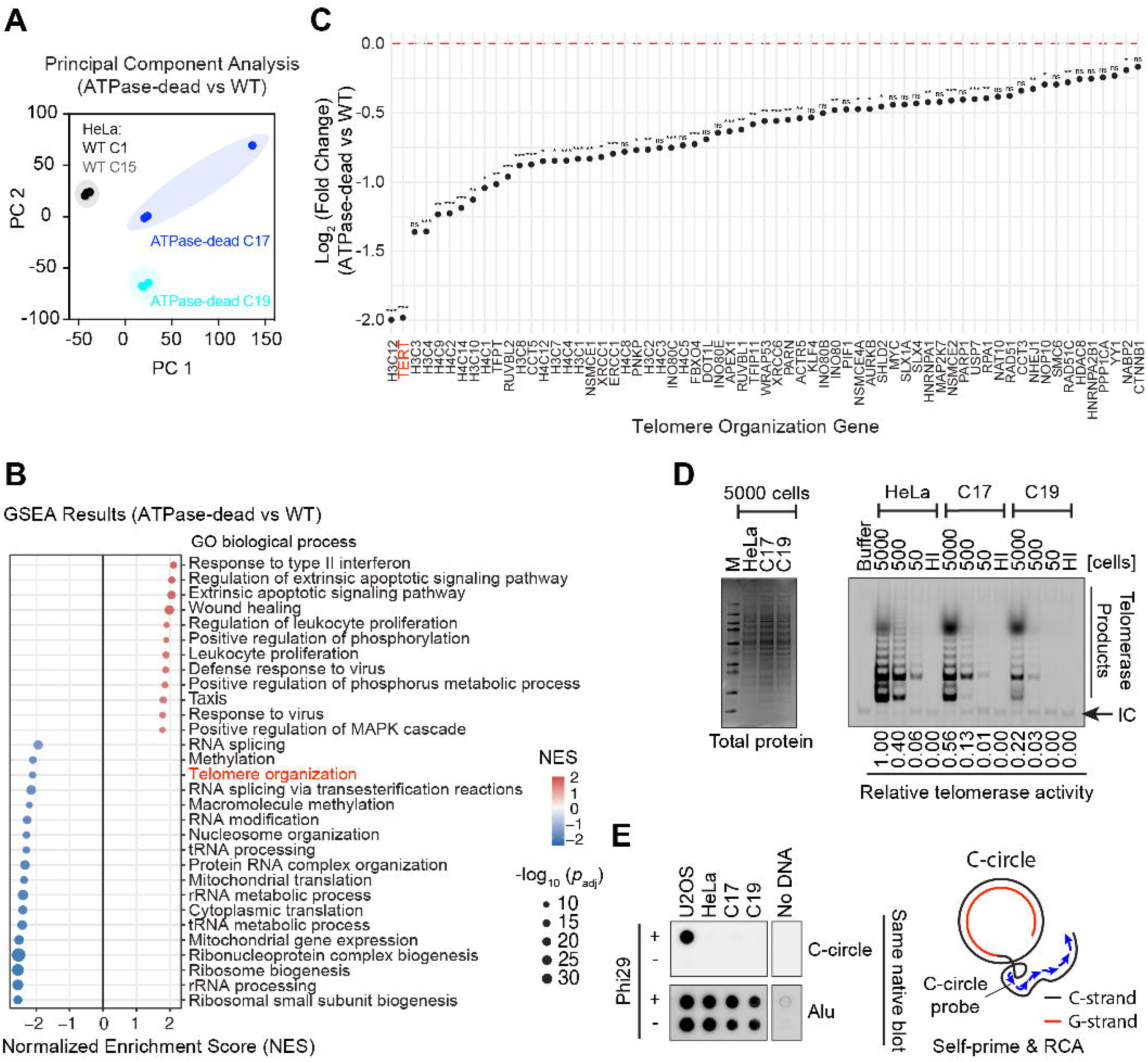
RTEL1 ATPase-dead cells show reduced TERT mRNA and decreased telomerase activity without ALT activation. (**A**) Principal component analysis of RNA-seq data from biological replicates of WT (C1, black; C15, gray) and ATPase-dead (C17, blue; C19, cyan) cells. Each group contains three independent replicates. (**B**) Gene set enrichment analysis (GSEA) showing the most upregulated and downregulated pathways in ATPase-dead cells. Normalized enrichment score (NES) is shown in red (upregulated) or blue (downregulated). Dot size represents -log_10_(adjusted p-value). GO, gene ontology. Complete GSEA results are provided in Supplementary Table 1; significant genes are provided in Supplementary Table 2. (**C**) Log_2_(fold change) of genes in the telomere organization pathway, comparing ATPase-dead cells to WT cells. ***p < 0.001, **p < 0.01, *p < 0.05. Raw data on TERT mRNA levels and a volcano plot comparing TERT to other genes are provided in Supplementary Figure 5A-B. (**D**) Telomerase activity measured by TRAP assay. Left: Coomassie blue-stained SDS-PAGE gel showing equal total protein amounts of cell lysates used in the TRAP assay. Right: TRAP products were resolved on a 10% native acrylamide gel, with the signal derived from Cy5-TS primer. Relative telomerase activity was determined by quantifying the intensity of the TRAP ladder. Quantitative comparison was performed at 500 cells per reaction, as HeLa signals were saturated at 5000 cells and signals for C17 and C19 approached the detection limit at 50 cells. Quantification at this cell number showed that telomerase activity was reduced by 65 ± 3% in C17 and 89 ± 3% in C19 relative to parental HeLa (mean ± range of two independent experiments; the relative activities shown below the gel are from the experiment displayed here). The number of cells used for each reaction is indicated above the gel. HI, heat-inactivated samples; IC, internal control for PCR amplification. An additional independent replicate is provided in Supplementary Figure 5C. (**E**) C-circle levels in parental HeLa, ATPase-dead, and U2-OS cells. C-circles were amplified by RCA using phi29 polymerase without an exogenous primer. Products were detected by dot blot with a ^32^P-end-labeled C-circle probe. U2-OS (ALT-positive) cells served as a positive control. No C-circle signal was detected in ATPase-dead cells, indicating the absence of ALT activity. Loading was verified by re-hybridization with an Alu probe after stripping.

We validated this finding using the TRAP assay(35), which showed an ∼80% reduction in telomerase activity in ATPase-dead RTEL1 cells (65 ± 3% and 89 ± 3% reduction in the two clones) relative to parental HeLa when 500 cells were tested (Figure 2D and Supplementary Figure 5C). To determine whether these cells activate the ALT pathway to compensate for reduced telomerase activity, we performed the C-circle assay (Figure 2E). Native blot results revealed a strong C-circle signal in U2-OS cells but undetectable in ATPase-dead cells, indicating that ALT is not activated in these cells. Thus, RTEL1 ATPase-dead cells maintain telomeres with significantly reduced telomerase activity, without activating ALT.

### Telomerase overexpression inhibits ATPase-dead RTEL1 cell growth

We reasoned that the low TERT and low telomerase activity in the ATPase-dead RTEL1 cells contributed to their telomere replication defects, which might then be ameliorated by restoring telomerase levels. To test this hypothesis, we initially attempted to overexpress telomerase directly in ATPase-dead cells. However, these poorly growing cells could not tolerate either transient transfection or lentiviral infection followed by antibiotic selection. We therefore adopted an alternative approach. First, we infected parental HeLa cells with lentiviruses carrying TERT or a control gene (EGFP). After Zeocin selection, Western blot confirmed Dox-inducible expression (Figure 3A). Next, we used CRISPR to introduce the ATPase-dead mutation into the endogenous RTEL1 locus, selecting correctly edited cells with puromycin (Figure 3B).

**Figure 3.**
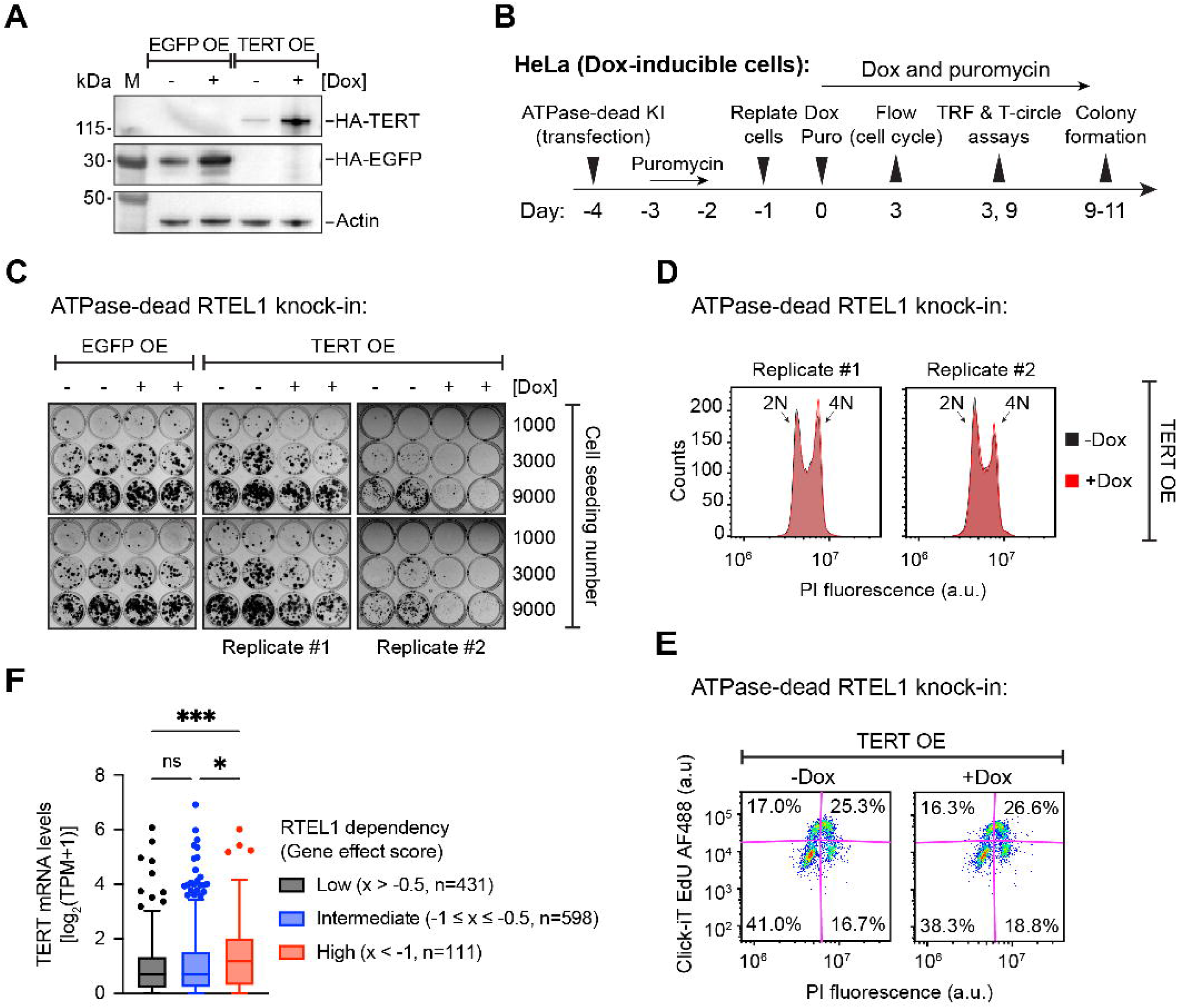
TERT overexpression inhibits RTEL1 ATPase-dead cell growth, and TERT expression influences RTEL1 dependency. (A) Western blot validation of Dox-inducible HA-tagged TERT and EGFP overexpression (OE). Leaky expression in the absence of Dox is due to basal activity of the Tet-inducible promoter. (**B**) Schematic representation of the experimental workflow for Figure 3C-E, Figure 4, Supplementary Figures 6B, 7C, and 12. (**C**) Colony formation assay. ATPase-dead cells were seeded at the indicated amounts and treated with Dox to induce EGFP or TERT OE. Colonies were stained 9-11 days later. TERT OE suppressed colony formation. Each column represents a biological replicate, and the rows within each column represent technical replicates. (**D**) DNA content analysis by propidium iodide (PI) staining after TERT OE. The 2N and 4N peaks are indicated. A subtle increase in the 4N population (3-4%) was consistently observed. (**E**) Cell cycle analysis combining PI staining and EdU labeling. Percentages of cells in G0/G1, early S, late S, and G2/M phases are shown. TERT OE leads to accumulation in late S and G2 phases. (**F**) Box-and-whisker plot of TERT mRNA levels across cell lines grouped by RTEL1 dependency. Cell lines from the Cancer Dependency Map (DepMap) were divided into three groups based on the RTEL1 gene effect score (x). RTEL1 dependency was defined as: low (> −0.5), intermediate (−0.5 to −1), and high (< −1). The number of cell lines (**n**) in each group is indicated. TERT mRNA levels are shown as log_2_(TPM+1). *p < 0.0332, ***p < 0.0002; Tukey’s multiple comparison test after one-way ANOVA.

Unexpectedly, colony formation assays showed that forced overexpression of TERT suppressed rather than rescued the growth of RTEL1 ATPase-dead cells (Figure 3C). EGFP had no effect in ATPase-dead cells, and TERT overexpression had no effect in parental HeLa cells (Figure 3C and Supplementary Figure 6A). Cell cycle analysis with propidium iodide (PI) staining revealed a subtle, consistent shift from the 2N to the 4N DNA peak upon TERT overexpression, with a 3-4% increase in the 4N population (Figure 3D). 5-ethynyl-2’-deoxyuridine (EdU) labeling indicated that this shift was primarily toward the late S and G2 phases (Figure 3E). These findings – which contradict our original hypothesis – instead suggest that high TERT levels inhibit ATPase-dead cell growth and that the naturally reduced telomerase activity observed in ATPase-dead cells may serve as an adaptive survival mechanism.

To determine if this growth inhibition required telomerase enzymatic activity, we overexpressed the catalytically inactive D712A TERT (Supplementary Figure 7A-B). In contrast to WT TERT, D712A TERT had no effect on colony formation in RTEL1 ATPase-dead cells or in parental HeLa cells (Supplementary Figure 7C), indicating that the growth inhibition depends on telomerase catalytic activity.

To test whether this phenomenon is generalizable across cancer cell panels, we analyzed data from the DepMap cancer vulnerability database. Cell lines were categorized into three groups based on RTEL1 dependency. Notably, the group with the highest RTEL1 dependency (i.e., lowest RTEL1 gene effect score) exhibited higher TERT mRNA levels (Figure 3F). Conversely, when cell lines were sorted by TERT mRNA levels (Supplementary Figure 8), the top decile with the highest TERT mRNA levels had a higher proportion of RTEL1-dependent cells than the remaining nine deciles combined. Collectively, these reveal a previously unappreciated association between TERT and RTEL1, suggesting that cells expressing very high TERT levels are likely sensitive to RTEL1 knockout.

Telomerase-mediated telomere elongation is not fully blocked by ATPase-dead RTEL1

To examine how telomerase and ATPase-dead RTEL1 affect telomere length, we knocked in ATPase-dead RTEL1 and treated cells with Dox to overexpress TERT (Figure 3B and Figure 4). Genomic DNA was isolated on days 3 and 9 after Dox treatment. As expected, overexpressing TERT in parental cells resulted in telomere elongation by day 9 (Figure 4A-B and Supplementary Figure 9). Knock-in of ATPase-dead RTEL1 caused telomere shortening (compare lanes 5 and 7), but TERT overexpression still led to telomere elongation (compare lanes 7 and 8). Notably, ATPase-dead RTEL1 partially inhibited telomerase-mediated telomere extension (Figure 4A-B). This is consistent with our earlier observation that telomerase remains active in stable ATPase-dead clones (Figure 2D). In addition, C-circle assays confirmed that ALT activity was undetectable upon acute RTEL1 inhibition, in contrast to U2-OS cells (Figure 4C). Since our previous studies revealed that Dox alone can have substantial effects on gene expression(54), we induced EGFP expression by adding Dox (Supplementary Figure 10). The similar telomere lengths in these EGFP control cells argue against this possibility.

**Figure 4.**
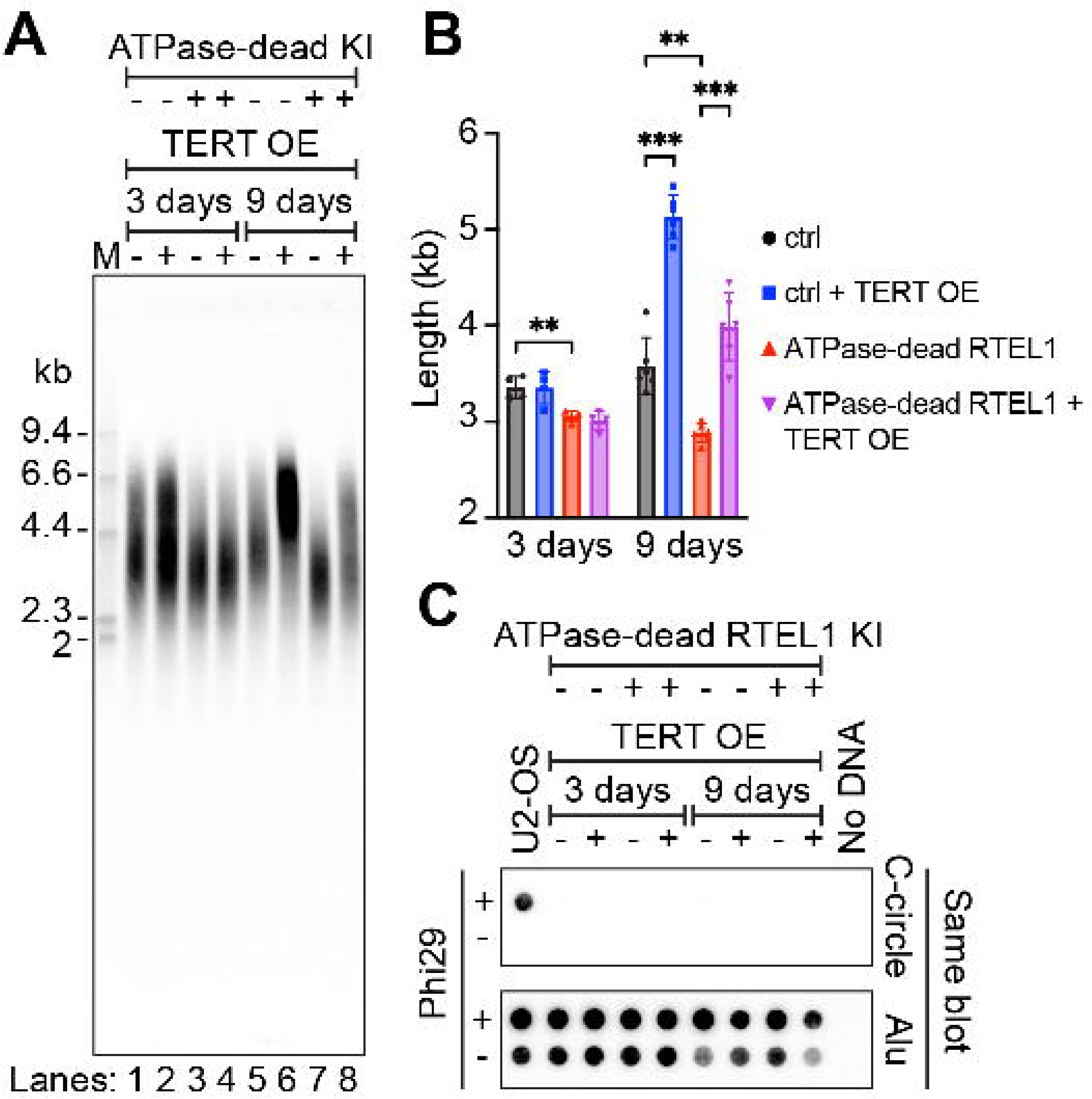
TERT overexpression extends telomeres in ATPase-dead cells. (A) Telomere restriction fragment (TRF) analysis of telomere length. Parental HeLa or ATPase-dead RTEL1 knock-in (KI) cells were treated with Dox to induce TERT overexpression (OE), as outlined in Figure 3B. Genomic DNA was isolated on day 3 and day 9 post-induction. Digested genomic DNA was separated on a 0.6% alkaline agarose gel and telomeric DNA was detected by in-gel hybridization with a P-end-labeled Tel-G probe. (**B**) Quantification of telomere length. Data are shown as mean ± SD, n ≥ 3 independent replicates. Additional telomere length and T-circle analyses are provided in Supplementary Figure 9. **p < 0.0021, ***p < 0.0002; Tukey’s multiple comparison test after two-way ANOVA. (**C**) Quantification of C-circle levels in the same genomic DNA samples. C-circles were amplified by RCA using phi29 polymerase without an exogenous primer. Products were detected by dot blot with a ^32^P-end-labeled C-circle probe. U2-OS (ALT-positive) cells served as a positive control. No C-circle signal was detected in ATPase-dead cells, confirming the absence of ALT activity. Loading was verified by re-hybridization with an Alu probe after stripping.

In our stable ATPase-dead clones (C17 and C19), T-circle levels showed no significant difference compared to parental cells (Figure 1C). We also examined T-circle levels after acute RTEL1 inhibition (Supplementary Figure 9). Although no consistent change was observed across time points or cell cultures, in one of the experimental replicates, we detected a transient increase in T-circles on day 3 (Supplementary Figure 9, cell culture #2). This change was reproducible in three independent rolling circle amplification (RCA) reactions using the same genomic DNA samples. By day 9, this difference had disappeared, while the baseline level in all groups was somewhat elevated compared to day 3. Given that the transient increase was not consistently reproduced at other time points or in other independent cell cultures (Supplementary Figure 9), we conclude that T-circle levels are dynamic and context-dependent in this system.

Overall, these results demonstrate that telomerase retains the ability to elongate telomeres even in the presence of ATPase-dead RTEL1, consistent with the absence of ALT activity in these cells.

### Failure to rescue ATPase-dead RTEL1 cells by expressing the longest RTEL1 isoform

RTEL1 mutations are associated with TBDs, and there is currently no curative treatment. We therefore sought to explore strategies to rescue the phenotype of ATPase-dead cells. The RTEL1 gene contains 35 exons and produces two primary isoforms [1300 and 1219 amino acids (aa)](55); the 1300 aa isoform has been shown to be the primary one for telomere maintenance(24,56). Additional isoforms may exist according to NCBI, UniProt, and Ensembl databases. Previously, we observed two well-separated RTEL1 bands in Western blot analysis of HeLa cell lysates(30), consistent with the presence of multiple isoforms.

Given the role of the 1300 aa RTEL1 isoform in telomere maintenance(24,56), we generated a Dox-inducible cell line expressing this isoform. Western blot analysis revealed a clear dose-dependent induction of RTEL1 expression upon Dox addition, with a minimal effective concentration of 5-10 ng/mL (Figure 5A). Consistent with this, immunostaining showed that 10 ng/mL Dox induced HA-tagged RTEL1 expression in most cells, and the protein was correctly localized in the cell nuclei (Figure 5B). To assess whether HA-tagged RTEL1 was recruited to telomeres, we simultaneously captured HA-RTEL1 and TRF2 by laser-scanning confocal microscopy (Figure 5C and Supplementary Figure 11), as TRF2 is important for RTEL1 recruitment to telomeres(56). Multiple HA-RTEL1 foci colocalized with TRF2 foci, indicating that HA-RTEL1 can still be recruited to telomeres.

**Figure 5.**
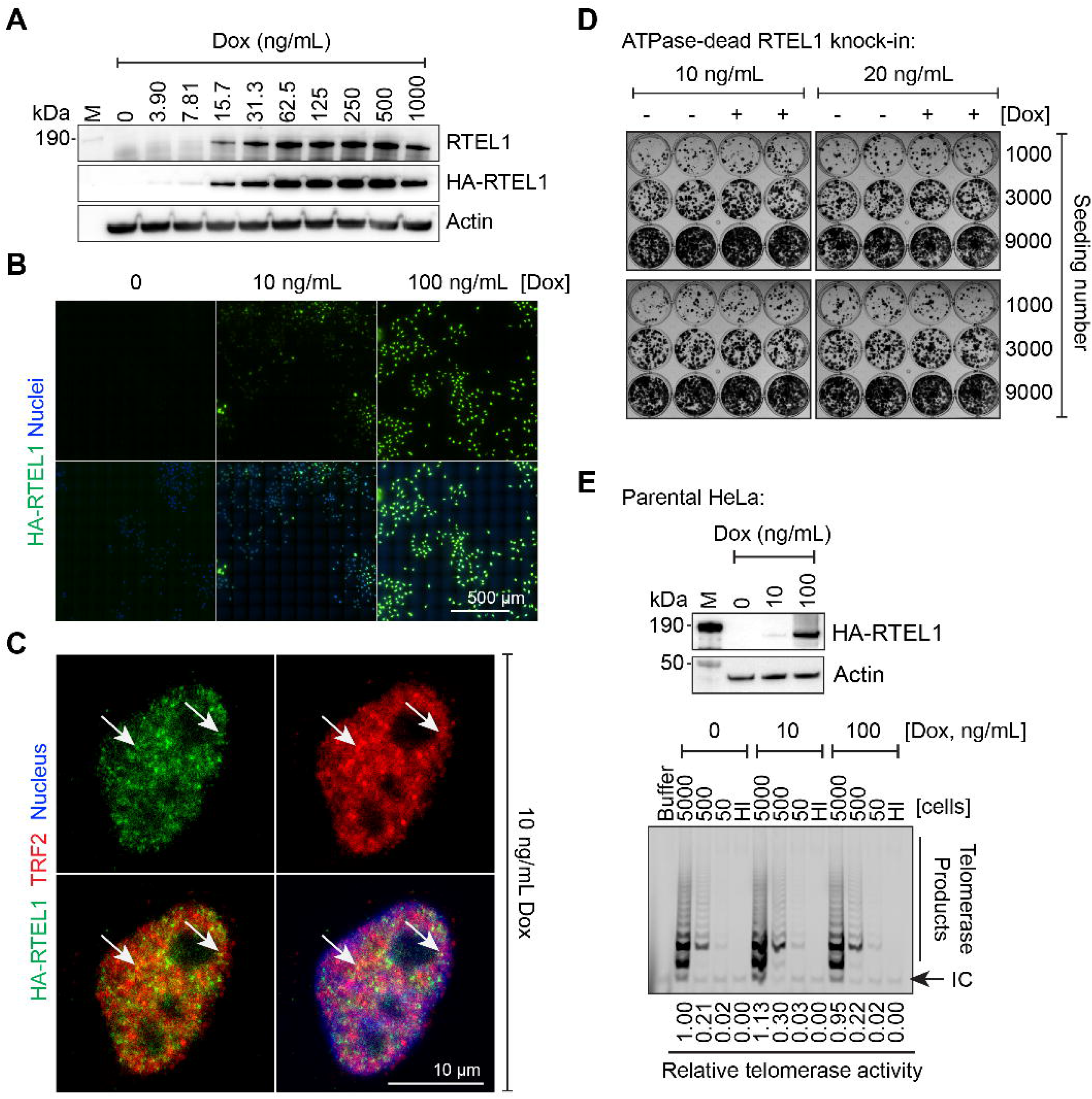
Expression of the longest RTEL1 isoform does not rescue ATPase-dead cell growth. (A) Western blot analysis of Dox-inducible HA-tagged WT RTEL1 expression. Dose-responsive RTEL1 expression was measured after cells were treated with different Dox concentrations for 48 hours. Due to its low abundance, endogenous RTEL1 was undetectable in Western blots when using cell lysates directly and required enrichment by IP, as shown in Figure 1B and previously reported(30). (**B**) Immunostaining analysis of HA-RTEL1 expression and subcellular localization. HA-RTEL1 was immunostained after cells were treated with 10 or 100 ng/mL Dox for 48 hours. HA-RTEL1 was readily detected in most cells treated with 10 ng/mL Dox. Images were captured and assembled from 100 images using a Nikon ECLIPSE Ti2 inverted widefield epi-fluorescence microscope equipped with a 100X NA 1.45 objective in scanning wizard mode. Green, HA-RTEL1; Blue, nuclei (DAPI). (**C**) Immunostaining analysis for the colocalization of HA-RTEL1 and TRF2. HA-RTEL1 and TRF2 were immunostained after cells were treated with 10 ng/mL Dox for 48 hours. Images were captured on a Nikon AXR laser scanning confocal microscope equipped with a 100X NA 1.45 objective. Green, HA-RTEL1; Red, TRF2; Blue, nucleus (DAPI). Arrows indicate examples of colocalized foci. Uncropped images are provided in Supplementary Figure 11. (**D**) Colony formation assay. ATPase-dead cells were seeded at the indicated amounts and treated with 10 or 20 ng/mL Dox to induce WT RTEL1 expression. Colonies were stained 9 days later. The overall workflow is the same as for TERT induction, as shown in Figure 3B. Each column represents a biological replicate, and the rows within each column represent technical replicates. (**E**) Telomerase activity measured by TRAP assay after RTEL1 induction in parental HeLa cells. Top: Western blot showing induction of RTEL1 expression in the cell lysates used for the TRAP assay. Bottom: TRAP products were resolved on a 10% native acrylamide gel, with the signal derived from Cy5-TS primer. Relative telomerase activity was determined by quantifying the intensity of the TRAP ladder. The number of cells used for each reaction is indicated above the gel. HI, heat-inactivated samples; IC, internal control for PCR amplification.

To test whether induction of WT RTEL1 could rescue the cells, we treated cells with 10 or 20 ng/mL Dox, as RTEL1 levels in this range are closer to the physiological level than at higher Dox concentrations. Colony formation results revealed that this was not sufficient to promote colony formation in the presence of ATPase-dead RTEL1 (Figure 5D), nor in the parental cells (Supplementary Figure 6B). To further determine whether the failure was caused by impaired telomere elongation, we measured telomere lengths after inducing expression of the RTEL1 1300 aa isoform (Supplementary Figure 12). The results showed that this isoform can facilitate telomere elongation regardless of the presence of ATPase-dead RTEL1. Our CRISPR knock-in introduced the ATPase-dead 1300 aa isoform into the endogenous RTEL1 loci while disrupting the other isoforms(30). The failure to rescue growth therefore suggests that other isoforms also contribute to cancer cell survival.

Overall, our results indicate that expressing the longest WT RTEL1 isoform cannot rescue the growth defects caused by the ATPase-dead mutation in the RTEL1 gene. Although TERT overexpression can elongate telomeres in the presence of ATPase-dead RTEL1 (Figure 4), it fails to promote cell survival and instead exacerbates cell cycle defects (Figure 3). These findings highlight the challenge of overcoming the ATPase-dead RTEL1 mutation by simply elongating telomere length or expressing the longest isoform. Further exploration is still needed to determine the functions of other isoforms of RTEL1 in cancer cell survival.

### The longest RTEL1 isoform does not regulate TERT expression

Because both TERT mRNA levels and activity were reduced in ATPase-dead RTEL1 cells, we asked whether RTEL1 regulates TERT expression. The TRAP assay uses an artificial, non-G-quadruplex-forming primer (TS primer) and therefore reports the levels of active telomerase rather than telomere extension by telomerase in cells. After overexpressing RTEL1 in parental HeLa cells, telomerase activity was similar across conditions (Figure 5E), indicating that the longest RTEL1 isoform does not directly regulate TERT expression.

## Discussion

In this study, we demonstrate that RTEL1 ATPase activity is essential for synthesizing both telomere strands, as evidenced by a 50-77% decrease in BrdU incorporation in newly synthesized telomeres in ATPase-dead cells. Unexpectedly, these cells show significantly lower TERT mRNA levels and a 65-89% reduction in telomerase activity, without activating ALT. Functional experiments reveal that forced overexpression of TERT in ATPase-dead cells suppresses cell growth and induces accumulation in late S/G2 phases, indicating that the natural reduction in telomerase activity serves as an adaptive mechanism for cell survival. Moreover, expressing the longest RTEL1 isoform at close to physiological levels failed to rescue cell proliferation, suggesting other isoforms also contribute to survival. Collectively, our results demonstrate the importance of RTEL1 in replicating both telomere strands, reveal unexpected crosstalk between RTEL1 and telomerase, and uncover how cancer cells can mitigate the harmful effects of catalytically inactive RTEL1 helicase by downregulating TERT.

The defects in synthesis on both telomere strands in ATPase-dead cells are likely due to impaired resolution of DNA secondary structures. Our previous single-molecule live-cell imaging studies revealed that ATPase-dead RTEL1 is still recruited to telomeres(30). This observation indicates that physical association with telomeres is not sufficient for its function. Rather, the resolution of DNA secondary structures, including G-quadruplexes and T-loops(27,57), by its ATPase-dependent helicase activity appears to be critical for promoting replication fork progression through difficult-to-replicate telomeric regions.

The most unexpected finding is the significant reduction of TERT mRNA in ATPase-dead cells. RNA-seq results revealed that TERT is among the most downregulated genes in the telomere organization pathway. Because TERT mRNA is so lowly expressed even in wild-type HeLa – with most cells having 0-2 mRNA molecules(58) – the observed four-fold reduction in TERT mRNA levels in the population of RTEL1 mutant cells likely means that many individual cells have zero mRNA molecules at any given time. TRAP experiments indicated that this reduction in TERT led to a reduction in cellular telomerase activity, which likely contributed to the shorter telomeres observed by in-gel hybridization. The functional significance of TERT reduction was assessed using Dox-inducible TERT overexpression. Although TERT overexpression was sufficient to elongate telomeres in ATPase-dead cells, it impaired colony formation and caused additional cell-cycle defects beyond those already caused by ATPase-dead RTEL1. These observations indicate that the naturally occurring reduction of TERT in ATPase-dead RTEL1 cells is beneficial to their survival. Therefore, we propose that low telomerase activity in ATPase-dead cells – either through active downregulation or selective enrichment of low telomerase cells during prolonged cell culture – limits telomere elongation and avoids replication stress on telomeres. This adaptive reduction is consistent with previous observations in mouse cells, where high telomerase levels stabilize reversed replication forks and lead to telomere catastrophe(29). Notably, overexpression of catalytically inactive D712A TERT did not impair colony formation, indicating that the inhibition requires TERT catalytic activity. How elevated telomerase activity impairs growth in the absence of RTEL1 function remains to be determined. DepMap data show that cell lines with high RTEL1 dependency are associated with elevated TERT mRNA levels, further supporting the model that TERT expression sensitizes cells to RTEL1 inhibition.

Notably, unlike the high abundance of T-circles observed in Rtel1-deficient mouse cells(25,27,56), stable RTEL1 ATPase-dead human HeLa cells show only baseline levels of T-circles. Such low levels of T-circles have also been reported in multiple studies using RTEL1-mutant human patient samples(19,24,59,60). The discrepancy between humans and mice likely reflects species differences, particularly that mouse telomeres are substantially longer than human telomeres. In one cell culture, we observed a transient increase in T-circle levels in HeLa cells by day 3, but the difference from the control group disappeared by day 9. Prolonged culture may have led to adaptation, potentially through downregulation of TERT and possibly other unknown factors, as suggested by the substantial transcriptomic differences between WT and ATPase-dead cells. If confirmed more generally, the transient T-circle increase observed in this single experiment could offer an alternative explanation for the discrepancy in T-circle levels between mouse and human RTEL1-deficient cells. However, given the variability of this observation, further studies are still needed. From a clinical perspective, using T-circles as a biomarker for RTEL1 inhibition or dysfunction may require careful consideration of disease stage and cellular adaptation status.

Expression of the longest RTEL1 isoform at near-physiological to modestly elevated levels failed to rescue growth defects in cells with an ATPase-dead mutation, even though the exogenous protein was correctly localized to the cell nuclei, colocalized with TRF2 at telomeres, and facilitated telomere elongation. Because RTEL1 has multiple isoforms, other isoforms may have distinct functions and also be important for cell survival. Together with the enhanced cell defects upon TERT overexpression in ATPase-dead cells, these findings underscore the complexity of overcoming RTEL1 dysfunction. Potential therapeutic strategies may need to target the underlying replication stress rather than simply restoring telomere length or the levels of the longest RTEL1 isoform.

Several limitations of this study should be acknowledged. First, all experiments were conducted using the HeLa cell line, which is HPV-transformed. It remains uncertain whether adaptive TERT reduction occurs in other human cancer cell lines or stem cells. Second, we were unable to purify RTEL1 with ATPase activity and an Fe-S cluster. Obtaining a functional protein will be essential for future biochemical studies and the development of RTEL1-targeted therapies. Third, the molecular mechanism underlying TERT downregulation remains unknown. Our transcriptomic data show upregulation of immune and apoptotic pathways and downregulation of many RNA-related pathways. Whether TERT downregulation results from these pathways or from ATPase-dead RTEL1 itself requires further investigation. For instance, identifying the chromatin-binding sites of endogenous RTEL1 is of interest but technically challenging due to its low abundance and the presence of multiple isoforms in HeLa cells(24,30). Additionally, it remains unclear whether RTEL1 can directly regulate RNA processing, given the extensive downregulation of RNA-related pathways. Lastly, while our study identifies TERT as a key factor influencing RTEL1 dependency, it is likely not the sole determinant. Future systematic analyses integrating the DepMap and TCGA databases will be essential for understanding the contributions of tumor lineage and other molecular features.

In conclusion, our study demonstrates that RTEL1 ATPase activity is essential for proper synthesis of both telomere strands. Additionally, sustained RTEL1 inhibition leads to adaptive resistance, which is associated with TERT downregulation. The longest WT RTEL1 isoform failed to rescue ATPase-dead mutations, suggesting other isoforms also play important roles. This insight should be considered when developing therapeutic strategies for patients with TBDs.

Since many cancers depend on RTEL1, understanding how cancer cells adapt to RTEL1 impairments could improve patient stratification. For instance, combining RTEL1 inhibition with strategies targeting TERT-high tumors might offer a promising therapeutic approach.

## Acknowledgements

We thank Annette H. Erbse for her valuable suggestions regarding CsCl ultracentrifugation.

Ultracentrifugation was carried out using a Beckman L8-70 ultracentrifuge in the Shared Instruments Pool (RRID: SCR_018986) at the Department of Biochemistry, University of Colorado Boulder. Cell culture and flow cytometry were carried out at the Biochemistry Cell Culture Facility (RRID: SCR_018988) and the Flow Cytometry Shared Facility (RRID: SCR_019309), which are supported by NIH grant S10ODO2160. The imaging work was performed at the BioFrontiers Institute’s Advanced Light Microscopy Core (RRID: SCR_018302).

## Author contributions

Conceptualization: G.W., T.R.C.; Investigation: all authors; Project Administration: T.R.C.; Supervision: T.R.C., J.L.R.; Funding acquisition: T.R.C., J.L.R.; Visualization: all authors; Writing – original draft: G.W., T.R.C.; Writing – review & editing: all authors.

## Supplementary data

Supplementary Data are available at NAR online.

## Data availability

The uncropped images for Figures 1-5 are provided in Supplementary Figure 13. The sequencing data generated in this study have been deposited in the NCBI Gene Expression Omnibus under accession number GSE338949. All code to reproduce the RNA-seq analyses is available on Zenodo: https://doi.org/10.5281/zenodo.22004058.

