## Supplementary Data for "Impaired synthesis of both telomere strands and adaptive TERT reduction in RTEL1 ATPase-dead cells"

**This file includes:**

Supplementary Figures 1 to 13

Supplementary Transfer Vector Sequences

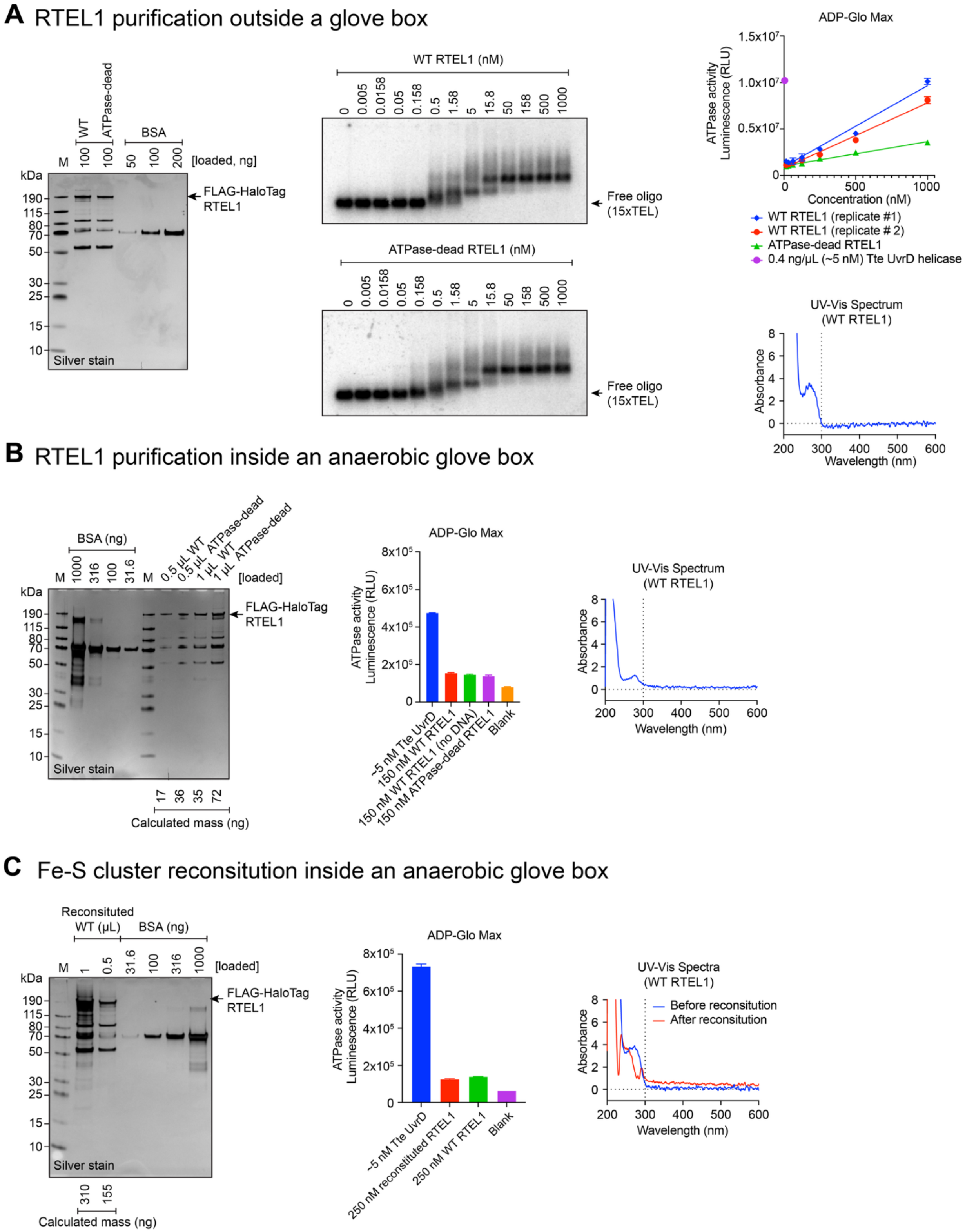

A RTEL1 purification outside a glove box
M
WT
ATPase-dead
BSA
100
100
50
100
200
[loaded, ng]
kDa
190
115
80
70
50
30
25
15
10
Silver stain
FLAG-HaloTag RTEL1
WT RTEL1 (nM)
0 0.005 0.0158 0.05 0.158 0.5 1.58 5 15.8 50 158 500 1000
Free oligo (15xTEL)
ATPase-dead RTEL1 (nM)
0 0.005 0.0158 0.05 0.158 0.5 1.58 5 15.8 50 158 500 1000
Free oligo (15xTEL)
ADP-Glo Max
ATPase activity Luminescence (RLU)
1.5x10^7
1.0x10^7
0.5x10^7
0
0
500
1000
Concentration (nM)
WT RTEL1 (replicate #1)
WT RTEL1 (replicate # 2)
ATPase-dead RTEL1
0.4 ng/μL (~5 nM) Tte UvrD helicase
UV-Vis Spectrum (WT RTEL1)
Absorbance
8
6
4
2
0
200 300 400 500 600
Wavelength (nm)
B RTEL1 purification inside an anaerobic glove box
BSA (ng)
M
1000
316
100
31.6
M
0.5 μL WT
0.5 μL ATPase-dead
1 μL WT
1 μL ATPase-dead
[loaded]
kDa
190
115
80
70
50
30
25
15
10
Silver stain
FLAG-HaloTag RTEL1
17 36 35 72
Calculated mass (ng)
ADP-Glo Max
ATPase activity Luminescence (RLU)
8x10^5
6x10^5
4x10^5
2x10^5
0
~5 nM Tte UvrD
150 nM WT RTEL1
150 nM WT RTEL1 (no DNA)
150 nM ATPase-dead RTEL1
Blank
UV-Vis Spectrum (WT RTEL1)
Absorbance
8
6
4
2
0
200 300 400 500 600
Wavelength (nm)
C Fe-S cluster reconsitution inside an anaerobic glove box
Reconsituted WT (μL)
BSA (ng)
M
1
0.5
31.6
100
316
1000
[loaded]
kDa
190
115
80
70
50
30
25
15
10
Silver stain
FLAG-HaloTag RTEL1
310 155
Calculated mass (ng)
ADP-Glo Max
ATPase activity Luminescence (RLU)
8x10^5
6x10^5
4x10^5
2x10^5
0
~5 nM Tte UvrD
250 nM reconstituted RTEL1
250 nM WT RTEL1
Blank
UV-Vis Spectra (WT RTEL1)
Before reconstitution
After reconstitution
Absorbance
8
6
4
2
0
200 300 400 500 600
Wavelength (nm)

**Supplementary Figure 1. Purified RTEL1 lacks ATPase activity and Fe-S cluster absorbance.**

**(A)** Characterization of WT and ATPase-dead FLAG-HaloTag-RTEL1 purified outside the glove box. Left: RTEL1 protein concentrations were estimated by quantifying band intensities of RTEL1 and BSA standards after silver staining. Middle: EMSA showed that WT and ATPase-dead RTEL1 bind the 15xTEL oligo (TTAGGG, 15 repeats). Upper right: WT protein showed ATPase activity slightly higher than that of ATPase-dead RTEL1 but much lower than that of Tte UvrD helicase (New England Biolabs). ATPase activity was measured at 37 °C for 40 min in reactions containing 20 mM Tris (pH 7.5), 0.56 mM EDTA, 20 mM $MgCl_2$, 3 mM KCl, 133 mM NaCl, 2 mM DTT, 100 ng/µL ΦX174 Virion ssDNA (New England Biolabs), and 5 mM ATP using the ADP-Glo Max Assay kit (Promega). Lower right: UV-Vis spectrum did not show characteristic absorbance for an Fe-S cluster.

**(B)** Characterization of WT and ATPase-dead FLAG-HaloTag-RTEL1 purified inside the glovebox. Left: RTEL1 protein concentrations were estimated by quantifying band intensities of RTEL1 and BSA standards after silver staining. Middle: WT RTEL1 had ATPase activity similar to that of ATPase-dead RTEL1, both of which were much lower than that of Tte UvrD helicase. Right: UV-Vis spectrum did not show characteristic absorbance for an Fe-S cluster. All buffer solutions for protein preparation and the ADP-Glo assay were degassed, and dissolved oxygen was removed by leaving the bottles uncapped inside the glovebox overnight. The solutions were then stored inside the glovebox.

**(C)** Characterization of WT FLAG-HaloTag-RTEL1 following attempted Fe-S cluster reconstitution inside a glove box. Reconstitution protocol used a reaction mixture containing 10 mM Tris (pH 8), 5 µM WT RTEL1, 400 µM $Na_2S$, 400 µM ferrous ammonium sulfate, and 5 mM DTT, incubated for 30 min at room temperature in the glovebox. The reconstituted protein was desalted using a P6 column (Bio-Rad) to remove excess salt. Left: After Fe-S cluster reconstitution, RTEL1 protein concentrations were estimated by quantifying band intensities of RTEL1 and BSA standards following silver staining. Middle: ATPase activity of WT RTEL1 remained similar before and after Fe-S cluster reconstitution. Right: Although some changes were observed in the UV-Vis spectra, they did not indicate the presence of an Fe-S cluster, which typically shows a peak near 400 nm.

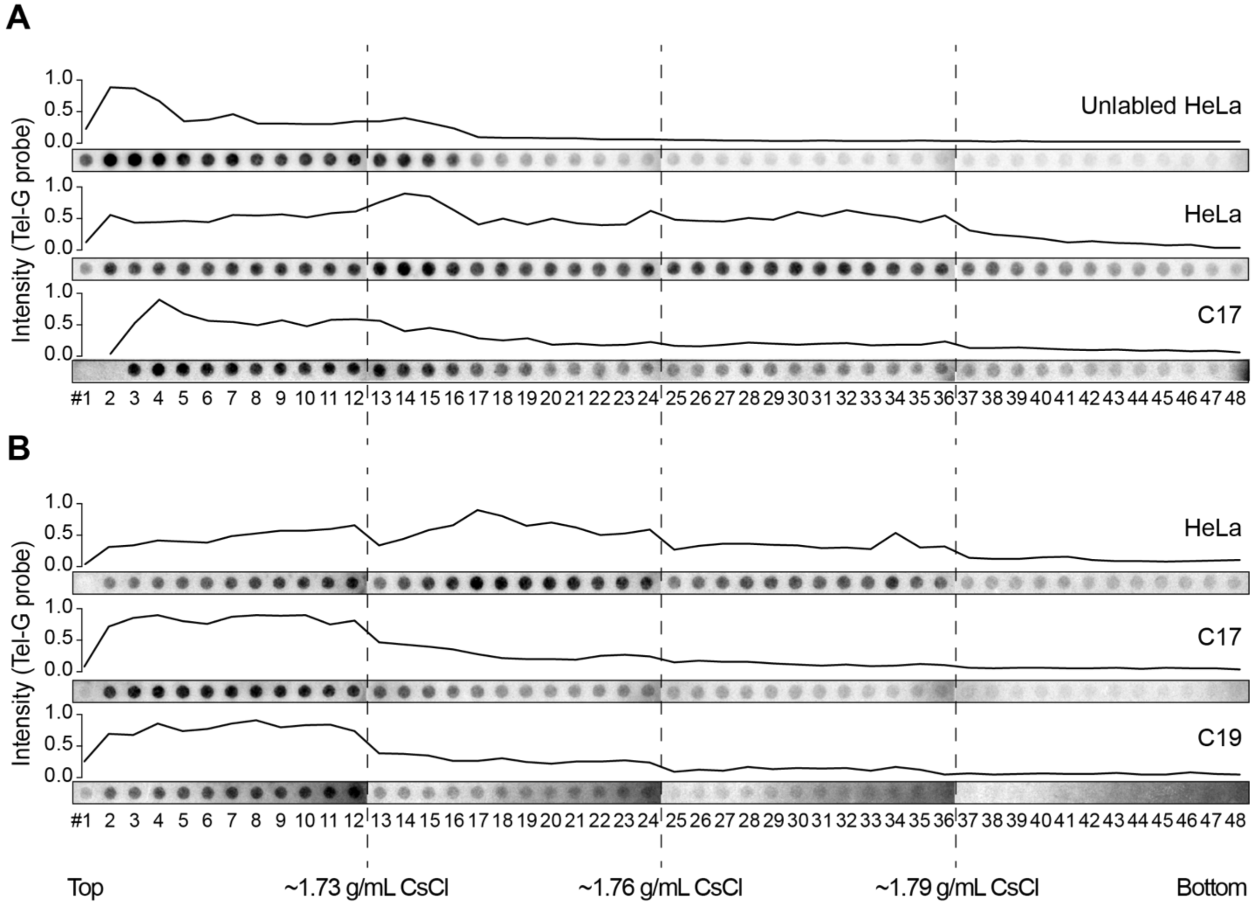


**Supplementary Figure 2. CsCl density-gradient ultracentrifugation with automated fractionation confirms reduced BrdU incorporation in RTEL1 ATPase-dead telomeres.**

**(A)** Validation of CsCl density-gradient ultracentrifugation using unlabeled HeLa, BrdU-labeled HeLa, and labeled ATPase-dead C17 cells. Fraction collection employed a TELEDYNE ISCO automated R1 fraction collector system. After fractionation, each telomere species was quantified by dot-blot hybridization with a telomere-specific probe (Tel-G). The results successfully distinguished labeled from unlabeled telomeres, but the resolution was insufficient to distinguish leading from lagging telomeres. Nonetheless, the data show that ATPase-dead C17 cells exhibited reduced BrdU incorporation into telomeres.

**(B)** An additional replicate using the automated fractionation system for parental HeLa and ATPase-dead (C17 and C19) cells. A trend similar to that in Figure 1A (manual fractionation) was observed, although the automated system could not resolve leading from lagging strands. “Top” and “bottom” correspond to the layer positions in Quick-Seal Ultra-Clear tubes (top = low CsCl density, bottom = high CsCl density).

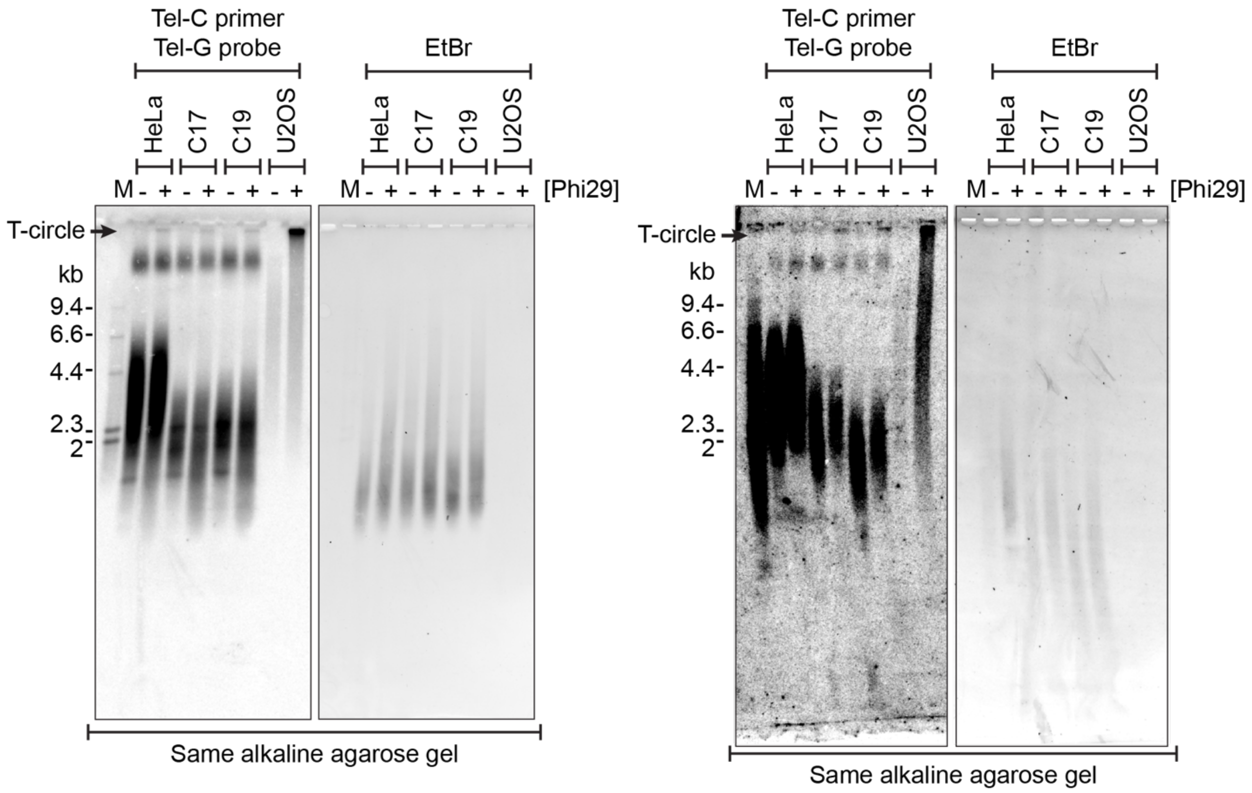


**Supplementary Figure 3. Additional replicates confirm similar T-circle levels in parental HeLa and ATPase-dead cells.**

Two additional independent replicates showing T-circle levels in parental HeLa and ATPase-dead (C17, C19) cells, with U2-OS (ALT-positive) cells as a positive control. RCA products were separated on a 0.6% alkaline agarose gel and detected by in-gel hybridization with a $^{32}$P-end-labeled Tel-G probe. EtBr staining was used to assess loading. Weak T-circle signals were observed in parental HeLa and ATPase-dead cells, in contrast to the strong signal from U2-OS.

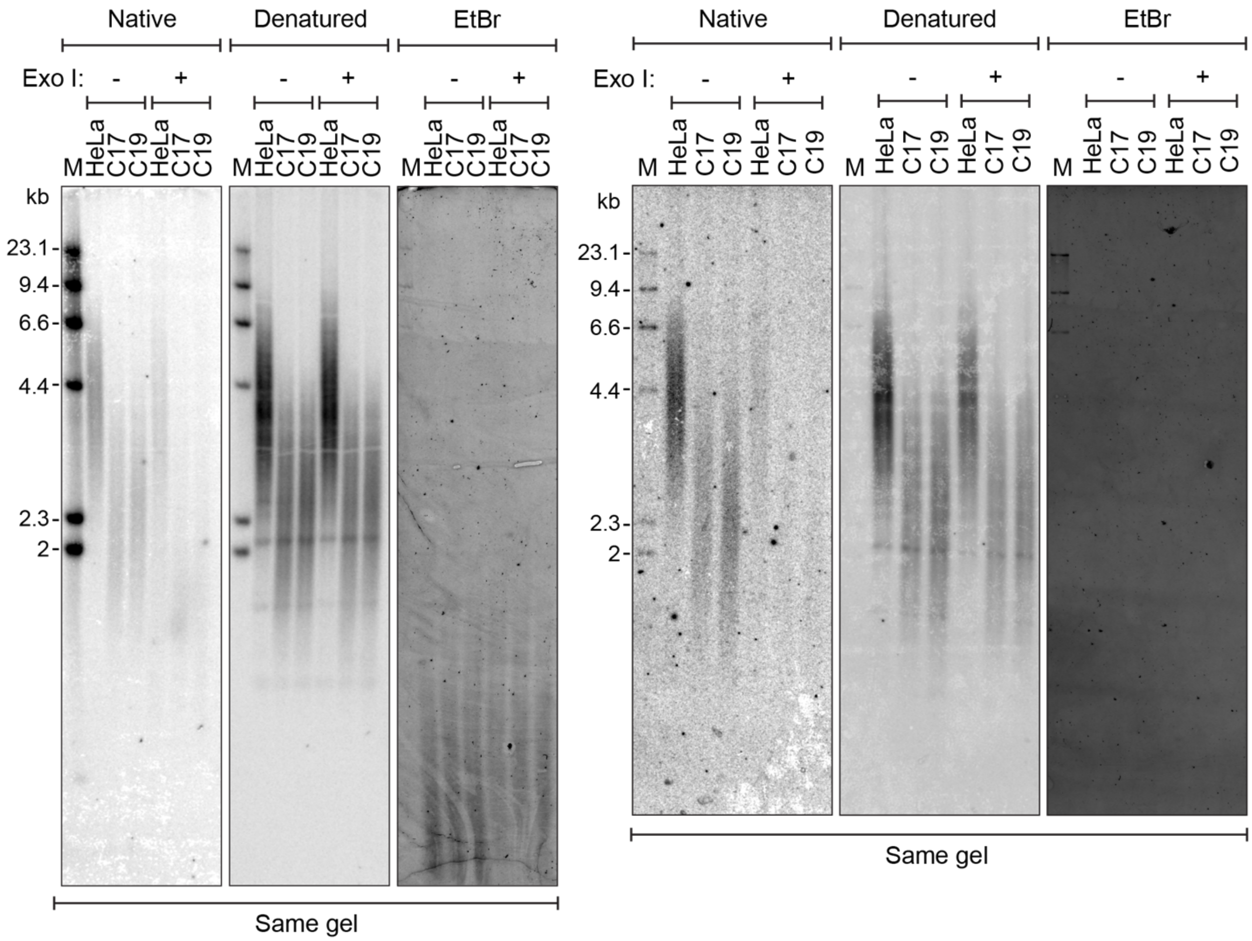


**Supplementary Figure 4. Additional replicates confirm similar G-strand overhangs in parental HeLa and ATPase-dead cells.**

Two additional independent replicates of in-gel hybridization analysis of telomeric G-strand exposure. Genomic DNA from parental HeLa and ATPase-dead (C17, C19) cells was separated on a native agarose gel, and single-stranded G-rich telomeric DNA was detected with a $^{32}$P-end-labeled Tel-C probe. After denaturation, total telomeric DNA was detected with the same probe. EtBr staining was used to assess loading.

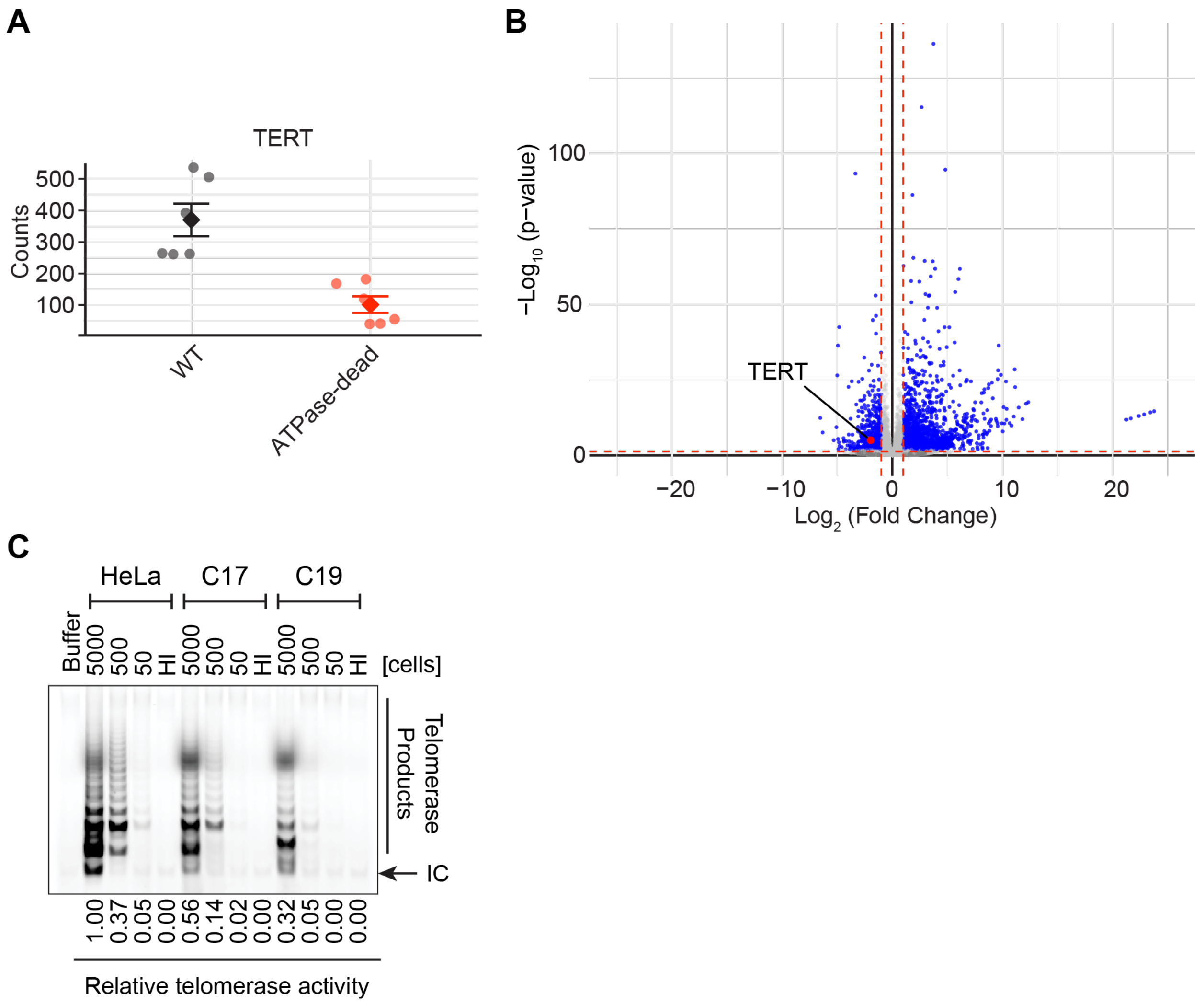


**Supplementary Figure 5. TERT mRNA levels and telomerase activity are reduced in RTEL1 ATPase-dead cells.**

**(A)** Raw counts of TERT mRNA in WT (C1, C15) and ATPase-dead (C17, C19) cells. Each dot represents an individual RNA-seq sample (n = 3 per cell line). TERT expression is markedly lower in ATPase-dead cells compared to WT cells.

**(B)** Volcano plots depicting the $\log_2$(fold change) versus $-\log_{10}$(p-value) for all genes in the RNA-seq data. The position of TERT is highlighted (red dot), showing its significant downregulation in ATPase-dead cells relative to other genes (gray dots). Red dashed lines indicate thresholds.

**(C)** An additional independent replicate of the TRAP assay shown in Figure 2D. TRAP products were resolved on a 10% native acrylamide gel, with the signal derived from Cy5-TS primer.

Relative telomerase activity was determined by quantifying the intensity of the TRAP ladder. The number of cells used for each reaction is indicated above the gel. HI, heat-inactivated samples; IC, internal control for PCR amplification.

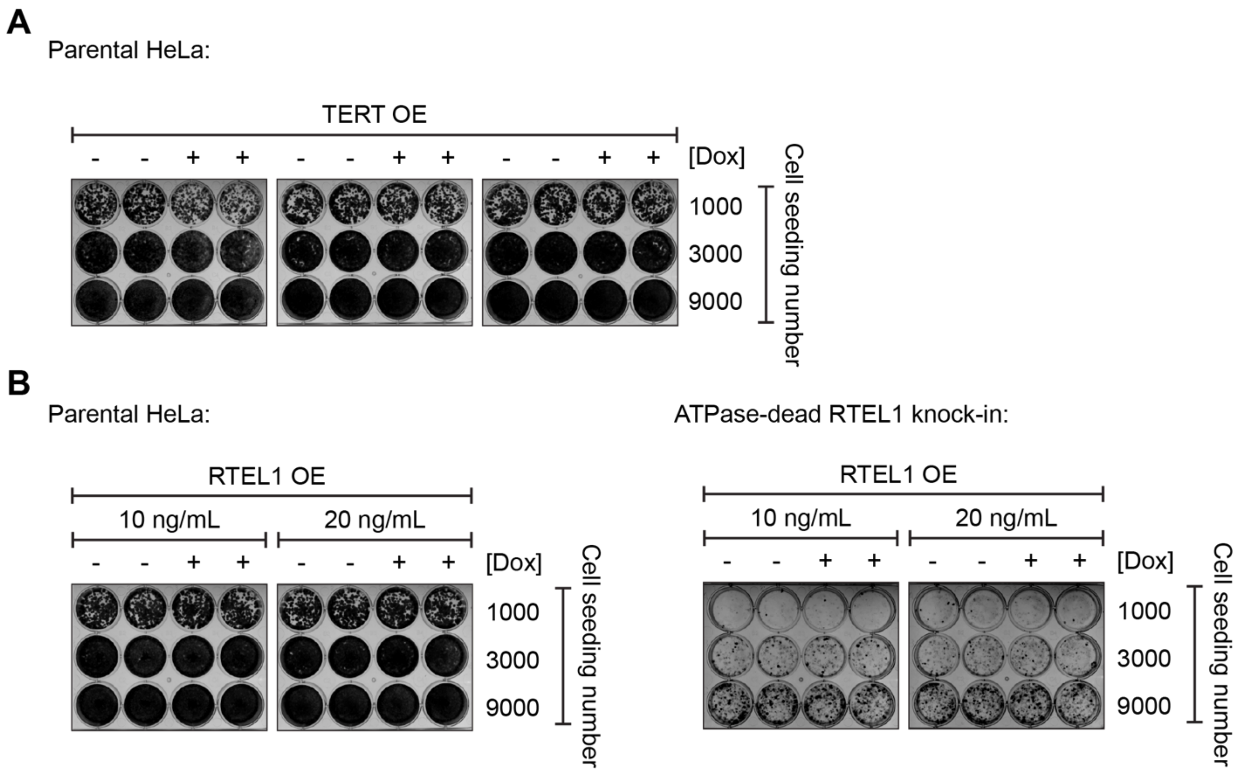


**Supplementary Figure 6. Additional colony formation experiments confirm that TERT or RTEL1 expression does not affect parental HeLa cell growth.**

**(A)** Colony formation assay. Parental HeLa cells (endogenous *RTEL1, TERT*-lentivirus) were seeded at the indicated numbers and treated with Dox to induce TERT overexpression (OE). Colonies were stained 8-9 days later. TERT OE had no effect on colony formation. Three technical replicates were generated from two biological replicates.

**(B)** Colony formation assay. Left: Parental HeLa cells (endogenous *RTEL1*, *RTEL1*-lentivirus) were seeded at the indicated numbers and treated with 10 or 20 ng/mL Dox to induce WT RTEL1 expression. Colonies were stained 9 days later. Two replicates were generated from the same experiment. Right: ATPase-dead RTEL1 cells were seeded at the indicated numbers and treated with 10 or 20 ng/mL Dox to induce WT RTEL1 expression. Colonies were stained 9 days later. The overall workflow is the same as for TERT induction, as shown in Figure 3B. Two replicates were generated from the same experiment.

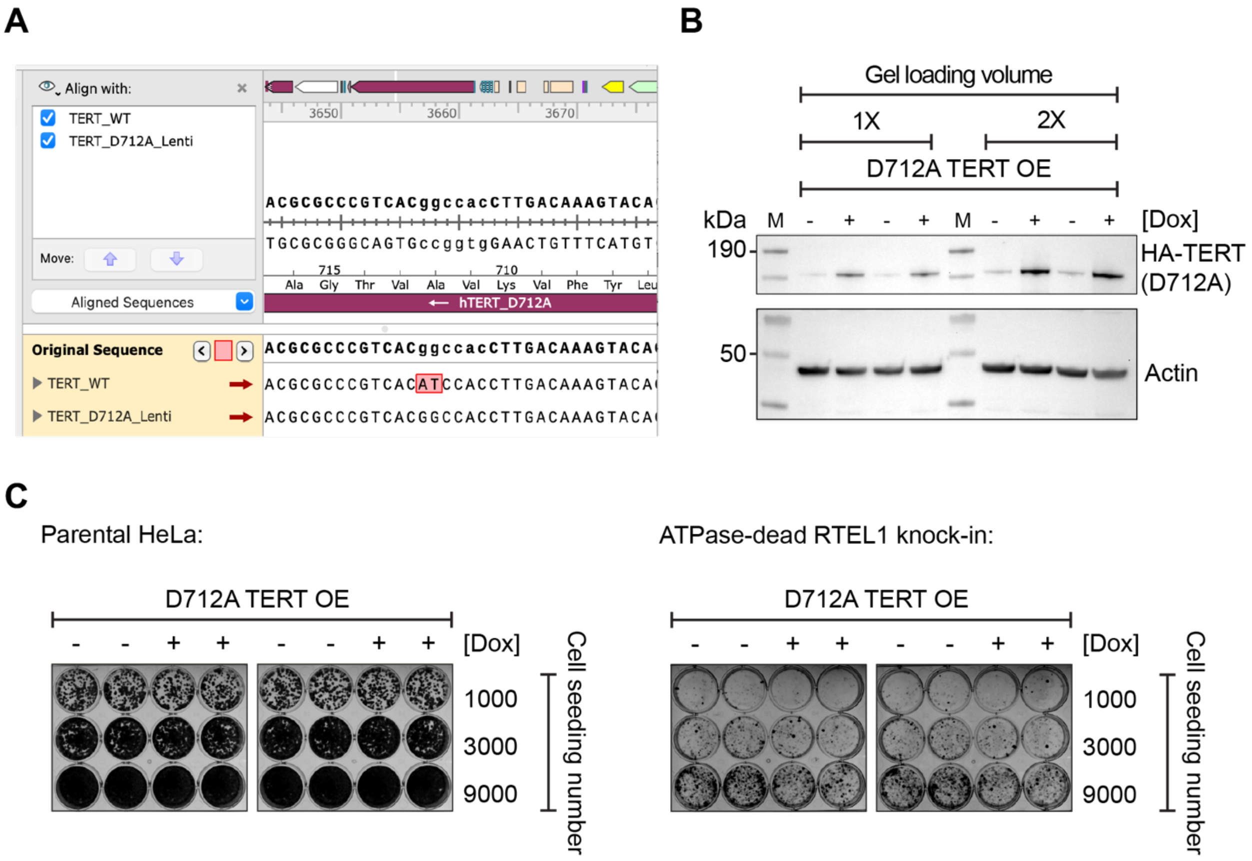


**Supplementary Figure 7. Overexpression of catalytically inactive D712A TERT does not affect colony formation in parental or ATPase-dead cells.**

**(A)** Validation of the D712A TERT transfer vector. A SnapGene window shows sequencing results of the D712A TERT transfer vector aligned to the WT TERT sequence, confirming the introduced D712A substitution.

**(B)** Western blot validation of Dox-inducible HA-tagged D712A TERT overexpression. Leaky expression in the absence of Dox is due to basal activity of the Tet-inducible promoter.

**(C)** Colony formation assay. Left: Parental HeLa cells (endogenous *RTEL1*, D712A *TERT*-lentivirus) were seeded at the indicated amounts and treated with Dox to induce D712A TERT OE. Colonies were stained 9-11 days later. D712A TERT OE had no effect on colony formation. Two replicates were generated from the same experiment. Right: ATPase-dead cells were seeded at the indicated amounts and treated with Dox to induce D712A TERT OE. Colonies were stained 9-11 days later. D712A TERT OE had no effect on colony formation. Two replicates were generated from the same experiment.

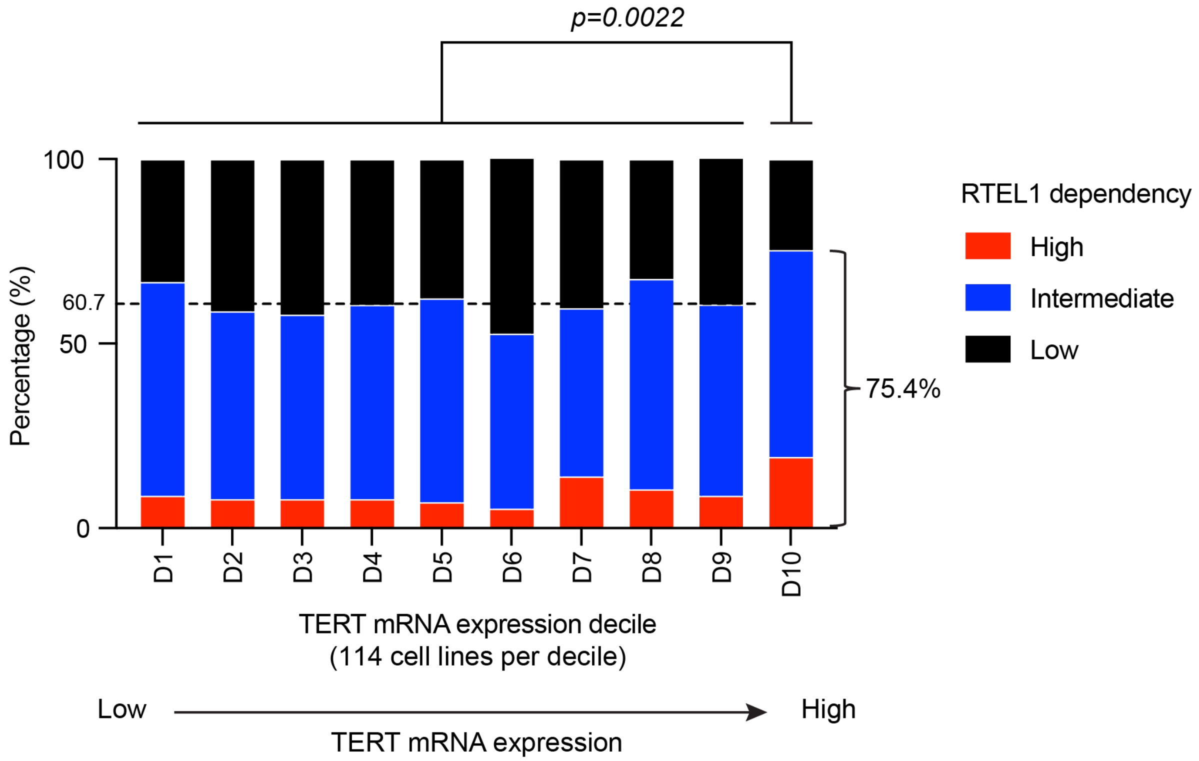


**Supplementary Figure 8. Cell lines with the highest TERT expression levels show the greatest RTEL1 dependency.**

RTEL1 dependency was compared across TERT mRNA expression deciles (n=1140 cell lines, 114 cell lines per decile). Cell lines in D10 showed a significantly higher proportion of RTEL1 dependency than the remaining nine deciles combined (75.4% vs 60.7%; *p=0.0022*, Fisher's exact test). This increase is primarily driven by the high-dependency subgroup (19.3% vs 8.7%; *p*=0.0012), whereas the intermediate-dependency subgroup had little change (56.1% vs 52%; *p*=0.43). RTEL1 dependency was defined as: low (> −0.5), intermediate (−0.5 to −1), and high (< −1). RTEL1 CRISPR (Chronos) gene effect scores and TERT mRNA levels were obtained from the DepMap database.

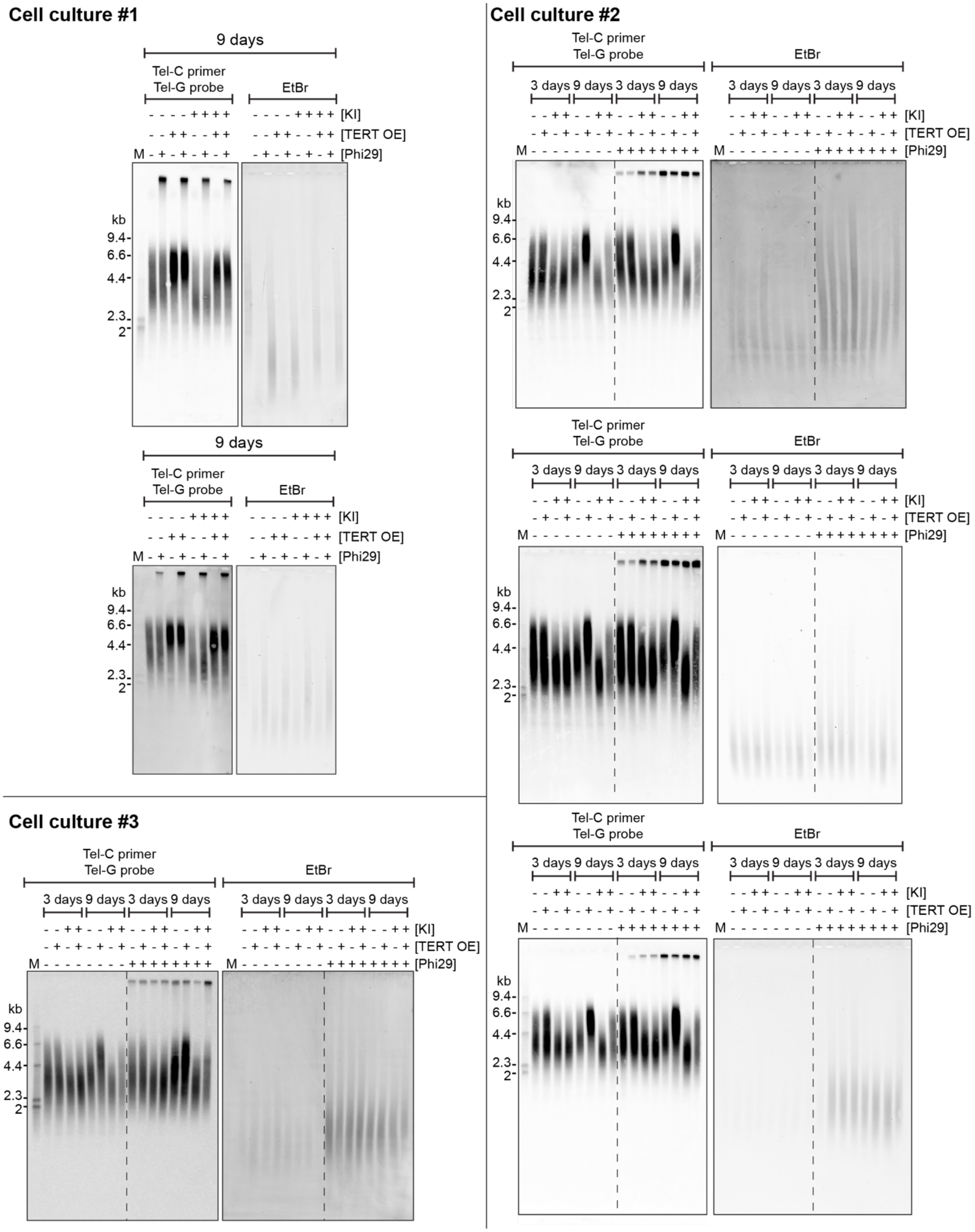

Cell culture #1
9 days
Tel-C primer
Tel-G probe
EtBr
- - - - + + + + [KI]
- - + + - - + + [TERT OE]
M - + - + - + - + [Phi29]
kb
9.4
6.6
4.4
2.3
2
Cell culture #2
3 days 9 days 3 days 9 days
- - + + - - + + - - + + - - + + [KI]
- + - + - + - + - + - + - + - + [TERT OE]
M - - - - - - - - + + + + + + + + [Phi29]
Cell culture #3

**Supplementary Figure 9. Analysis of telomere lengths and T-circles from three independent cell cultures after knock-in (KI) of ATPase-dead RTEL1 and TERT overexpression.**

After knocking in ATPase-dead RTEL1, cells were treated with Dox to induce TERT overexpression, as outlined in Figure 3B. Genomic DNA was isolated 3 or 9 days post-induction. T-circles were amplified by RCA with a Tel-C primer across three independent cell cultures, separated on a 0.6% alkaline agarose gel, and detected by in-gel hybridization with a $^{32}$P-end-labeled Tel-G probe. A transient increase was observed in one cell culture (right panel), whereas no consistent change was detected in the other two. EtBr staining was used to assess loading.

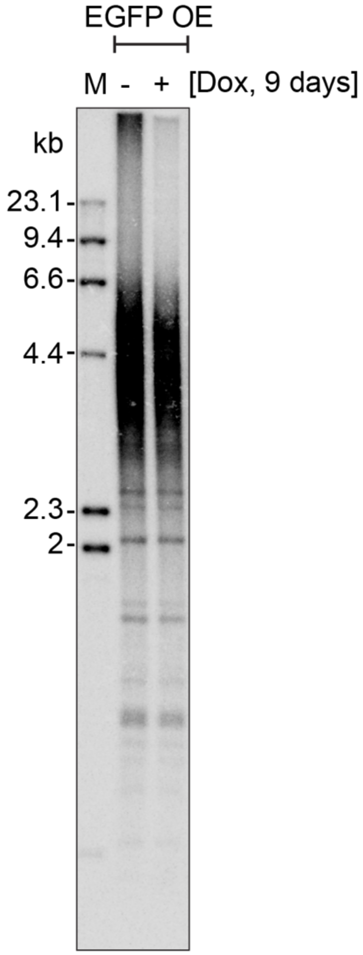


**Supplementary Figure 10. No increase in telomere length after EGFP overexpression.**

To control for possible effects of Dox on telomere length in the TERT overexpression experiments, cells were treated with Dox to induce EGFP overexpression (OE). Genomic DNA was isolated 9 days post-induction and separated on a 0.8% agarose gel. Telomere DNA was detected by in-gel hybridization following denaturation with a $^{32}$P-end-labeled Tel-G probe.

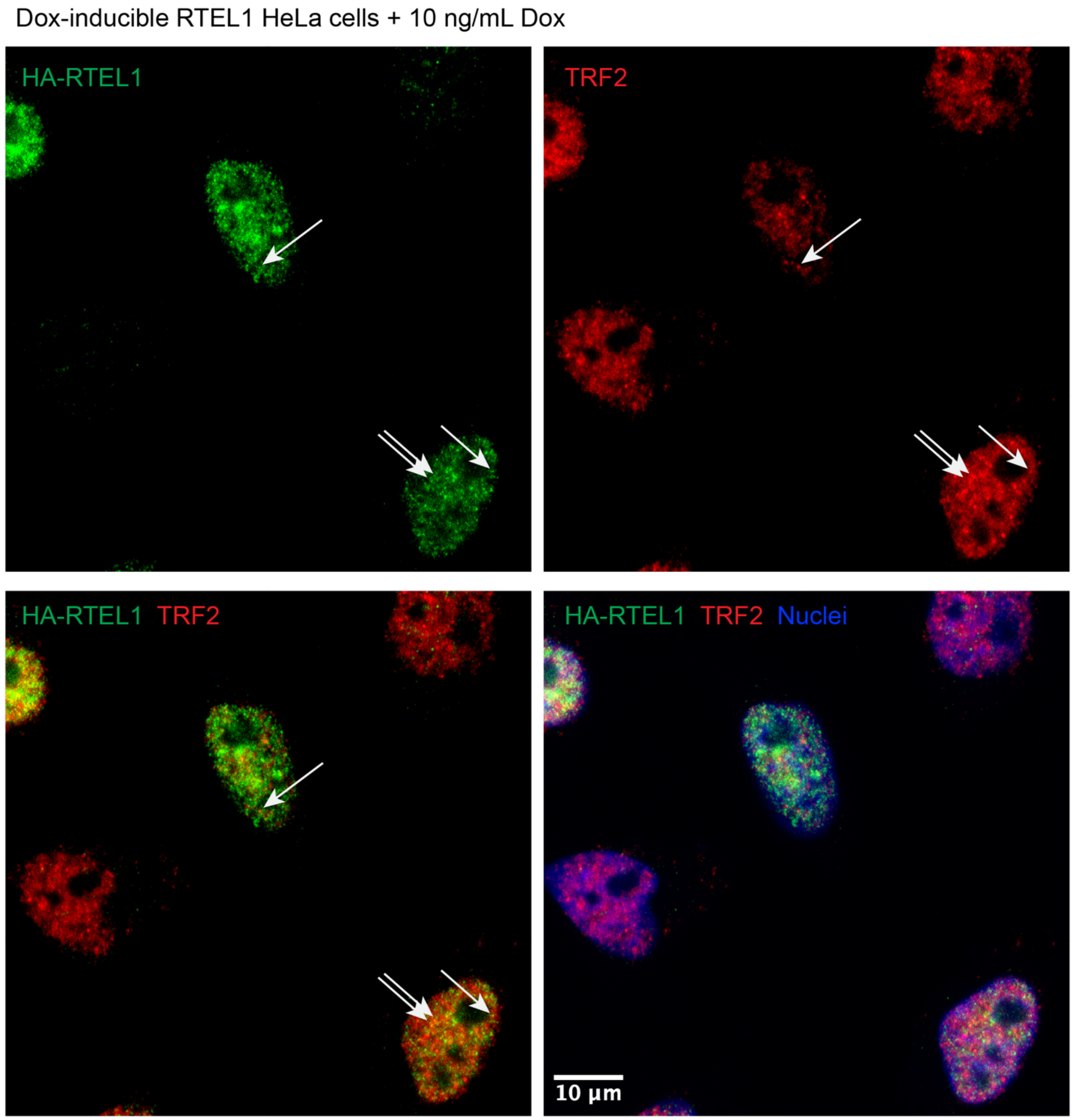


**Supplementary Figure 11. Immunostaining analysis of HA-RTEL1 localization relative to TRF2.**

HA-RTEL1 and TRF2 were immunostained after cells were treated with 10 ng/mL Dox for 48 hours to induce WT HA-RTEL1 expression. Images were captured on a Nikon AXR laser scanning confocal microscope equipped with a 100X NA 1.45 objective. Green, HA-RTEL1; Red, TRF2; Blue, nuclei (DAPI). Arrows indicate examples of colocalized foci.

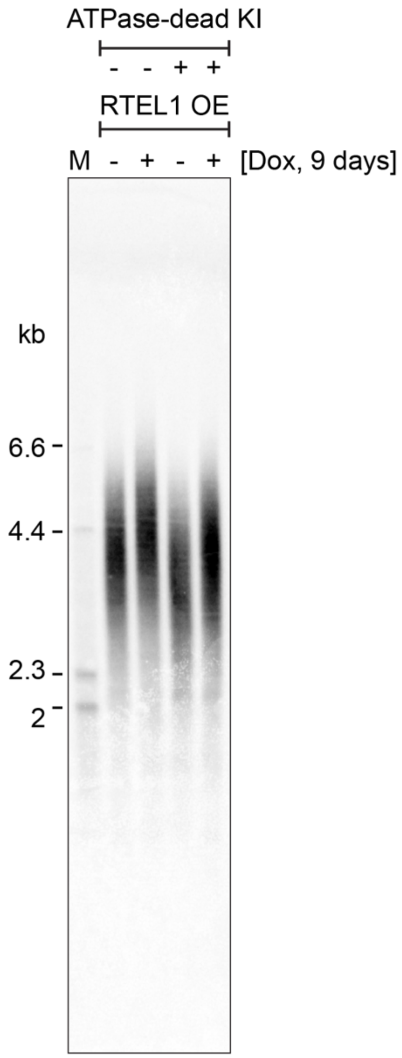


**Supplementary Figure 12. Expression of the longest RTEL1 isoform facilitates telomere elongation.**

Telomere restriction fragment (TRF) analysis of telomere length. Parental HeLa or ATPase-dead RTEL1 knock-in (KI) cells, both transduced with RTEL1-lentivirus, were treated with 20 ng/mL Dox to induce RTEL1 expression, as outlined in Figure 3B. Genomic DNA was isolated on day 9 post-induction. Digested genomic DNA was separated on a 0.6% alkaline agarose gel, and telomeric DNA was detected by in-gel hybridization with a $^{32}$P-end-labeled Tel-G probe.

**Figure 1A**

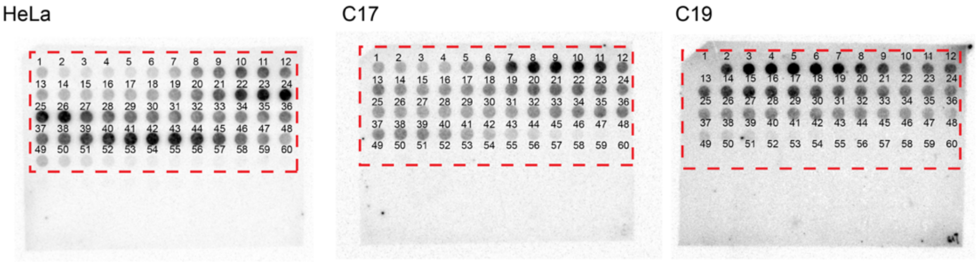


**Figure 1B**

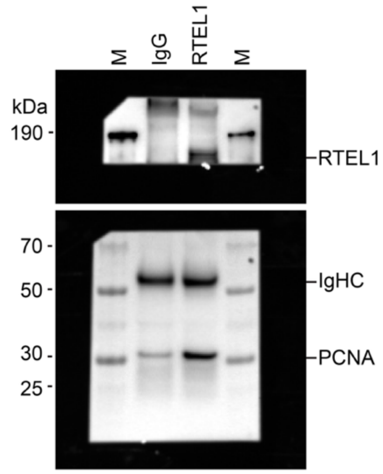


**Figure 1C**

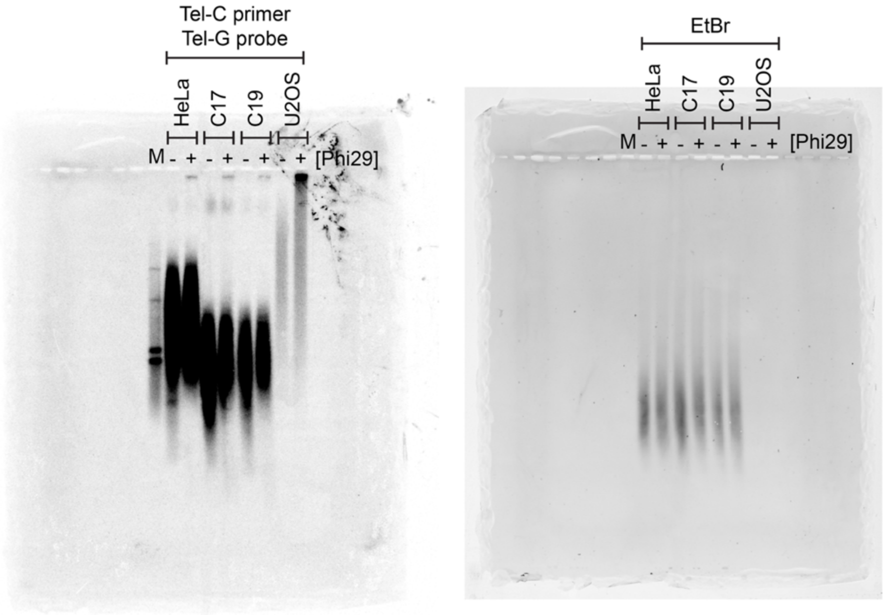


**Supplementary Figure 13. Uncropped images.**

**Figure 1D**

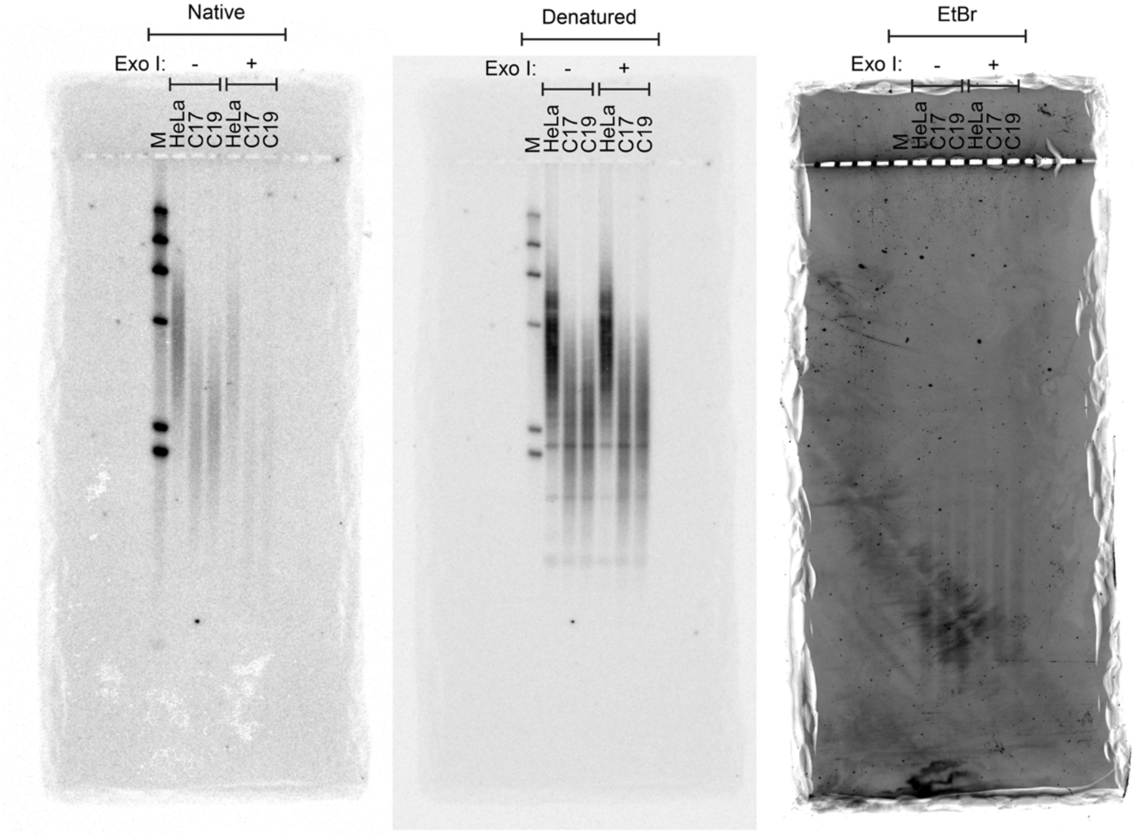


**Figure 2D**

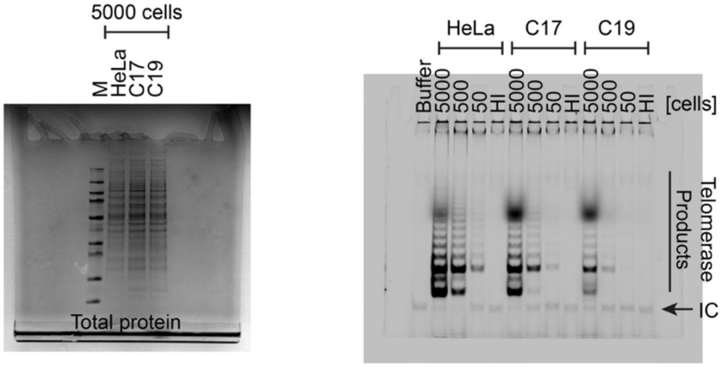


**Figure 2E**

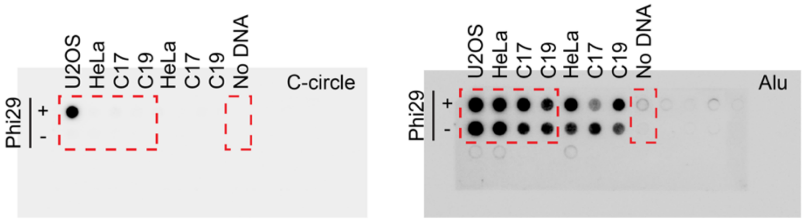


**Supplementary Figure 13. Uncropped images (continued).**

### Figure 3A

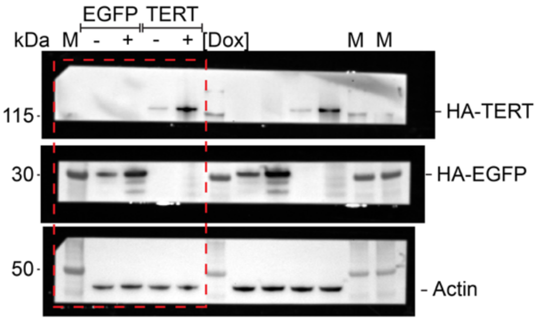


### Figure 3C

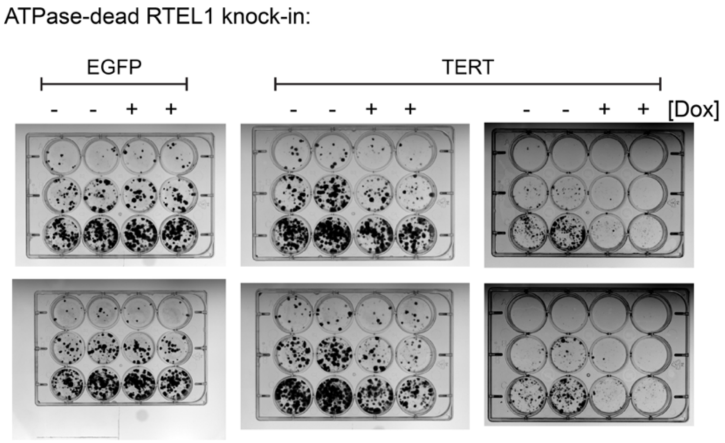


### Figure 4A

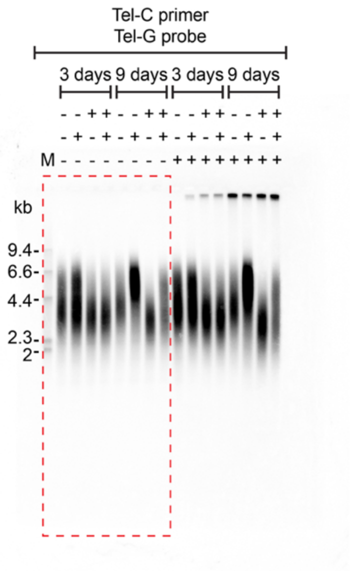


### Figure 4C

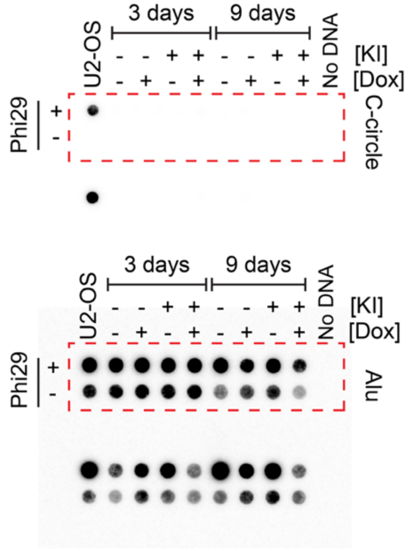


**Supplementary Figure 13. Uncropped images (continued).**

### Figure 5A

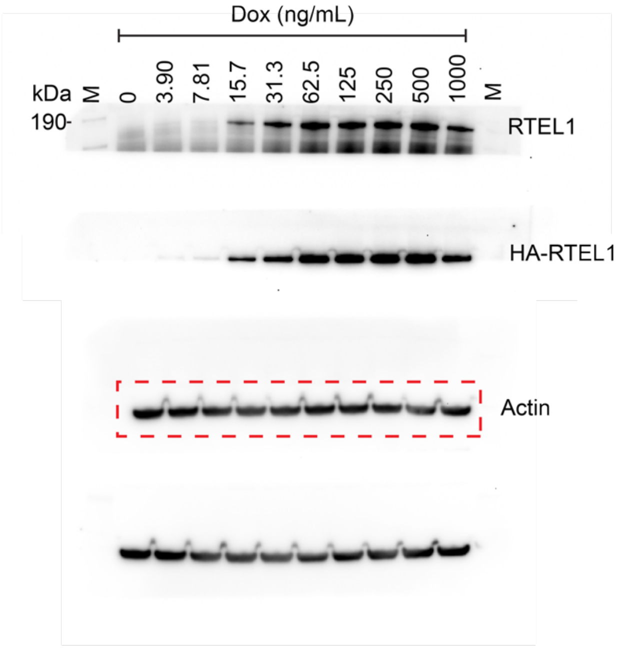


### Figure 5D

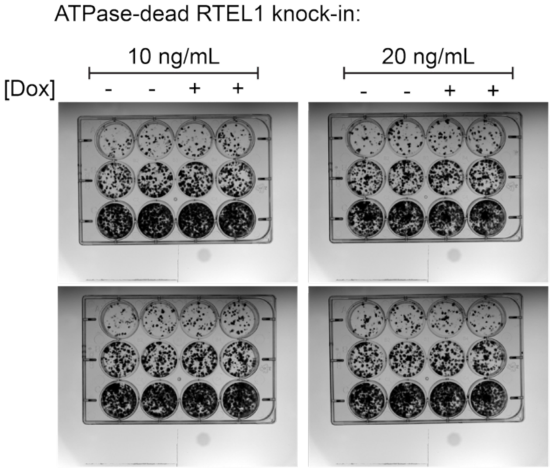


### Figure 5E

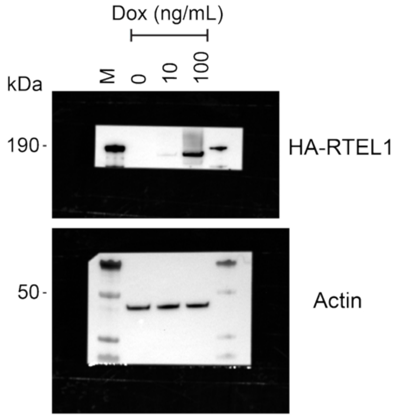


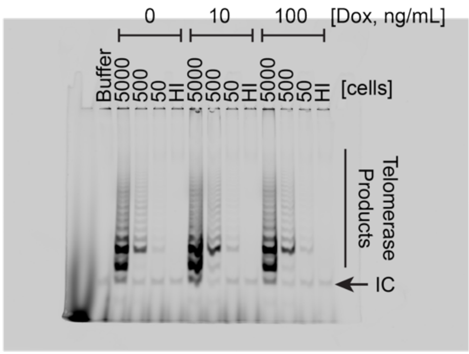


**Supplementary Figure 13. Uncropped images (continued).**

**Supplementary Transfer Vector Sequences**

> EGFP_lenti_vector

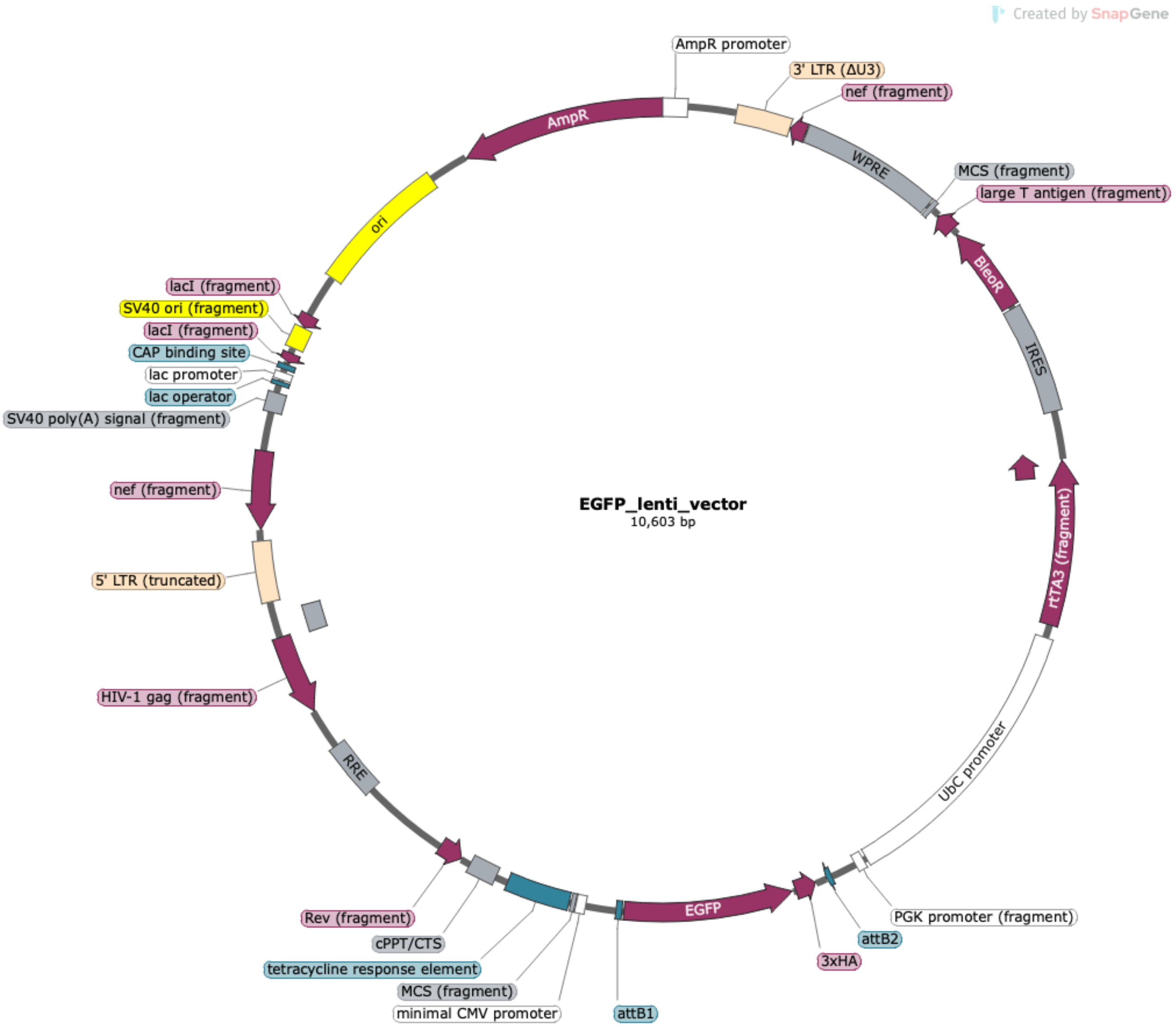


ACTCTTCCTTTTTCAATATTATTGAAGCATTTATCAGGGTTATTGTCTCATGAGCGGATACATATTTGAATGTA

TTTAGAAAAATAAACAAATAGGGGTTCCGCGCACATTTCCCCGAAAAGTGCCACCTGACGTCTAAGAAAC

CATTATTATCATGACATTAACCTATAAAAATAGGCGTATCACGAGGCCCTTTCGTCTTCAAGAATGATCTAG

CCCTTTCCTTAATTAACCCGGGCCTCTCACTCTCTGATATTCATTTCTTTGCAAGTTATAAATACTGAATAATA

AGATGACATGAACTACTACTGCTAGAGATTTTCCACACTGACTAAAAGGGTCTGAGGGATCTCTAGTTACC

AGAGTCACACAACAGACGGGCACACACTACTTGAAGCACTCAAGGCAAGCTTTATTGAGGCTTAAGCAG

TGGGTTCCCTAGTTAGCCAGAGAGCTCCCAGGCTCAGATCTGGTCTAACCAGAGAGACCCAGTACAAGC

AAAAAGCAGATCTTGTCTTCGTTGGGAGTGAATTAGCCCTTCCAGTCCCCCCTTTTCTTTTAAAAAGTGGC
TAAGATCTACAGCTGCCTTGTAAGTCATTGGTCTTAAAGGATCTCAGGCGGGGAGGCGGCCCAAAGGGA
GATCCGACTCGTCTGAGGGCGAAGGCGAAGACGCGGAAGAGGCCGCAGAGCCGGCAGCAGGCCGCG
GGAAGGAAGGTCCGCTGGATTGAGGGCCGAAGGGACGTAGCAGAAGGACGTCCCGCGCAGAATCCAG
GTGGCAACACAGGCGAGCAGCCAAGGAAAGGACGATGATTTCCCCGACAACACCACGGAATTGTCAGT
GCCCAACAGCCGAGCCCCTGTCCAGCAGCGGGCAAGGCAGGCGGCGATGAGTTCCGCCGTGGCAATAG
GGAGGGGGAAAGCGAAAGTCCCGGAAAGGAGCTGACAGGTGGTGGCAATGCCCCAACCAGTGGGGGT
TGCGTCAGCAAACACAGTGCACACCACGCCACGTTGCCTGACAACGGGCCACAACTCCTCATAAAGAGA
CAGCAACCAGGATTTATACAAGGAGGAGAAAATGAAAGCCATACGGGAAGCAATAGCATGATACAAAG
GCATTAAAGCAGCGTATCCACATAGCGTAAAAGGAGCAACATAGTTAAGAATACCAGTCAATCTTTCACA
AATTTTGTAATCCAGAGGTTGATTAGGATCTATCGCGATAAGCTTGATATCGAATTGGGAGGGACTAAACA
ACAACAATTGCATTCATTTTATGTTTCAGGTTCAGGGGGAGGTGTGGGAGGTTTTTTAAAGCAAGTAAAA
CCTCTACAAATGTGGTATGGCTGATTATGATCTAGAGTCGCGGCCGCTTTACTTGTACATCAGTCCTGCTCC
TCGGCCACGAAGTGCACGCAGTTGCCGGCCGGGTCGCGCAGGGCGAACTCCCGCCCCCACGGCTGCTC
GCCGATCTCGGTCATGGCCGGCCCGGAGGCGTCCCGGAAGTTCGTGGACACGACCTCCGACCACTCGGC
GTACAGCTCGTCCAGGCCGCGCACCCACACCCAGGCCAGGGTGTTGTCCGGCACCACCTGGTCCTGGAC
CGCGCTGATGAACAGGGTCACGTCGTCCCGGACCACACCGGCGAAGTCGTCCTCCACGAAGTCCCGGGA
GAACCCGAGCCGGTCGGTCCAGAACTCGACCGCTCCGGCGACGTCGCGCGCGGTGAGCACCGGAACGG
CACTGGTCAACTTGGCCATGGTGGACCGGTAAGCTTATCATCGTGTTTTTCAAAGGAAAACCACGTCCCC
GTGGTTCGGGGGGCCTAGACGTTTTTTTAACCTCGACTAAACACATGTAAAGCATGTGCACCGAGGCCCC
AGATCAGATCCCATACAATGGGGTACCTTCTGGGCATCCTTCAGCCCCTTGTTGAATACGCTTGAGGAGAG
CCATTTGACTCTTTCCACAACTATCCAACTCACAACGTGGCACTGGGGTTGTGCCGCCTTTGCAGGTGTAT
CTTATACACGTGGCTTTTGGCCGCAGAGGCACCTGTCGCCAGGTGGGGGGTTCCGCTGCCTGCAAAGGG
TCGCTACAGACGTTGTTTGTCTTCAAGAAGCTTCCAGAGGAACTGCTTCCTTCACGACATTCAACAGACCT
TGCATTCCTTTGGCGAGAGGGGAAAGACCCCTAGGAATGCTCGTCAAGAAGACAGGGCCAGGTTTCCGG
GCCCTCACATTGCCAAAAGACGGCAATATGGTGGAAAATAACATATAGACAAACGCACACCGGCCTTATT
CCAAGCGGCTTCGGCCAGTAACGTTAGGGGGGGGGGAGGGGGGGGGGAGAGGGGCGGAATTCCTCTA
GTGCGGCCGCGGATCCTTACTTAGTTACCCGGGGAGCATGTCAAGGTCAAAATCGTCAAGAGCGTCAGC
AGGCAGCATATCAAGGTCAAAGTCGTCAAGGGCATCGGCTGGGAGCATGTCTAAGTCAAAATCGTCAAG

GGCGTCGGTCGGCCCGCCGCTTTCGCACTTTAGCTGTTTCTCCAGGCCACATATGATTAGTTCCAGGCCGA
AAAGGAAGGCAGGTTCGGCTCCCTGCCGGTCGAACAGCTCAATTGCTTGTCTCAGAAGTGGGGGCATAG
AATCGGTGGTAGGTGTCTCTCTTTCCTCTTTTGCTACTTGATGCTCCTGTTCCTCCAATACGCAGCCCAGTGT
AAAGTGGCCCACGGCGGACAGAGCGTACAGTGCGTTCTCCAGGGAGAAGCCTTGCTGACACAGGAACG
CGAGCTGATTTTCCAGGGTTTCGTACTGTTTCTCTGTTGGGCGGGTGCCGAGATGCACTTTAGCCCCGTCG
CGATGTGAGAGGAGAGCACAGCGGTATGACTTGGCGTTGTTCCGCAGAAAGTCTTGCCATGACTCGCCTT
CCAGGGGGCAGAAGTGGGTATGATGCCTGTCCAGCATCTCGATTGGCAGGGCATCGAGCAGGGCCCGCT
TGTTCTTCACGTGCCAGTACAGGGTAGGCTGCTCAACTCCCAGCTTTTGAGCGAGTTTCCTTGTCGTCAGG
CCTTCGATACCGACACCATTGAGTAATTCCAGAGCTCCGTTTATGACTTTGCTCTTGTCCAGTCTAGACATG
GTGAATTCGACTGCAGGACCGGTACGGGGTGGAGATCCGAGCTCGGTACCAAGCTTCGTCTAACAAAAA
AGCCAAAAACGGCCAGAATTTAGCGGACAATTTACTAGTCTAACACTGAAAATTACATATTGACCCAAAT
GATTACATTTCAAAAGGTGCCTAAAAAACTTCACAAAACACACTCGCCAACCCCGAGCGCATAGTTCAAA
ACCGGAGCTTCAGCTACTTAAGAAGATAGGTACATAAAACCGACCAAAGAAACTGACGCCTCACTTATCC
CTCCCCTCACCAGAGGTCCGGCGCCTGTCGATTCAGGAGAGCCTACCCTAGGCCCGAACCCTGCGTCCTG
CGACGGAGAAAAGCCTACCGCACACCTACCGGCAGGTGGCCCCACCCTGCATTATAAGCCAACAGAACG
GGTGACGTCACGACACGACGAGGGCGCGCTCCCAAAGGTACGGGTGCACTGCCCAACGGCACCGCC
ATAACTGCCGCCCCCGCAACAGACGACAAACCGAGTTCTCCAGTCAGTGACAAACTTCACGTCAGGGTC
CCCAGATGGTGCCCCAGCCCATCTCACCCGAATAAGAGCTTTCCCGCATTAGCGAAGGCCTCAAGACCTT
GGGTTCTTGCCGCCCACCATGCCCCCCACCTTGTTTCAACGACCTCACAGCCCGCCTCACAAGCGTCTTCC
ATTCAAGACTCGGGAACAGCCGCCATTTTGCTGCGCTCCCCCCAACCCCCAGTTCAGGGCAACCTTGCTC
GCGGACCCAGACTACAGCCCTTGGCGGTCTCTCCACACGCTTCCGTCCCACCGAGCGGCCCGGCGGCCA
CGAAAGCCCCGGCCAGCCCAGCAGCCCGCTACTCACCAAGTGACGATCACAGCGATCCACAAACAAGA
ACCGCGACCCAAATCCCGGCTGCGACGGAACTAGCTGTGCCACACCCGGCGCGTCCTTATATAATCATCG
GCGTTCACCGCCCCACGGAGATCCCTCCGCAGAATCGCCGAGAAGGGACTACTTTTCCTCGCCTGTTCCG
CTCTCTGGAAAGAAAACCAGTGCCCTAGAGTCACCCAAGTCCCGTCCTAAAATGTCCTTCTGCTGATACTG
GGGTTCTAAGGCCGAGTCTTATGAGCAGCGGGCCGCTGTCCTGAGCGTCCGGGCGGAAGGATCAGGAC
GCTCGCTGCGCCCTTCGTCTGACGTGGCAGCGCTCGCCGTGAGGAGGGGGGCGCCCGCGGGAGGCGCC
AAAACCCGGCGCGGAGGCCAGATCTTGGGTGGGTTACTCCAGACTGCCTTGGGAAAAGCGCCTCCCCTA
CCCGGTAGAATTTCTAGTTTAATTAATCATTACTAAGCGTAGTCTGGGACGTCGTATGGGTATTCGAACCGC

GGGCCCTCTAGACTCGAGCGGCCGCCACTGTGCTGGATATCAACCACTTTGTACAAGAAAGCTGGGTCTA
GATATCTCGAGGCGGCCGCTTATCCGGATTCGAATCATTAGGATCCAGCGTAATCTGGAACGTCATAAGGA
TACGATCCTGCATAGTCCGGGACGTCATAGGGATAGCCCGCATAGTCAGGAACATCGTATGGGTACCCGCC
GGTGCCCTTGTACAGCTCGTCCATGCCGAGAGTGATCCCGGCGGCGGTCACGAACTCCAGCAGGACCAT
GTGATCGCGCTTCTCGTTGGGGTCTTTGCTCAGGGCGGACTGGGTGCTCAGGTAGTGGTTGTCGGGCAGC
AGCACGGGGCCGTCGCCGATGGGGGTGTTCTGCTGGTAGTGGTCGGCGAGCTGCACGCTGCCGTCCTCG
ATGTTGTGGCGGATCTTGAAGTTCACCTTGATGCCGTTCTTCTGCTTGTCGGCCATGATATAGACGTTGTGG
CTGTTGTAGTTGTACTCCAGCTTGTGCCCCAGGATGTTGCCGTCCTCCTTGAAGTCGATGCCCTTCAGCTCG
ATGCGGTTCACCAGGGTGTCGCCCTCGAACTTCACCTCGGCGCGGGTCTTGTAGTTGCCGTCGTCCTTGAA
GAAGATGGTGCGCTCCTGGACGTAGCCTTCGGGCATGGCGGACTTGAAGAAGTCGTGCTGCTTCATGTGG
TCGGGGTAGCGGCTGAAGCACTGCACGCCGTAGGTCAGGGTGGTCACGAGGGTGGGCCAGGGCACGG
GCAGCTTGCCGGTGGTGCAGATGAACTTCAGGGTCAGCTTGCCGTAGGTGGCATCGCCCTCGCCCTCGCC
GGACACGCTGAACTTGTGGCCGTTTACGTCGCCGTCCAGCTCGACCAGGATGGGCACCACCCCGGTGAA
CAGCTCCTCGCCCTTGCTCACCATGGTGGCGCCAGCCTGCTTTTTTGTACAAACTTGTTGATATCTGCAGA
ATTCCACCACACTGGACTAGTTCGGGGCCGCGGAGGCTGGATCGGTCCCGGTGTCTTCTATGGAGGTCAA
AACAGCGTGGATGGCGTCTCCAGGCGATCTGACGGTTCACTAAACGAGCTCTGCTTATATAGGCCTCCCA
CCGTACACGCCTACCTCGACCCGGGTACCGAGCTCGACTTTCACTTTTCTCTATCACTGATAGGGAGTGGT
AAACTCGACTTTCACTTTTCTCTATCACTGATAGGGAGTGGTAAACTCGACTTTCACTTTTCTCTATCACTG
ATAGGGAGTGGTAAACTCGACTTTCACTTTTCTCTATCACTGATAGGGAGTGGTAAACTCGACTTTCACTTT
TCTCTATCACTGATAGGGAGTGGTAAACTCGACTTTCACTTTTCTCTATCACTGATAGGGAGTGGTAAACTC
GACTTTCACTTTTCTCTATCACTGATAGGGAGTGGTAAAGGATCCTAGTCCAAACTGGATCTCTGCTGTCCC
TGTAATAAACCCGAAAATTTTGAATTTTTGTAATTTGTTTTTGTAATTCTTTAGTTTGTATGTCTGTTGCTATTA
TGTCTACTATTCTTTCCCCTGCACTGTACCCCCCAATCCCCCCTTTTCTTTTAAAATTGTGGATGAATACTGC
CATTTGTGAATTCGGCGATACCGTCGAGATCCGTTCACTAATCGAATGGATCTGTCTCTGTCTCTCTCTCCAC
CTTCTTCTTCTATTCCTTCGGGCCTGTCGGGTCCCCTCGGGGTTGGGAGGTGGGTCTGAAACGATAATGGT
GAATATCCCTGCCTAACTCTATTCACTATAGAAAGTACAGCAAAAACTATTCTTAAACCTACCAAGCCTCCT
ACTATCATTATGAATAATTTTATATACCACAGCCAATTTGTTATGTTAAACCAATTCCACAAACTTGCCCATTT
ATCTAATTCCAATAATTCTTGTTCATTCTTTTCTTGCTGGTTTTGCGATTCTTCAATTAAGGAGTGTATTAAGC
TTGTGTAATTGTTAATTTCTCTGTCCCACTCCATCCAGGTCGTGTGATTCCAAATCTGTTCCAGAGATTTATTA

CTCCAACTAGCATTCCAAGGCACAGCAGTGGTGCAAATGAGTTTTCCAGAGCAACCCCAAATCCCCAGG
AGCTGTTGATCCTTTAGGTATCTTTCCACAGCCAGGATTCTTGCCTGGAGCTGCTTGATGCCCCAGACTGT
GAGTTGCAACAGATGCTGTTGCGCCTCAATAGCCCTCAGCAAATTGTTCTGCTGCTGCACTATACCAGACA
ATAATTGTCTGGCCTGTACCGTCAGCGTCATTGACGCTGCGCCCATAGTGCTTCCTGCTGCTCCCAAGAAC
CCAAGGAACAAAGCTCCTATTCCCACTGCTCTTTTTCTCTCTGCACCACTCTTCTCTTTGCCTTGGTGGGT
GCTACTCCTAATGGTTCAATTTTTACTACTTTATATTTATATAATTCACTTCTCCAATTGTCCCTCATATCTCCT
CCTCCAGGTCTGAAGATCAGCGGCCGGCCGCTTGCTGTGCGGTGGTCTTACTTTTGTTTTGCTCTTCCTCTA
TCTTGTCTAAAGCTTCCTTGGTGTCTTTTATCTCTATCCTTTGATGCACACAATAGAGGGTTGCTACTGTATTA
TATAATGATCTAAGTTCTTCTGATCCTGTCTGAAGGGATGGTTGTAGCTGTCCCAGTATTTGTCTACAGCCTT
CTGATGTTTCTAACAGGCCAGGATTAACTGCGAATCGTTCTAGCTCCCTGCTTGCCCATACTATATGTTTTAA
TTTATATTTTTTCTTTCCCCCTGGCCTTAACCGAATTTTTTCCCATCGCGATCTAATTCTCCCCCGCTTAATAC
TGACGCTCTCGCACCCATCTCTCTCCTTCTAGCCTCCGCTAGTCAAAATTTTTGGCGTACTCACCAGTCGCC
GCCCCTCGCCTCTTGCCGTGCGCGCTTCAGCAAGCCGAGTCCTGCGTCGAGAGAGCTCCTCTGGTTTCCC
TTTCGCTTTCAAGTCCCTGTTCGGGCGCCACTGCTAGAGATTTTCCACACTGACTAAAAGGGTCTGAGGG
ATCTCTAGTTACCAGAGTCACACAACAGACGGGCACACACTACTTGAAGCACTCAAGGCAAGCTTTATTG
AGGCTTAAGCAGTGGGTTCCCTAGTTAGCCAGAGAGCTCCCAGGCTCAGATCTGGTCTAACCAGAGAGA
CCCAGTACAGGCAAAAAGCAGCTGCTTATATGCAGGATCTGAGGGCTCGCCACTCCCCAGTCCCGCCCA
GGCCACGCCTCCCTGGAAAGTCCCCAGCGGAAAGTCCCTTGTAGCAAGCTCGATATCAGCAGTTCTTGAA
GTACTCCGGATGCAGCTCTCGGGCCACGTGATGAAATGCTAGGCGGCTGTCAAACCTCCACTCTAACACT
TCTCTCTCCGGGTCATCCATCCCATGCAGGCTCACAGGGTGTAACAAGCTGGTGTTCTCTCCTTTATTGGCC
TCTTCTACCTTATCTGGCTCAACTGGTACTAGCTTGTAGCACCATCCAAAGGTCAGTGGATATCTGACCCCT
GGCCCTGGTGTGTAGTTCTGCTAATCAGGGAAGTAGCCTTGTGTGTGGTAGATCCACAGATCAAGGATATC
TTGTCTTCTTTGGGAGTGAATTAGCCCTTCCAACTACTAAGTTTGTAGTACATATTTAACAAATACAATTTCT
TTAAAATGAAAATAATTCAGAGGAATCACAGGTTTAGAGTAAATGAAACCACAGGTAATTGGCAGTGGTA
ATAGGGTATGGGGTGGGAAGTTTGGGATGATTTTGGTTAGCTTGAGTTATCCAGTTGATCCAGACATGATA
AGATACATTGATGAGTTTGGACAAACCACAACTAGAATGCAGTGAAAAAAATGCTTTATTTGTGAAATTCG
TAATCATGTCATAGCTGTTTCCTGTGTGAAATTGTTATCCGCTCACAATTCCACACAACATACGAGCCGGAA
GCATAAAGTGTAAAGCCTGGGGTGCCTAATGAGTGAGCTAACTCACATTAATTGCGTTGCGCTCACTGCCC
GCTTTCCAGTCGGGAAACCTGTCGTGCCACGGGGAGCTTTTTGCAAAAGCCTAGGCCTCCAAAAAAGCC

TCCTCACTACTTCTGGAATAGCTCAGAGGCCGAGGCGGCCTCGGCCTCTGCATAAATAAAAAAAATTAGT

CAGCCATGAGCTTGCTGCATTAATGAATCGGCCAACGCGCGGGGAGAGGCGGTTTGCGTATTGGGCGCT

CTTCCGCTTCCTCGCTCACTGACTCGCTGCGCTCGGTCGTTCGGCTGCGGCGAGCGGTATCAGCTCACTCA

AAGGCGGTAATACGGTTATCCACAGAATCAGGGGATAACGCAGGAAAGAACATGTGAGCAAAAGGCCA

GCAAAAGGCCAGGAACCGTAAAAAGGCCGCGTTGCTGGCGTTTTTCCATAGGCTCCGCCCCCCTGACGA

GCATCACAAAAATCGACGCTCAAGTCAGAGGTGGCGAAACCCGACAGGACTATAAAGATACCAGGCGTT

TCCCCCTGGAAGCTCCCTCGTGCGCTCTCCTGTTCCGACCCTGCCGCTTACCGGATACCTGTCCGCCTTTCT

CCCTTCGGGAAGCGTGGCGCTTTCTCATAGCTCACGCTGTAGGTATCTCAGTTCGGTGTAGGTCGTTCGCT

CCAAGCTGGGCTGTGTGCACGAACCCCCCGTTCAGCCCGACCGCTGCGCCTTATCCGGTAACTATCGTCTT

GAGTCCAACCCGGTAAGACACGACTTATCGCCACTGGCAGCAGCCACTGGTAACAGGATTAGCAGAGCG

AGGTATGTAGGCGGTGCTACAGAGTTCTTGAAGTGGTGGCCTAACTACGGCTACACTAGAAGAACAGTAT

TTGGTATCTGCGCTCTGCTGAAGCCAGTTACCTTCGGAAAAAGAGTTGGTAGCTCTTGATCCGGCAAACA

AACCACCGCTGGTAGCGGTGGTTTTTTTGTTTGCAAGCAGCAGATTACGCGCAGAAAAAAAGGATCTCA

AGAAGATCCTTTGATCTTTTCTACGGGGTCTGACGCTCAGTGGAACGAAAACTCACGTTAAGGGATTTTG

GTCATGAGATTATCAAAAAGGATCTTCACCTAGATCCTTTTAAATTAAAAATGAAGTTTTAAATCAATCTAA

AGTATATATGAGTAAACTTGGTCTGACAGTTACCAATGCTTAATCAGTGAGGCACCTATCTCAGCGATCTGT

CTATTTCGTTCATCCATAGTTGCCTGACTCCCCGTCGTGTAGATAACTACGATACGGGAGGGCTTACCATCT

GGCCCCAGTGCTGCAATGATACCGCGAGACCCACGCTCACCGGCTCCAGATTTATCAGCAATAAACCAGC

CAGCCGGAAGGGCCGAGCGCAGAAGTGGTCCTGCAACTTTATCCGCCTCCATCCAGTCTATTAATTGTTG

CCGGGAAGCTAGAGTAAGTAGTTCGCCAGTTAATAGTTTGCGCAACGTTGTTGCCATTGCTACAGGCATC

GTGGTGTCACGCTCGTCGTTTGGTATGGCTTCATTCAGCTCCGGTTCCCAACGATCAAGGCGAGTTACATG

ATCCCCCATGTTGTGCAAAAAAGCGGTTAGCTCCTTCGGTCCTCCGATCGTTGTCAGAAGTAAGTTGGCCG

CAGTGTTATCACTCATGGTTATGGCAGCACTGCATAATTCTCTTACTGTCATGCCATCCGTAAGATGCTTTTC

TGTGACTGGTGAGTACTCAACCAAGTCATTCTGAGAATAGTGTATGCGGCGACCGAGTTGCTCTTGCCCGG

CGTCAACACGGGATAATACCGCGCCACATAGCAGAACTTTAAAAGTGCTCATCATTGGAAAACGTTCTTC

GGGGCGAAAACTCTCAAGGATCTTACCGCTGTTGAGATCCAGTTCGATGTAACCCACTCGTGCACCCAAC

TGATCTTCAGCATCTTTTACTTTCACCAGCGTTTCTGGGTGAGCAAAAACAGGAAGGCAAAATGCCGCAA

AAAAGGGAATAAGGGCGACACGGAAATGTTGAATACTCAT

> RTEL1_lenti_vector

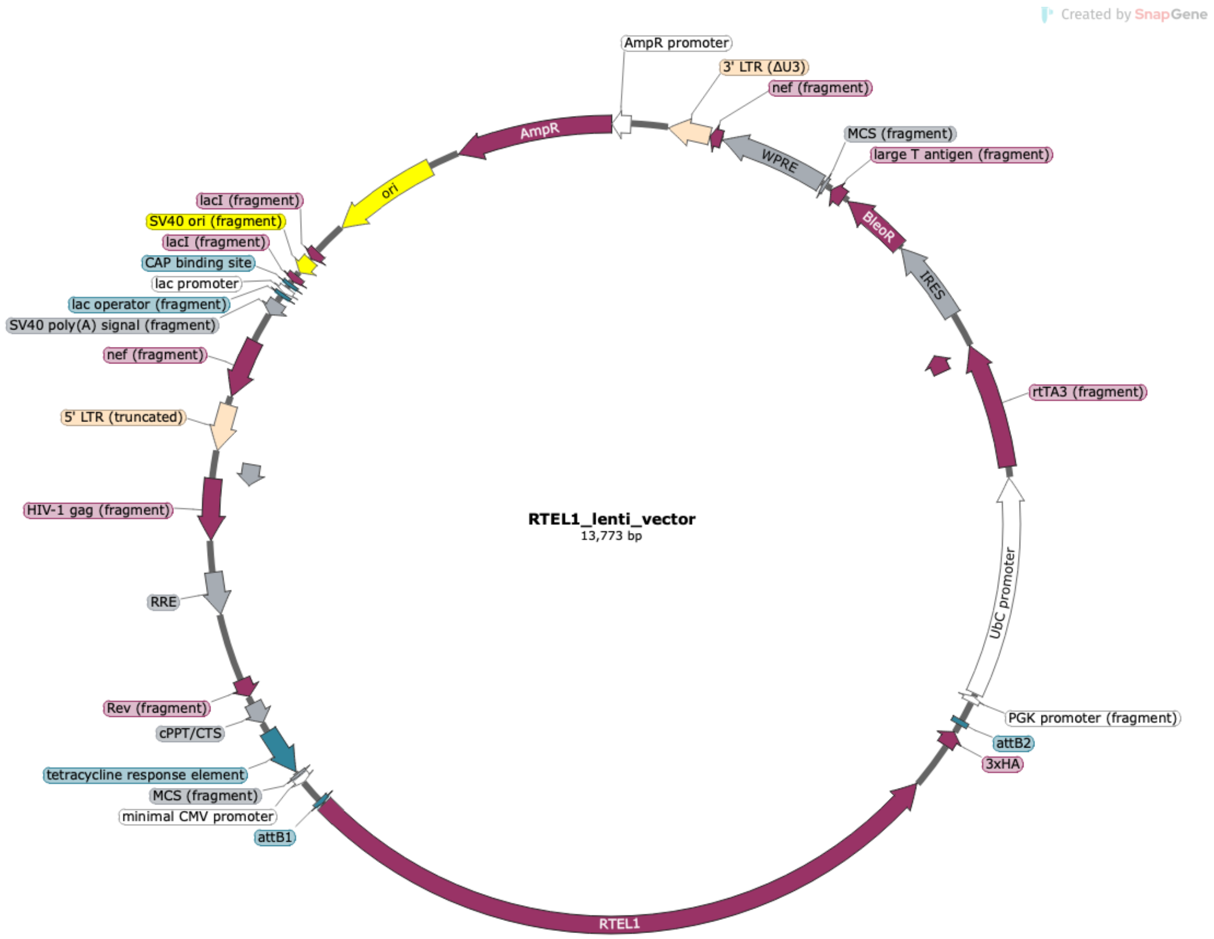


CATACTCTTCCTTTTTCAATATTATTGAAGCATTTATCAGGGTTATTGTCTCATGAGCGGATACATATTTGAAT

GTATTTAGAAAAATAAACAAATAGGGGTTCCGCGCACATTTCCCCGAAAAGTGCCACCTGACGTCTAAGA

AACCATTATTATCATGACATTAACCTATAAAAATAGGCGTATCACGAGGCCCTTTCGTCTTCAAGAATGATC

TAGCCCTTTCCTTAATTAACCCGGGCCTCTCACTCTCTGATATTCATTTCTTTGCAAGTTATAAATACTGAAT

AATAAGATGACATGAACTACTACTGCTAGAGATTTTCCACACTGACTAAAAGGGTCTGAGGGATCTCTAGT

TACCAGAGTCACACAACAGACGGGCACACACTACTTGAAGCACTCAAGGCAAGCTTTATTGAGGCTTAA

GCAGTGGGTTCCCTAGTTAGCCAGAGAGCTCCCAGGCTCAGATCTGGTCTAACCAGAGAGACCCAGTAC

AAGCAAAAAGCAGATCTTGTCTTCGTTGGGAGTGAATTAGCCCTTCCAGTCCCCCCTTTTCTTTTAAAAAG

TGGCTAAGATCTACAGCTGCCTTGTAAGTCATTGGTCTTAAAGGATCTCAGGCGGGGAGGCGGCCCAAAG

GGAGATCCGACTCGTCTGAGGGCGAAGGCGAAGACGCGGAAGAGGCCGCAGAGCCGGCAGCAGGCC

GCGGGAAGGAAGGTCCGCTGGATTGAGGGCCGAAGGGACGTAGCAGAAGGACGTCCCGCGCAGAATC

CAGGTGGCAACACAGGCGAGCAGCCAAGGAAAGGACGATGATTTCCCCGACAACACCACGGAATTGTC
AGTGCCCAACAGCCGAGCCCCTGTCCAGCAGCGGGCAAGGCAGGCGGCGATGAGTTCCGCCGTGGCAA
TAGGGAGGGGGAAAGCGAAAGTCCCGGAAAGGAGCTGACAGGTGGTGGCAATGCCCCAACCAGTGGG
GGTTGCGTCAGCAAACACAGTGCACACCACGCCACGTTGCCTGACAACGGGCCACAACTCCTCATAAAG
AGACAGCAACCAGGATTTATACAAGGAGGAGAAAATGAAAGCCATACGGGAAGCAATAGCATGATACA
AAGGCATTAAAGCAGCGTATCCACATAGCGTAAAAGGAGCAACATAGTTAAGAATACCAGTCAATCTTTC
ACAAATTTTGTAATCCAGAGGTTGATTAGGATCTATCGCGATAAGCTTGATATCGAATTGGGAGGGACTAA
ACAACAACAATTGCATTCATTTTATGTTTCAGGTTCAGGGGGAGGTGTGGGAGGTTTTTTAAAGCAAGTA
AAACCTCTACAAATGTGGTATGGCTGATTATGATCTAGAGTCGCGGCCGCTTTACTTGTACATCAGTCCTGC
TCCTCGGCCACGAAGTGCACGCAGTTGCCGGCCGGGTCGCGCAGGGCGAACTCCCGCCCCCACGGCTG
CTCGCCGATCTCGGTCATGGCCGGCCCGGAGGCGTCCCGGAAGTTCGTGGACACGACCTCCGACCACTC
GGCGTACAGCTCGTCCAGGCCGCGCACCCACACCCAGGCCAGGGTGTTGTCCGGCACCACCTGGTCCTG
GACCGCGCTGATGAACAGGGTCACGTCGTCCCGGACCACACCGGCGAAGTCGTCCTCCACGAAGTCCCG
GGAGAACCCGAGCCGGTCGGTCCAGAACTCGACCGCTCCGGCGACGTCGCGCGCGGTGAGCACCGGAA
CGGCACTGGTCAACTTGGCCATGGTGGACCGGTAAGCTTATCATCGTGTTTTTCAAAGGAAAACCACGTC
CCCGTGGTTCGGGGGGCCTAGACGTTTTTTTAACCTCGACTAAACACATGTAAAGCATGTGCACCGAGGC
CCCAGATCAGATCCCATACAATGGGGTACCTTCTGGGCATCCTTCAGCCCCTTGTTGAATACGCTTGAGGA
GAGCCATTTGACTCTTTCCACAACTATCCAACTCACAACGTGGCACTGGGGTTGTGCCGCCTTTGCAGGT
GTATCTTATACACGTGGCTTTTGGCCGCAGAGGCACCTGTCGCCAGGTGGGGGGTTCCGCTGCCTGCAAA
GGGTCGCTACAGACGTTGTTTGTCTTCAAGAAGCTTCCAGAGGAACTGCTTCCTTCACGACATTCAACAG
ACCTTGCATTCCTTTGGCGAGAGGGGAAAGACCCCTAGGAATGCTCGTCAAGAAGACAGGGCCAGGTTT
CCGGGCCCTCACATTGCCAAAAGACGGCAATATGGTGGAAAATAACATATAGACAAACGCACACCGGCC
TTATTCCAAGCGGCTTCGGCCAGTAACGTTAGGGGGGGGAGAGGGGCGGAATTCCTCTAGTGCGGCCGC
GGATCCTTACTTAGTTACCCGGGGAGCATGTCAAGGTCAAAATCGTCAAGAGCGTCAGCAGGCAGCATAT
CAAGGTCAAAGTCGTCAAGGGCATCGGCTGGGAGCATGTCTAAGTCAAAATCGTCAAGGGCGTCGGTCG
GCCCGCCGCTTTCGCACTTTAGCTGTTTCTCCAGGCCACATATGATTAGTTCCAGGCCGAAAAGGAAGGC
AGGTTCGGCTCCCTGCCGGTCGAACAGCTCAATTGCTTGTCTCAGAAGTGGGGGCATAGAATCGGTGGTA
GGTGTCTCTCTTTCCTCTTTTGCTACTTGATGCTCCTGTTCCTCCAATACGCAGCCCAGTGTAAAGTGGCCC
ACGGCGGACAGAGCGTACAGTGCGTTCTCCAGGGAGAAGCCTTGCTGACACAGGAACGCGAGCTGATT

TTCCAGGGTTTCGTACTGTTTCTCTGTTGGGCGGGTGCCGAGATGCACTTTAGCCCCGTCGCGATGTGAGA
GGAGAGCACAGCGGTATGACTTGGCGTTGTTCCGCAGAAAGTCTTGCCATGACTCGCCTTCCAGGGGGC
AGAAGTGGGTATGATGCCTGTCCAGCATCTCGATTGGCAGGGCATCGAGCAGGGCCCGCTTGTTCTTCAC
GTGCCAGTACAGGGTAGGCTGCTCAACTCCCAGCTTTTGAGCGAGTTTCCTTGTCGTCAGGCCTTCGATAC
CGACACCATTGAGTAATTCCAGAGCTCCGTTTATGACTTTGCTCTTGTCCAGTCTAGACATGGTGAATTCGA
CTGCAGGACCGGTACGGGGTGGAGATCCGAGCTCGGTACCAAGCTTCGTCTAACAAAAAAGCCAAAAAC
GGCCAGAATTTAGCGGACAATTTACTAGTCTAACACTGAAAATTACATATTGACCCAAATGATTACATTTCA
AAAGGTGCCTAAAAAACTTCACAAAACACACTCGCCAACCCCGAGCGCATAGTTCAAAACCGGAGCTTC
AGCTACTTAAGAAGATAGGTACATAAAACCGACCAAAGAAACTGACGCCTCACTTATCCCTCCCCTCACC
AGAGGTCCGGCGCCTGTCGATTCAGGAGAGCCTACCCTAGGCCCGAACCCTGCGTCCTGCGACGGAGAA
AAGCCTACCGCACACCTACCGGCAGGTGGCCCCACCCTGCATTATAAGCCAACAGAACGGGTGACGTCA
CGACACGACGAGGGCGCGCGCTCCCAAAGGTACGGGTGCACTGCCCAACGGCACCGCCATAACTGCCG
CCCCCGCAACAGACGACAAACCGAGTTCTCCAGTCAGTGACAAACTTCACGTCAGGGTCCCCAGATGGT
GCCCCAGCCCATCTCACCCGAATAAGAGCTTTCCCGCATTAGCGAAGGCCTCAAGACCTTGGGTTCTTGC
CGCCCACCATGCCCCCCACCTTGTTTCAACGACCTCACAGCCCGCCTCACAAGCGTCTTCCATTCAAGACT
CGGGAACAGCCGCCATTTTGCTGCGCTCCCCCCAACCCCCAGTTCAGGGCAACCTTGCTCGCGGACCCA
GACTACAGCCCTTGGCGGTCTCTCCACACGCTTCCGTCCCACCGAGCGGCCCGGCGGCCACGAAAGCCC
CGGCCAGCCCAGCAGCCCGCTACTCACCAAGTGACGATCACAGCGATCCACAAACAAGAACCGCGACC
CAAATCCCGGCTGCGACGGAACTAGCTGTGCCACACCCGGCGCGTCCTTATATAATCATCGGCGTTCACC
GCCCCACGGAGATCCCTCCGCAGAATCGCCGAGAAGGGACTACTTTTCCTCGCCTGTTCCGCTCTCTGGA
AAGAAAACCAGTGCCCTAGAGTCACCCAAGTCCCGTCCTAAAATGTCCTTCTGCTGATACTGGGGTTCTAA
GGCCGAGTCTTATGAGCAGCGGGCCGCTGTCCTGAGCGTCCGGGCGGAAGGATCAGGACGCTCGCTGC
GCCCTTCGTCTGACGTGGCAGCGCTCGCCGTGAGGAGGGGGGCGCCCGCGGGAGGCGCCAAAACCCG
GCGCGGAGGCCAGATCTTGGGTGGGTTACTCCAGACTGCCTTGGGAAAAGCGCCTCCCCTACCCGGTAG
AATTTCTAGTTTAATTAATCATTACTAAGCGTAGTCTGGGACGTCGTATGGGTATTCGAACCGCGGGCCCTC
TAGACTCGAGCGGCCGCCACTGTGCTGGATATCAACCACTTTGTACAAGAAAGCTGGGTCTAGATATCTCG
AGGCGGCCGCTTATCCGGATTCGAATCATTAGGATCCAGCGTAATCTGGAACGTCATAAGGATACGATCCT
GCATAGTCCGGGACGTCATAGGGATAGCCCGCATAGTCAGGAACATCGTATGGGTACCCGCCGGTGCCCT
GGGGCTCTGGCCAGAAGACCTGCATGACGCTCTGCTTCCTGGAGGCGGTGTGGCAGGCTGGGCACATCC

TAGAGGCCTGAAGGTGCCGTTGCCAGCAGGCTTGGCAGCGCTGGAAGTCACAGGCAGGGCACTGGAA
GGGCACCACGTCCTCTGCCCCACAGCCCTGGCACACACAGCCTGCTGAGAGGGGCCCGCCCGGAGCTC
CCGTGGCCTGCTGCCCAGCGATGTCTCTCCCATGAGGCTCACCCCACTCAGATGCTGCAGGCCCGTGGGG
AGGTCCTGAGGACTGGCTGGGACCTGCATCCTCACCGCCCGCCCCCACAGTCCCTGCTGGCCTCTGTCTA
AGGAAGGACGAGATCTTGCTCTGGGTCTTCCCGGTCTTCTCGGACCGTGAGGGGCCTGGTTGGGGAGCC
CTGTGGGTAAGCACAGGAGGCACGGCAAGCCTCTCCTCCTGGGGTCCCGGTGGCTCCATGCCCGGGTAG
GGCCGGCCGGTCAGGTCTGTGCACGTCTGTGAGAAGCGCTGCTTGTGGTGTGGACGCACAAACATGCTG
AACCTGTGCAGCAGGGGGAAGTCCTCTGGCTTTGCAGTGGTCAGGGCGGCCAACACAGCCAGCACCTT
GTCGAGGTCGTCGTCTTGCTTATAGGCTGTCAGCGCTGCCAAGAGTTGGCTACAGCCCGCGGACCCCAGG
GCCCTGCGGGCATCAGCCAGGTAGGCGCTCACGGCGTGCTGGCCCTGCTTCCCTGCTCTGGGCACTCCA
GACCCCCACTGTGGTTGGCTGCCAGGGTCTCCTGTTGGGGGTGGCCTGGGCGACAGGTGGGGCCTGCCC
TGGTTCAGGTGCTCTTGGGGGTCCAGCTGCTGGGCTGCAGCCGTGGACACGGTCAGCTTGGGATCCGGC
GCCGTTCTTCCAGTGGGGTCCAGGACCGGCTGTGCCCGCTGCCTTCGGGGAATGCTGTGCTCAGGCCGAT
AGCCACAGCCTCGTCCTGTCAGCTGGATACAGACCTCCTCAAACTGCTGCTTATGGTGGGGCCGCACAAA
CTGGTAGAAGCCTTGGAGCAGGTTGTGCTTCTTGGGGTCCTCAGCAAAGAGGGGGCCGAGACAGGCGG
CCAGGGCGGCGAAGTCATCGGAACCCTTGTAGTCCTGCAGGGCCTGGGTGAAGGTGGCAAAGTTGGCTT
GGCTCAACTCCTGCTTCACGGCCACCATGAAGAGCTTGGCCCTGTCCGTCTGTGCACCAGCCACGGGCTC
CTCCGGGTGGCTGACCAGCCGGATCTTCTTCCTCCCTCCTCGCGGTTCTTCTGCCGGCCTCTTCTCAGACA
GGAGGGACAGGGTGGAGCAGCTGTGGGCCTGCTCCTCGCCAGGGCTCCCCGCCCGCTGTTCGCTGTGCT
CCAGGGCGGCCAGCAGCCCCCTGGGCCTCTGCCGGGCAGGAACTGGCTCCTGCTCATACTCCACACACA
GGCTACTCTCGGGGTCCCCGGCAGCTGGTGACCCTGAGGACCTCTGCTTCAGGCTGGGGACATGCAGGT
CCAGACTCTTAGCTTTCCTGGTGGAGAAGAAGGGGCCAGGCGACTTGGCCTCGCTGACAGCATCTTCTC
CACGCACACTGGGTGCTGTAGCCCGGGGGGCCGGCGCTGGCATAGTTCGCTCGGCAACACGGAAGAAC
TGGGCCACGTCTCGGATGACATGGCCAAAGTTGTCATACACCCTGACGTGGGGACGCACCCAGGAGGGC
AGTTGGGCTCTTGCGTCGGCAAAGGCGAACCTGTGGTCACAGAGGAAGACAGCTCCGTAGTCCTGGCGG
TGCCGGATCACTCGCCCGATGGCCTGGTTCACAGCCCTGGACGCCTGCTGCCGGTACCACTCCTGCCCAG
AGAGGAACTGGCCCCCAGCCCCACCCTGGCCCTTCATCTCATCCAGGAACTGCATCTTGAGGACAACCCG
GGGGTCCATGCGTGGGGGGTACGGGAGGCCCGTGACAATCACACCACGGCCATTCGTGTCTGAGAAGTC
CAGCCCCTCGCTGGCCTTGCCCCGGCAGACCGCCAGGAAGGTGGCGCCGGTGGACCCAGGGGCGGCA

ACCCTTGCATAGTAAGCACTGATGGTCTCGGAGAAGCTGCCTTTGCTCCTGGGCTCCACAAACAGCGGCT
TCAGCGCCTCCATCTTCCTGGCCAAGTCGCGGGCCCGCCAGAACTCCAGGCTCTTCTCCATGACAGGATA
GGAAGGGAAGAAGATCAGGAGCCCATAGGGCACCACGCGGGCGATGTTGCCCAGAGCCTTCCCCAGG
GAGGATAAGCACTCCTCGGAAAACCGTCTGTCAAACGCGGAGCTCAACTGGGCTCCATCGGGGCCTCTG
GGGACGACCCCCACCCAGATCTGGTGCTTGTCGATGATGTGTGGGTTCTCCAGGCAGACTGGGAAAGGG
ATCTGCATCTCCAGAGCAAAGGAGGACACCGGGGCCAGCGTGCCGCTGGTAAGGATGAGGGAGCGGAC
GCCCTGGCGGACCAGCTCGTGCATGCTGTGGCCGGGACTGAAGCACCAGTAGCTCAGCACCTTCCCTCG
CTTTCTGGCTGCAGTGGTGCTCCAGGCATCAGACCGCTGAGCCGTCCTCCGGTGACCAGCATCAGGATGG
ATGTGCACCTTATAGGACTGTAAGGCCCCCAGCCCTGCTGGGGAACCAGGGCTGCCCTCGGAGGGGTCC
ACACTGAACACAATCTGGATAATGTCCGCCAGCTTCTGCAGTCCGGCCGTGTTGGTGAACACTCCAGCAC
GTCCTGCCAGGTGCTGGATGATCTGGTCCAGCGAGTCCAGGATGCAGCCCTTGGTCTGAAACGTGATCTG
GGCTTCAGCAAACAGCTCAAAGATGTAGCTCCCTGGCTTGGTGACACCGCTGTCGTCTCCAGGCAGCTCA
ACAGCATCGATGGCCCCCTCCAGGCGCAGCAGGATCATCTTCAGCTTTGCAATGTCTTCCAGCTCCATGTT
CAGCCCTGGGCTGGGGGAGTCCGCGCTGAACTCCGGGTGGGGCTCACCCTGCTGCGCTGCCTTGGTCTG
CTCCTCCAGCACCTGGTCTATGACGTCCAGTCCTGAAGCCAGGTCATGGGGAGTCAGGTCAAAGGATGCC
GATTCTTCACACATCTTCTCCACGTTGTGAGCTTCGTCAAAGATCACGACTGTCCCCTTCAGGTCAATGTTG
TGTGCTCTGCGGCTCTTGGCATCCAACAAGTAATTGTACGGCATGAATATGATGTCGGCTTGCTGCTTCAG
GTTCCGGGACAGGTAGTAAGGGCACACCCTGTGCTTGCTTCCGCTCTTGACCAAGTCCTCAATGTCCAGG
ATGGGGCTGGCCAGCTCCTGCTCCAGGCTTTTTTCTTCTACGTTGTTGTAGAAATGACAGGAGCGACTTGC
CACCTTCTTACGGCACAAGTGGATCTGTAGATGGTTACTCTCTTGTTTCTTCACCTCAGGATGGATGCACAG
CTGCTCCCGGGAGCCCAGCACACACACCTTAGGCCGGTAGGAGGTGTTCCGAAGCTCGTTGATGACCTG
TGTGAGTTGCGAGTGGGTCCTGGAGGCGTAAATAATCTTTGGGATGTCCGTGTAGCAAGCTATGGGGTCTC
CAGCAGCAGCAGCAGCGTTGCCCCAGGATGACAAGGCCCGATCCGGGAAAAGCTCTCCTTGCGCCCTCT
CGGCAATCTTGCGGGCAGAGATGCCGTCTCGGAGGTGTTCTCGCCAGGCCAGCGTGGTGCACAGCAGGC
ACAGCGTCTTCCCTGTACCCGTAGGGCTCTCCAGGATGCCATTCACCTTCTGCTGCAGACATTCCAGGACC
TTGGTCATGTACTCCTGTTGGCATTTGTAGGGCTGGAAAGGGAAGTCTACGGTCACACCATTCAGGACTAT
CTTGGGCATGGTGGCGCCAGCCTGCTTTTTTGTACAAACTTGTTGATATCTGCAGAATTCCACCACACTGG
ACTAGTTCGGGGCCGCGGAGGCTGGATCGGTCCCGGTGTCTTCTATGGAGGTCAAAACAGCGTGGATGG
CGTCTCCAGGCGATCTGACGGTTCACTAAACGAGCTCTGCTTATATAGGCCTCCCACCGTACACGCCTACC

TCGACCCGGGTACCGAGCTCGACTTTCACTTTTCTCTATCACTGATAGGGAGTGGTAAACTCGACTTTCAC
TTTTCTCTATCACTGATAGGGAGTGGTAAACTCGACTTTCACTTTTCTCTATCACTGATAGGGAGTGGTAAA
CTCGACTTTCACTTTTCTCTATCACTGATAGGGAGTGGTAAACTCGACTTTCACTTTTCTCTATCACTGATAG
GGAGTGGTAAACTCGACTTTCACTTTTCTCTATCACTGATAGGGAGTGGTAAACTCGACTTTCACTTTTCTC
TATCACTGATAGGGAGTGGTAAAGGATCCTAGTCCAAACTGGATCTCTGCTGTCCCTGTAATAAACCCGAA
AATTTTGAATTTTTGTAATTTGTTTTTGTAATTCTTTAGTTTGTATGTCTGTTGCTATTATGTCTACTATTCTTTC
CCCTGCACTGTACCCCCCAATCCCCCCTTTTCTTTTAAAATTGTGGATGAATACTGCCATTTGTGAATTCGG
CGATACCGTCGAGATCCGTTCACTAATCGAATGGATCTGTCTCTGTCTCTCTCTCCACCTTCTTCTTCTATTCC
TTCGGGCCTGTCGGGTCCCCTCGGGGTTGGGAGGTGGGTCTGAAACGATAATGGTGAATATCCCTGCCTA
ACTCTATTCACTATAGAAAGTACAGCAAAAACTATTCTTAAACCTACCAAGCCTCCTACTATCATTATGAATA
ATTTTATATACCACAGCCAATTTGTTATGTTAAACCAATTCCACAAACTTGCCCATTTATCTAATTCCAATAAT
TCTTGTTCATTCTTTTCTTGCTGGTTTTGCGATTCTTCAATTAAGGAGTGTATTAAGCTTGTGTAATTGTTAAT
TTCTCTGTCCCACTCCATCCAGGTCGTGTGATTCCAAATCTGTTCCAGAGATTTATTACTCCAACTAGCATTC
CAAGGCACAGCAGTGGTGCAAATGAGTTTTCCAGAGCAACCCCAAATCCCCAGGAGCTGTTGATCCTTTA
GGTATCTTTCCACAGCCAGGATTCTTGCCTGGAGCTGCTTGATGCCCCAGACTGTGAGTTGCAACAGATG
CTGTTGCGCCTCAATAGCCCTCAGCAAATTGTTCTGCTGCTGCACTATACCAGACAATAATTGTCTGGCCTG
TACCGTCAGCGTCATTGACGCTGCGCCCATAGTGCTTCCTGCTGCTCCCAAGAACCCAAGGAACAAAGCT
CCTATTCCCACTGCTCTTTTTTCTCTCTGCACCACTCTTCTCTTTGCCTTGGTGGGTGCTACTCCTAATGGTT
CAATTTTTACTACTTTATATTTATATAATTCACTTCTCCAATTGTCCCTCATATCTCCTCCTCCAGGTCTGAAGA
TCAGCGGCCGGCCGCTTGCTGTGCGGTGGTCTTACTTTTGTTTTGCTCTTCCTCTATCTTGTCTAAAGCTTC
CTTGGTGTCTTTTATCTCTATCCTTTGATGCACACAATAGAGGGTTGCTACTGTATTATATAATGATCTAAGTT
CTTCTGATCCTGTCTGAAGGGATGGTTGTAGCTGTCCCAGTATTTGTCTACAGCCTTCTGATGTTTCTAACAG
GCCAGGATTAACTGCGAATCGTTCTAGCTCCCTGCTTGCCCATACTATATGTTTTAATTTATATTTTTTCTTTC
CCCCTGGCCTTAACCGAATTTTTTCCCATCGCGATCTAATTCTCCCCCGCTTAATACTGACGCTCTCGCACC
CATCTCTCTCCTTCTAGCCTCCGCTAGTCAAAATTTTTGGCGTACTCACCAGTCGCCGCCCCTCGCCTCTTG
CCGTGCGCGCTTCAGCAAGCCGAGTCCTGCGTCGAGAGAGCTCCTCTGGTTTCCCTTTCGCTTTCAAGTCC
CTGTTCGGGCGCCACTGCTAGAGATTTTCCACACTGACTAAAAGGGTCTGAGGGATCTCTAGTTACCAGA
GTCACACAACAGACGGGCACACACTACTTGAAGCACTCAAGGCAAGCTTTATTGAGGCTTAAGCAGTGG
GTTCCCTAGTTAGCCAGAGAGCTCCCAGGCTCAGATCTGGTCTAACCAGAGAGACCCAGTACAGGCAAA

AAGCAGCTGCTTATATGCAGGATCTGAGGGCTCGCCACTCCCCAGTCCCGCCCAGGCCACGCCTCCCTGG
AAAGTCCCCAGCGGAAAGTCCCTTGTAGCAAGCTCGATATCAGCAGTTCTTGAAGTACTCCGGATGCAGC
TCTCGGGCCACGTGATGAAATGCTAGGCGGCTGTCAAACCTCCACTCTAACACTTCTCTCTCCGGGTCATC
CATCCCATGCAGGCTCACAGGGTGTAACAAGCTGGTGTTCTCTCCTTTATTGGCCTCTTCTACCTTATCTGG
CTCAACTGGTACTAGCTTGTAGCACCATCCAAAGGTCAGTGGATATCTGACCCCTGGCCCTGGTGTGTAGT
TCTGCTAATCAGGGAAGTAGCCTTGTGTGTGGTAGATCCACAGATCAAGGATATCTTGTCTTCTTTGGGAGT
GAATTAGCCCTTCCAACTACTAAGTTTGTAGTACATATTTAACAAATACAATTTCTTTAAAATGAAAATAATT
CAGAGGAATCACAGGTTTAGAGTAAATGAAACCACAGGTAATTGGCAGTGGTAATAGGGTATGGGGTGG
GAAGTTTGGGATGATTTTGGTTAGCTTGAGTTATCCAGTTGATCCAGACATGATAAGATACATTGATGAGTT
TGGACAAACCACAACTAGAATGCAGTGAAAAAAATGCTTTATTTGTGAAATTCGTAATCATGTCATAGCTG
TTTCCTGTGTGAAATTGTTATCCGCTCACAATTCCACACAACATACGAGCCGGAAGCATAAAGTGTAAAGC
CTGGGGTGCCTAATGAGTGAGCTAACTCACATTAATTGCGTTGCGCTCACTGCCCGCTTTCCAGTCGGGAA
ACCTGTCGTGCCACGGGGAGCTTTTTGCAAAAGCCTAGGCCTCCAAAAAAGCCTCCTCACTACTTCTGGA
ATAGCTCAGAGGCCGAGGCGGCCTCGGCCTCTGCATAAATAAAAAAAATTAGTCAGCCATGAGCTTGCTG
CATTAATGAATCGGCCAACGCGCGGGGAGAGGCGGTTTGCGTATTGGGCGCTCTTCCGCTTCCTCGCTCA
CTGACTCGCTGCGCTCGGTCGTTCGGCTGCGGCGAGCGGTATCAGCTCACTCAAAGGCGGTAATACGGTT
ATCCACAGAATCAGGGGATAACGCAGGAAAGAACATGTGAGCAAAAGGCCAGCAAAAGGCCAGGAAC
CGTAAAAAGGCCGCGTTGCTGGCGTTTTTCCATAGGCTCCGCCCCCCTGACGAGCATCACAAAAATCGAC
GCTCAAGTCAGAGGTGGCGAAACCCGACAGGACTATAAAGATACCAGGCGTTTCCCCCTGGAAGCTCCC
TCGTGCGCTCTCCTGTTCCGACCCTGCCGCTTACCGGATACCTGTCCGCCTTTCTCCCTTCGGGAAGCGTG
GCGCTTTCTCATAGCTCACGCTGTAGGTATCTCAGTTCGGTGTAGGTCGTTCGCTCCAAGCTGGGCTGTGT
GCACGAACCCCCCGTTCAGCCCGACCGCTGCGCCTTATCCGGTAACTATCGTCTTGAGTCCAACCCGGTAA
GACACGACTTATCGCCACTGGCAGCAGCCACTGGTAACAGGATTAGCAGAGCGAGGTATGTAGGCGGTG
CTACAGAGTTCTTGAAGTGGTGGCCTAACTACGGCTACACTAGAAGAACAGTATTTGGTATCTGCGCTCTG
CTGAAGCCAGTTACCTTCGGAAAAAGAGTTGGTAGCTCTTGATCCGGCAAACAAACCACCGCTGGTAGC
GGTGGTTTTTTTGTTTGCAAGCAGCAGATTACGCGCAGAAAAAAAGGATCTCAAGAAGATCCTTTGATCT
TTTCTACGGGGTCTGACGCTCAGTGGAACGAAAACTCACGTTAAGGGATTTTGGTCATGAGATTATCAAA
AAGGATCTTCACCTAGATCCTTTTAAATTAAAAATGAAGTTTTAAATCAATCTAAAGTATATATGAGTAAACT
TGGTCTGACAGTTACCAATGCTTAATCAGTGAGGCACCTATCTCAGCGATCTGTCTATTTCGTTCATCCATAG

TTGCCTGACTCCCCGTCGTGTAGATAACTACGATACGGGAGGGCTTACCATCTGGCCCCAGTGCTGCAATG
ATACCGCGAGACCCACGCTCACCGGCTCCAGATTTATCAGCAATAAACCAGCCAGCCGGAAGGGCCGAG
CGCAGAAGTGGTCCTGCAACTTTATCCGCCTCCATCCAGTCTATTAATTGTTGCCGGGAAGCTAGAGTAAG
TAGTTCGCCAGTTAATAGTTTGCGCAACGTTGTTGCCATTGCTACAGGCATCGTGGTGTCACGCTCGTCGTT
TGGTATGGCTTCATTCAGCTCCGGTTCCCAACGATCAAGGCGAGTTACATGATCCCCCATGTTGTGCAAAA
AAGCGGTTAGCTCCTTCGGTCCTCCGATCGTTGTCAGAAGTAAGTTGGCCGCAGTGTTATCACTCATGGTT
ATGGCAGCACTGCATAATTCTCTTACTGTCATGCCATCCGTAAGATGCTTTTCTGTGACTGGTGAGTACTCA
ACCAAGTCATTCTGAGAATAGTGTATGCGGCGACCGAGTTGCTCTTGCCCGGCGTCAACACGGGATAATA
CCGCGCCACATAGCAGAACTTTAAAAGTGCTCATCATTGGAAAACGTTCTTCGGGGCGAAAACTCTCAAG
GATCTTACCGCTGTTGAGATCCAGTTCGATGTAACCCACTCGTGCACCCAACTGATCTTCAGCATCTTTTAC
TTTCACCAGCGTTTCTGGGTGAGCAAAAACAGGAAGGCAAAATGCCGCAAAAAAGGGAATAAGGGCGA
CACGGAAATGTTGAATACT

> TERT_lenti_vector

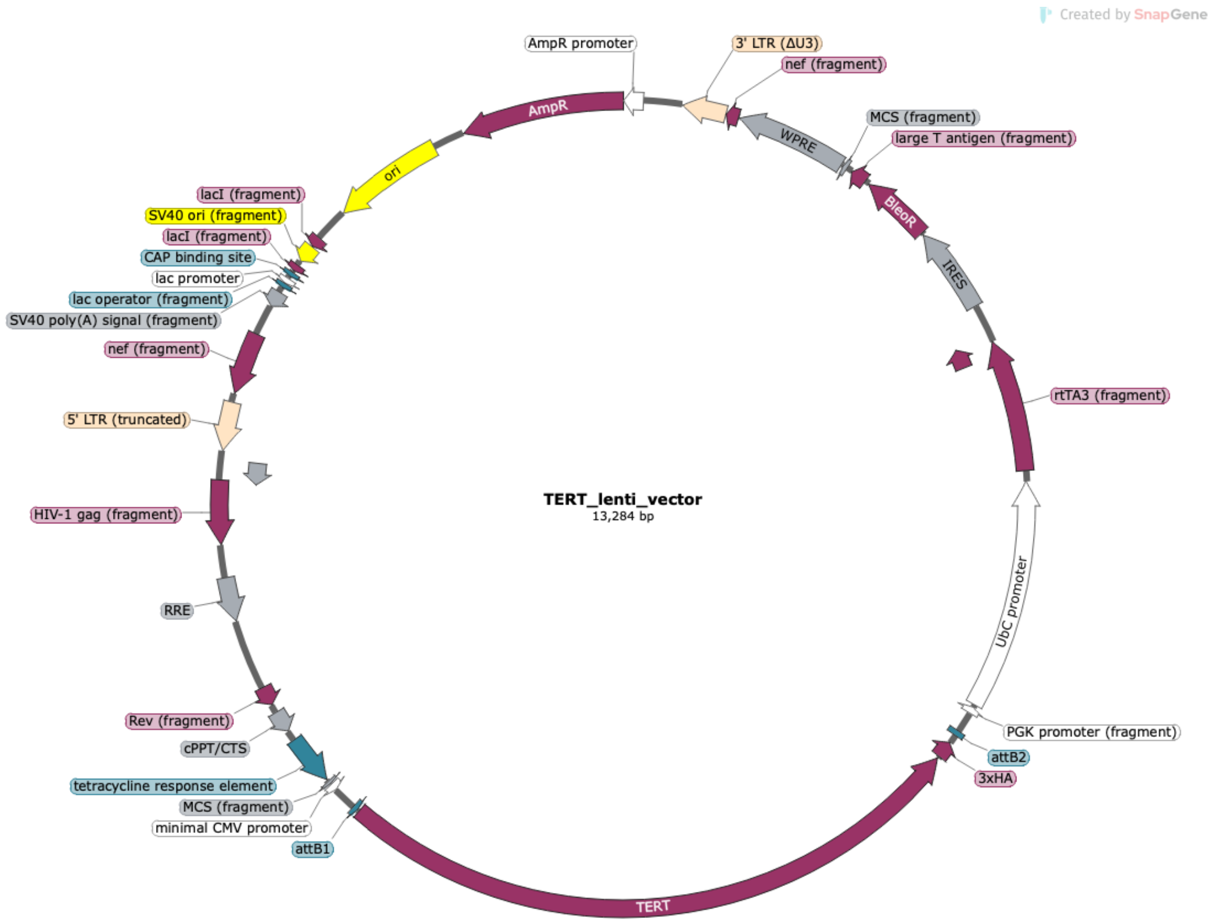


CATACTCTTCCTTTTTCAATATTATTGAAGCATTTATCAGGGTTATTGTCTCATGAGCGGATACATATTTGAAT
GTATTTAGAAAAATAAACAAATAGGGGTTCCGCGCACATTTCCCCGAAAAGTGCCACCTGACGTCTAAGA
AACCATTATTATCATGACATTAACCTATAAAAATAGGCGTATCACGAGGCCCTTTCGTCTTCAAGAATGATC
TAGCCCTTTCCTTAATTAACCCGGGCCTCTCACTCTCTGATATTCATTTCTTTGCAAGTTATAAATACTGAAT
AATAAGATGACATGAACTACTACTGCTAGAGATTTTCCACACTGACTAAAAGGGTCTGAGGGATCTCTAGT
TACCAGAGTCACACAACAGACGGGCACACACTACTTGAAGCACTCAAGGCAAGCTTTATTGAGGCTTAA
GCAGTGGGTTCCCTAGTTAGCCAGAGAGCTCCCAGGCTCAGATCTGGTCTAACCAGAGAGACCCAGTAC
AAGCAAAAAGCAGATCTTGTCTTCGTTGGGAGTGAATTAGCCCTTCCAGTCCCCCCTTTTCTTTTAAAAAG
TGGCTAAGATCTACAGCTGCCTTGTAAGTCATTGGTCTTAAAGGATCTCAGGCGGGGAGGCGGCCCAAAG
GGAGATCCGACTCGTCTGAGGGCGAAGGCGAAGACGCGGAAGAGGCCGCAGAGCCGGCAGCAGGCC
GCGGGAAGGAAGGTCCGCTGGATTGAGGGCCGAAGGGACGTAGCAGAAGGACGTCCCGCGCAGAATC

CAGGTGGCAACACAGGCGAGCAGCCAAGGAAAGGACGATGATTTCCCCGACAACACCACGGAATTGTC
AGTGCCCAACAGCCGAGCCCCTGTCCAGCAGCGGGCAAGGCAGGCGGCGATGAGTTCCGCCGTGGCAA
TAGGGAGGGGGAAAGCGAAAGTCCCGGAAAGGAGCTGACAGGTGGTGGCAATGCCCCAACCAGTGGG
GGTTGCGTCAGCAAACACAGTGCACACCACGCCACGTTGCCTGACAACGGGCCACAACTCCTCATAAAG
AGACAGCAACCAGGATTTATACAAGGAGGAGAAAATGAAAGCCATACGGGAAGCAATAGCATGATACA
AAGGCATTAAAGCAGCGTATCCACATAGCGTAAAAGGAGCAACATAGTTAAGAATACCAGTCAATCTTTC
ACAAATTTTGTAATCCAGAGGTTGATTAGGATCTATCGCGATAAGCTTGATATCGAATTGGGAGGGACTAA
ACAACAACAATTGCATTCATTTTATGTTTCAGGTTCAGGGGGAGGTGTGGGAGGTTTTTTAAAGCAAGTA
AAACCTCTACAAATGTGGTATGGCTGATTATGATCTAGAGTCGCGGCCGCTTTACTTGTACATCAGTCCTGC
TCCTCGGCCACGAAGTGCACGCAGTTGCCGGCCGGGTCGCGCAGGGCGAACTCCCGCCCCCACGGCTG
CTCGCCGATCTCGGTCATGGCCGGCCCGGAGGCGTCCCGGAAGTTCGTGGACACGACCTCCGACCACTC
GGCGTACAGCTCGTCCAGGCCGCGCACCCACACCCAGGCCAGGGTGTTGTCCGGCACCACCTGGTCCTG
GACCGCGCTGATGAACAGGGTCACGTCGTCCCGGACCACACCGGCGAAGTCGTCCTCCACGAAGTCCCG
GGAGAACCCGAGCCGGTCGGTCCAGAACTCGACCGCTCCGGCGACGTCGCGCGCGGTGAGCACCGGAA
CGGCACTGGTCAACTTGGCCATGGTGGACCGGTAAGCTTATCATCGTGTTTTTCAAAGGAAAACCACGTC
CCCGTGGTTCGGGGGGCCTAGACGTTTTTTTAACCTCGACTAAACACATGTAAAGCATGTGCACCGAGGC
CCCAGATCAGATCCCATACAATGGGGTACCTTCTGGGCATCCTTCAGCCCCTTGTTGAATACGCTTGAGGA
GAGCCATTTGACTCTTTCCACAACTATCCAACTCACAACGTGGCACTGGGGTTGTGCCGCCTTTGCAGGT
GTATCTTATACACGTGGCTTTTGGCCGCAGAGGCACCTGTCGCCAGGTGGGGGGTTCCGCTGCCTGCAAA
GGGTCGCTACAGACGTTGTTTGTCTTCAAGAAGCTTCCAGAGGAACTGCTTCCTTCACGACATTCAACAG
ACCTTGCATTCCTTTGGCGAGAGGGGAAAGACCCCTAGGAATGCTCGTCAAGAAGACAGGGCCAGGTTT
CCGGGCCCTCACATTGCCAAAAGACGGCAATATGGTGGAAAATAACATATAGACAAACGCACACCGGCC
TTATTCCAAGCGGCTTCGGCCAGTAACGTTAGGGGGGGGGGAGGGGGGGGGGGAGAGGGGCGGAAT
TCCTCTAGTGCGGCCGCGGATCCTTACTTAGTTACCCGGGGAGCATGTCAAGGTCAAAATCGTCAAGAGC
GTCAGCAGGCAGCATATCAAGGTCAAAGTCGTCAAGGGCATCGGCTGGGAGCATGTCTAAGTCAAAATC
GTCAAGGGCGTCGGTCGGCCCGCCGCTTTCGCACTTTAGCTGTTTCTCCAGGCCACATATGATTAGTTCCA
GGCCGAAAAGGAAGGCAGGTTCGGCTCCCTGCCGGTCGAACAGCTCAATTGCTTGTCTCAGAAGTGGG
GGCATAGAATCGGTGGTAGGTGTCTCTCTTTCCTCTTTTGCTACTTGATGCTCCTGTTCCTCCAATACGCAG
CCCAGTGTAAAGTGGCCCACGGCGGACAGAGCGTACAGTGCGTTCTCCAGGGAGAAGCCTTGCTGACAC

AGGAACGCGAGCTGATTTTCCAGGGTTTCGTACTGTTTCTCTGTTGGGCGGGTGCCGAGATGCACTTTAG
CCCCGTCGCGATGTGAGAGGAGAGCACAGCGGTATGACTTGGCGTTGTTCCGCAGAAAGTCTTGCCATG
ACTCGCCTTCCAGGGGGCAGAAGTGGGTATGATGCCTGTCCAGCATCTCGATTGGCAGGGCATCGAGCA
GGGCCCGCTTGTTCTTCACGTGCCAGTACAGGGTAGGCTGCTCAACTCCCAGCTTTTGAGCGAGTTTCCT
TGTCGTCAGGCCTTCGATACCGACACCATTGAGTAATTCCAGAGCTCCGTTTATGACTTTGCTCTTGTCCAG
TCTAGACATGGTGAATTCGACTGCAGGACCGGTACGGGGTGGAGATCCGAGCTCGGTACCAAGCTTCGTC
TAACAAAAAAGCCAAAAACGGCCAGAATTTAGCGGACAATTTACTAGTCTAACACTGAAAATTACATATT
GACCCAAATGATTACATTTCAAAAGGTGCCTAAAAAACTTCACAAAACACACTCGCCAACCCCGAGCGC
ATAGTTCAAAACCGGAGCTTCAGCTACTTAAGAAGATAGGTACATAAAACCGACCAAAGAAACTGACGC
CTCACTTATCCCTCCCCTCACCAGAGGTCCGGCGCCTGTCGATTCAGGAGAGCCTACCCTAGGCCCGAAC
CCTGCGTCCTGCGACGGAGAAAAGCCTACCGCACACCTACCGGCAGGTGGCCCCACCCTGCATTATAAG
CCAACAGAACGGGTGACGTCACGACACGACGAGGGCGCGCTCCCAAAGGTACGGGTGCACTGCCCA
ACGGCACCGCCATAACTGCCGCCCCCGCAACAGACGACAAACCGAGTTCTCCAGTCAGTGACAAACTTC
ACGTCAGGGTCCCCAGATGGTGCCCCAGCCCATCTCACCCGAATAAGAGCTTTCCCGCATTAGCGAAGGC
CTCAAGACCTTGGGTTCTTGCCGCCCACCATGCCCCCCACCTTGTTTCAACGACCTCACAGCCCGCCTCAC
AAGCGTCTTCCATTCAAGACTCGGGAACAGCCGCCATTTTGCTGCGCTCCCCCCAACCCCCAGTTCAGGG
CAACCTTGCTCGCGGACCCAGACTACAGCCCTTGGCGGTCTCTCCACACGCTTCCGTCCCACCGAGCGGC
CCGGCGGCCACGAAAGCCCCGGCCAGCCCAGCAGCCCGCTACTCACCAAGTGACGATCACAGCGATCC
ACAAACAAGAACCGCGACCCAAATCCCGGCTGCGACGGAACTAGCTGTGCCACACCCGGCGCGTCCTTA
TATAATCATCGGCGTTCACCGCCCCACGGAGATCCCTCCGCAGAATCGCCGAGAAGGGACTACTTTTCCTC
GCCTGTTCCGCTCTCTGGAAAGAAAACCAGTGCCCTAGAGTCACCCAAGTCCCGTCCTAAAATGTCCTTCT
GCTGATACTGGGGTTCTAAGGCCGAGTCTTATGAGCAGCGGGCCGCTGTCCTGAGCGTCCGGGCGGAAG
GATCAGGACGCTCGCTGCGCCCTTCGTCTGACGTGGCAGCGCTCGCCGTGAGGAGGGGGGCGCCCGCG
GGAGGCGCCAAAACCCGGCGCGGAGGCCAGATCTTGGGTGGGTTACTCCAGACTGCCTTGGGAAAAGC
GCCTCCCCTACCCGGTAGAATTTCTAGTTTAATTAATCATTACTAAGCGTAGTCTGGGACGTCGTATGGGTAT
TCGAACCGCGGGCCCTCTAGACTCGAGCGGCCGCCACTGTGCTGGATATCAACCACTTTGTACAAGAAAG
CTGGGTCTAGATATCTCGAGGCGGCCGCTTATCCGGATTCGAATCATTAGGATCCAGCGTAATCTGGAACG
TCATAAGGATACGATCCTGCATAGTCCGGGACGTCATAGGGATAGCCCGCATAGTCAGGAACATCGTATGG
GTACCCGCCGGTGCCGTCCAGGATGGTCTTGAAGTCTGAGGGCAGTGCCGGGTTGGCTGCGGCCTCCAG

GGCAGTCAGCGTCGTCCCCGGGAGCTTCCGACTCAGCTGCGTCTGGGCTGTCCTGAGTGACCCCAGGAG
TGGCACGTAGGTGACACGGTGTCGAGTCAGCTTGAGCAGGAATGCTTGGTGGCACAGCCACTGCACGGC
CTCGGAGGGCAGAGGGCCGGCGGCGCCCTTGGCCCCCAGCGACATCCCTGCGTTCTTGGCTTTCAGGAT
GGAGTAGCAGAGGGAGGCCGTGTCAGAGATGACGCGCAGGAAAAATGTGGGGTTCTTCCAAACTTGCT
GATGAAATGGGAGCTGCAGCACACATGCGTGAAACCTGTACGCCTGCAGCAGGAGGATCTTGTAGATGT
TGGTGCACACCGTCTGGAGGCTGTTCACCTGCAAATCCAGAAACAGGCTGTGACACTTCAGCCGCAAGA
CCCCAAAGAGTTTGCGACGCATGTTCCTCCCAGCCTTGAAGCCGCGGTTGAAGGTGAGACTGGCTCTGAT
GGAGGTCCGGGCATAGCTGGAGTAGTCGCTCTGCACCTCCAGGGTCCGGGTATCCAGCAGCAGGCCGCA
CCAGGGGAATAGGCCGTGGGCCGGCATCTGAACAAAAGCCGTGCCACCCAGGGCCTCGTCTTCTACAGG
GAAGTTCACCACTGTCTTCCGCAAGTTCACCACGCAGCCATACTCAGGGACACCTCGGACCAGGGTCCTG
AGGAAGGTTTTCGCGTGGGTGAGGTGAGGTGTCACCAACAAGAAATCATCCACCAAACGCAGGAGCAG
CCCGTCCCGCCGAATCCCCGCAAACAGCTTGTTCTCCATGTCGCCGTAGCACAGGCTGCAGAGCAGCGTG
GAGAGGATGGAGCCCTGCGGGATCCCCTGGCACTGGACGTAGGACTTGCCCCTGATGCGCACGGCGTGG
TGGCACATGAAGCGTAGGAAGACGTCGAAGAGGCCACTGCTGGCCTCATTCAGGGAGGAGCTCTGCTCG
ATGACGACGGCATCCCTCAGCGGGCTGGTCTCCTGCAGGTGAGCCACGAACTGTCGCATGTACGGCTGG
AGGTCTGTCAAGGTAGAGACGTGGCTCTTGAAGGCCTTGCGGACGTGCCCATGGGCGGCCTTCTGGACC
ACGGCATACCGACGCACGCAGTACGTGTTCTGGGGTTTGATGATGCTGGCGATGACCTCCGTGAGCCTGT
CCTGGGGGATGGTGTCGTACGCGCCCGTCACATCCACCTTGACAAAGTACAGCTCAGGCGGCGGGTCCT
GGGCCCGCACACGCAGCACGAAGGTGCGCCAGGCCCTGTGGATATCGTCCAGGCCCAGCACAGAGGCG
CCCAGGAGGCCGGGGCGCCGCGCCCGCTCGTAGTTGAGCACGCTGAACAGTGCCTTCACCCTCGAGGT
GAGACGCTCGGCCCTCTTTTCTCTGCGGAACGTTCTGGCTCCCACGACGTAGTCCATGTTCACAATCGGCC
GCAGCCCGTCAGGCTTGGGGATGAAGCGGAGTCTGGACGTCAGCAGGGCGGGCCTGGCTTCCCGATGC
TGCCTGACCTCTGCTTCCGACAGCTCCCGCAGCTGCACCCTCTTCAAGTGCTGTCTGATTCCAATGCTTTG
CAACTTGCTCCAGACACTCTTCCGGTAGAAAAAGAGCCTGTTCTTTTGAAACGTGGTCTCCGTGACATAAA
AGAAAGACCTGAGCAGCTCGACGACGTACACACTCATCAGCCAGTGCAGGAACTTGGCCAGGATCTCCT
CACGCAGACGGTGCTCTGCGGCCGGAACACAGCCAACCCCTGGGCTCCTGCGCAGCCAAGCGCAGTCC
CGCACGCTCATCTTCCACGTCAGCTCCTGCAGCGAGAGCTTGGCATGCTTCCCCAGGGAGATGAACTTCT
TGGTGTTCCTGAGGAAGCGGCGTTCGTTGTGCCTGGAGCCCCAGAGGCCTGGGGGCACCAGCCGGCGC
AGGCAGGCCCGCACGAAGCCGTACACCTGCCAGGGGCTGCTGTGCTGGCGGAGCAGCTGCACCAGGC

GACGGGGGTCTGTGTCCTCCTCCTCGGGGGCCGCCACAGAGCCCTGGGGCTTCTCCCGGGCACAGACAC
CGGCTGCTGGGGTGACCGCAGCTCGCAGCGGGCAGTGCGTCTTGAGGAGCACCCCGTAGGGGCACTGC
GCGTGGTTCCCAAGCAGCTCCAGAAACAGGGGCCGCATTTGCCAGTAGCGCTGGGGCAGGCGGGGCA
ACCTGCGGGGAGTCCCTGGCATCCAGGGCCTGGAACCCAGAAAGATGGTCTCCACGAGCCTCCGAGCGC
CAGTCAGGCTGGGCCTCAGAGAGCTGAGTAGGAAGGAGGGCCGCAGCTGCTCCTTGTCGCCTGAGGAG
TAGAGGAAGTGCTTGGTCTCGGCGTACACCGGGGGACAAGGCGTGTCCCAGGGACGTGGTGGCCGCGA
TGTGGATGGGGGGCCCGCGTGGTGCTGGCGGCCCACGGATGGGTGGGAGTGGCGCGTGCCAGAGAGC
GCACCCTCCAAAGAGGTGGCTTCTTCGGCGGGTCTGGCAGGTGACACCACACAGAAACCACGGTCACTC
GGTCCACGCGTCCTGCCCGGGTGGGCCCAGGACCCCTGCCCAACGGGCGTCCGCTCCGGCTCAGGGGC
AGCGCCACGCCTGGGCCTCTTGGGCAACGGCAGACTTCGGCTGGCACTGCCCCCGCGCCTCCTCGCACC
CGGGGCTGGCAGGCCCAGGGGGACCCCGGCCTCCCTGACGCTATGGTTCCAGGCCCGTTCGCATCCCAG
ACGCCTTCGGGGTCCACTAGCGTGTGGCGGGGGCCGGGCCTGAGTGGCAGCGCCGAGCTGGTACAGCG
GCGGCCCGCACACCTGGTAGGCGCAGCTGGGAGCCACCAGCACAAAGAGCGCGCAGCGTGCCAGCAG
GTGAACCAGCACGTCGTCGCCCACGCGGCGCAGCAGCAGCCCCCACGCCCCGCTCCCCCGCAGTGCGTC
GGTCACCGTGTTGGGCAGGTAGCTGCGCACGCTGGTGGTGAAGGCCTCGGGGGGGCCCCCGCGGGCCC
CGTCCAGCAGCGCGAAGCCGAAGGCCAGCACGTTCTTCGCGCCGCGCTCGCACAGCCTCTGCAGCACTC
GGGCCACCAGCTCCTTCAGGCAGGACACCTGGCGGAAGGAGGGGGCGGCGGGGGGCGGCCGTGCGT
CCCAGGGCACGCACACCAGGCACTGGGCCACCAGCGCGCGGAAAGCCGCCGGGTCCCCGCGCTGCAC
CAGCCGCCAGCCCTGGGGCCCCAGGCGCCGCACGAACGTGGCCAGCGGCAGCACCTCGCGGTAGTGG
CTGCGCAGCAGGGAGCGCACGGCTCGGCAGCGGGGAGCGCGCGGCATGGTGGCGCCAGCCTGCTTTT
TTGTACAAACTTGTTGATATCTGCAGAATTCCACCACACTGGACTAGTTCGGGGCCGCGGAGGCTGGATC
GGTCCCGGTGTCTTCTATGGAGGTCAAAACAGCGTGGATGGCGTCTCCAGGCGATCTGACGGTTCACTAA
ACGAGCTCTGCTTATATAGGCCTCCCACCGTACACGCCTACCTCGACCCGGGTACCGAGCTCGACTTTCAC
TTTTCTCTATCACTGATAGGGAGTGGTAAACTCGACTTTCACTTTTCTCTATCACTGATAGGGAGTGGTAAA
CTCGACTTTCACTTTTCTCTATCACTGATAGGGAGTGGTAAACTCGACTTTCACTTTTCTCTATCACTGATAG
GGAGTGGTAAACTCGACTTTCACTTTTCTCTATCACTGATAGGGAGTGGTAAACTCGACTTTCACTTTTCTC
TATCACTGATAGGGAGTGGTAAACTCGACTTTCACTTTTCTCTATCACTGATAGGGAGTGGTAAAGGATCCT
AGTCCAAACTGGATCTCTGCTGTCCCTGTAATAAACCCGAAAATTTTGAATTTTTGTAATTTGTTTTTGTAAT
TCTTTAGTTTGTATGTCTGTTGCTATTATGTCTACTATTCTTTCCCCTGCACTGTACCCCCCAATCCCCCCTTTT

CTTTTAAAATTGTGGATGAATACTGCCATTTGTGAATTCGGCGATACCGTCGAGATCCGTTCACTAATCGAA
TGGATCTGTCTCTGTCTCTCTCTCCACCTTCTTCTTCTATTCCTTCGGGCCTGTCGGGTCCCCTCGGGGTTGG
GAGGTGGGTCTGAAACGATAATGGTGAATATCCCTGCCTAACTCTATTCACTATAGAAAGTACAGCAAAAA
CTATTCTTAAACCTACCAAGCCTCCTACTATCATTATGAATAATTTTATATACCACAGCCAATTTGTTATGTTA
AACCAATTCCACAAACTTGCCCATTTATCTAATTCCAATAATTCTTGTTCATTCTTTTCTTGCTGGTTTTGCG
ATTCTTCAATTAAGGAGTGTATTAAGCTTGTGTAATTGTTAATTTCTCTGTCCCACTCCATCCAGGTCGTGTG
ATTCCAAATCTGTTCCAGAGATTTATTACTCCAACTAGCATTCCAAGGCACAGCAGTGGTGCAAATGAGTT
TTCCAGAGCAACCCCAAATCCCCAGGAGCTGTTGATCCTTTAGGTATCTTTCCACAGCCAGGATTCTTGCC
TGGAGCTGCTTGATGCCCCAGACTGTGAGTTGCAACAGATGCTGTTGCGCCTCAATAGCCCTCAGCAAAT
TGTTCTGCTGCTGCACTATACCAGACAATAATTGTCTGGCCTGTACCGTCAGCGTCATTGACGCTGCGCCCA
TAGTGCTTCCTGCTGCTCCCAAGAACCCAAGGAACAAAGCTCCTATTCCCACTGCTCTTTTTTCTCTCTGCA
CCACTCTTCTCTTTGCCTTGGTGGGTGCTACTCCTAATGGTTCAATTTTTACTACTTTATATTTATATAATTCA
CTTCTCCAATTGTCCCTCATATCTCCTCCTCCAGGTCTGAAGATCAGCGGCCGGCCGCTTGCTGTGCGGTG
GTCTTACTTTTGTTTTGCTCTTCCTCTATCTTGTCTAAAGCTTCCTTGGTGTCTTTTATCTCTATCCTTTGATGC
ACACAATAGAGGGTTGCTACTGTATTATATAATGATCTAAGTTCTTCTGATCCTGTCTGAAGGGATGGTTGTA
GCTGTCCCAGTATTTGTCTACAGCCTTCTGATGTTTCTAACAGGCCAGGATTAACTGCGAATCGTTCTAGCT
CCCTGCTTGCCCATACTATATGTTTTAATTTATATTTTTTCTTTCCCCCTGGCCTTAACCGAATTTTTTCCCATC
GCGATCTAATTCTCCCCCGCTTAATACTGACGCTCTCGCACCCATCTCTCTCCTTCTAGCCTCCGCTAGTCAA
AATTTTTGGCGTACTCACCAGTCGCCGCCCCTCGCCTCTTGCCGTGCGCGCTTCAGCAAGCCGAGTCCTG
CGTCGAGAGAGCTCCTCTGGTTTCCCTTTCGCTTTCAAGTCCCTGTTCGGGCGCCACTGCTAGAGATTTTC
CACACTGACTAAAAGGGTCTGAGGGATCTCTAGTTACCAGAGTCACACAACAGACGGGCACACACTACT
TGAAGCACTCAAGGCAAGCTTTATTGAGGCTTAAGCAGTGGGTTCCCTAGTTAGCCAGAGAGCTCCCAG
GCTCAGATCTGGTCTAACCAGAGAGACCCAGTACAGGCAAAAAGCAGCTGCTTATATGCAGGATCTGAG
GGCTCGCCACTCCCCAGTCCCGCCCAGGCCACGCCTCCCTGGAAAGTCCCCAGCGGAAAGTCCCTTGTA
GCAAGCTCGATATCAGCAGTTCTTGAAGTACTCCGGATGCAGCTCTCGGGCCACGTGATGAAATGCTAGG
CGGCTGTCAAACCTCCACTCTAACACTTCTCTCTCCGGGTCATCCATCCCATGCAGGCTCACAGGGTGTAA
CAAGCTGGTGTTCTCTCCTTTATTGGCCTCTTCTACCTTATCTGGCTCAACTGGTACTAGCTTGTAGCACCAT
CCAAAGGTCAGTGGATATCTGACCCCTGGCCCTGGTGTGTAGTTCTGCTAATCAGGGAAGTAGCCTTGTGT
GTGGTAGATCCACAGATCAAGGATATCTTGTCTTCTTTGGGAGTGAATTAGCCCTTCCAACTACTAAGTTTG

TAGTACATATTTAACAAATACAATTTCTTTAAAATGAAAATAATTCAGAGGAATCACAGGTTTAGAGTAAAT
GAAACCACAGGTAATTGGCAGTGGTAATAGGGTATGGGGTGGGAAGTTTGGGATGATTTTGGTTAGCTTG
AGTTATCCAGTTGATCCAGACATGATAAGATACATTGATGAGTTTGGACAAACCACAACTAGAATGCAGTG
AAAAAAATGCTTTATTTGTGAAATTCGTAATCATGTCATAGCTGTTTCCTGTGTGAAATTGTTATCCGCTCAC
AATTCCACACAACATACGAGCCGGAAGCATAAAGTGTAAAGCCTGGGGTGCCTAATGAGTGAGCTAACTC
ACATTAATTGCGTTGCGCTCACTGCCCGCTTTCCAGTCGGGAAACCTGTCGTGCCACGGGGAGCTTTTTG
CAAAAGCCTAGGCCTCCAAAAAAGCCTCCTCACTACTTCTGGAATAGCTCAGAGGCCGAGGCGGCCTCG
GCCTCTGCATAAATAAAAAAAATTAGTCAGCCATGAGCTTGCTGCATTAATGAATCGGCCAACGCGCGGG
GAGAGGCGGTTTGCGTATTGGGCGCTCTTCCGCTTCCTCGCTCACTGACTCGCTGCGCTCGGTCGTTCGG
CTGCGGCGAGCGGTATCAGCTCACTCAAAGGCGGTAATACGGTTATCCACAGAATCAGGGGATAACGCA
GGAAAGAACATGTGAGCAAAAGGCCAGCAAAAGGCCAGGAACCGTAAAAAGGCCGCGTTGCTGGCGT
TTTTCCATAGGCTCCGCCCCCCTGACGAGCATCACAAAAATCGACGCTCAAGTCAGAGGTGGCGAAACCC
GACAGGACTATAAAGATACCAGGCGTTTCCCCCTGGAAGCTCCCTCGTGCGCTCTCCTGTTCCGACCCTGC
CGCTTACCGGATACCTGTCCGCCTTTCTCCCTTCGGGAAGCGTGGCGCTTTCTCATAGCTCACGCTGTAGG
TATCTCAGTTCGGTGTAGGTCGTTCGCTCCAAGCTGGGCTGTGTGCACGAACCCCCCGTTCAGCCCGACCG
CTGCGCCTTATCCGGTAACTATCGTCTTGAGTCCAACCCGGTAAGACACGACTTATCGCCACTGGCAGCAG
CCACTGGTAACAGGATTAGCAGAGCGAGGTATGTAGGCGGTGCTACAGAGTTCTTGAAGTGGTGGCCTA
ACTACGGCTACACTAGAAGAACAGTATTTGGTATCTGCGCTCTGCTGAAGCCAGTTACCTTCGGAAAAAG
AGTTGGTAGCTCTTGATCCGGCAAACAAACCACCGCTGGTAGCGGTGGTTTTTTTGTTTGCAAGCAGCAG
ATTACGCGCAGAAAAAAAGGATCTCAAGAAGATCCTTTGATCTTTTCTACGGGGTCTGACGCTCAGTGGA
ACGAAAACTCACGTTAAGGGATTTTGGTCATGAGATTATCAAAAAGGATCTTCACCTAGATCCTTTTAAAT
TAAAAATGAAGTTTTAAATCAATCTAAAGTATATATGAGTAAACTTGGTCTGACAGTTACCAATGCTTAATC
AGTGAGGCACCTATCTCAGCGATCTGTCTATTTCGTTCATCCATAGTTGCCTGACTCCCCGTCGTGTAGATAA
CTACGATACGGGAGGGCTTACCATCTGGCCCCAGTGCTGCAATGATACCGCGAGACCCACGCTCACCGGC
TCCAGATTTATCAGCAATAAACCAGCCAGCCGGAAGGGCCGAGCGCAGAAGTGGTCCTGCAACTTTATC
CGCCTCCATCCAGTCTATTAATTGTTGCCGGGAAGCTAGAGTAAGTAGTTCGCCAGTTAATAGTTTGCGCA
ACGTTGTTGCCATTGCTACAGGCATCGTGGTGTCACGCTCGTCGTTTGGTATGGCTTCATTCAGCTCCGGTT
CCCAACGATCAAGGCGAGTTACATGATCCCCCATGTTGTGCAAAAAAGCGGTTAGCTCCTTCGGTCCTCCG
ATCGTTGTCAGAAGTAAGTTGGCCGCAGTGTTATCACTCATGGTTATGGCAGCACTGCATAATTCTCTTACT

GTCATGCCATCCGTAAGATGCTTTTCTGTGACTGGTGAGTACTCAACCAAGTCATTCTGAGAATAGTGTATG
CGGCGACCGAGTTGCTCTTGCCCGGCGTCAACACGGGATAATACCGCGCCACATAGCAGAACTTTAAAA
GTGCTCATCATTGGAAAACGTTCTTCGGGGCGAAAACTCTCAAGGATCTTACCGCTGTTGAGATCCAGTTC
GATGTAACCCACTCGTGCACCCAACTGATCTTCAGCATCTTTTACTTTCACCAGCGTTTCTGGGTGAGCAA
AAACAGGAAGGCAAAATGCCGCAAAAAAGGGAATAAGGGCGACACGGAAATGTTGAATACT
